# Probing Mechanisms of Allosteric Regulation in AAA+ ATPases for Microtubule Severing and Protein Disaggregation

**DOI:** 10.64898/2026.08.01.742241

**Authors:** Maryum Irshad, Krishan Walpalage, Zhaocheng Zhang, Maria S. Gillen, Ruxandra I. Dima, George Stan

## Abstract

Ring-like AAA+ (ATPases Associated with diverse cellular Activities) biological machines mediate protein remodeling to assist a broad range of essential cellular functions. The nucleotide-dependent remodeling action involves intra- and inter-ring allosteric communication to generate mechanical force applied onto the substrate by a set of loops that protrude into the central channel. In this study, we probe these allosteric mechanisms through a comparative study of the katanin, a microtubule severing protein including a clade 3 AAA domain, and the double-ring ClpB, a protein disaggregase including both a clade 3 and a clade 5 AAA domain. Our molecular dynamics simulations, combined with machine learning and bioinformatic analysis, reveal both similar mechanisms involving the clade 3 domain and ClpB-specific ones involving communication with the clade 5 domain. We find that both nucleotide and substrate polypeptide binding restrict the conformational landscape sampled by katanin and ClpB, with ligand-specific conformations observed in the latter case. Allosteric contributions of secondary structure elements, ranked by using SHapley Additive exPlanations analysis in machine learning approaches and binary classification of features in ligand states, highlight the important role of regions adjacent to the nucleotide-binding site and the pore loops. Amino acid-level analysis of the allosteric paths reveals that intra-ring cooperativity modulates long-distance communication within the AAA+ protomers.

## I. INTRODUCTION

AAA+ (ATPases Associated with diverse cellular Activities) proteins carry out intricate cellular functions ranging from DNA replication to the targeted degradation of proteins, the dynamic fusion of membranes, the precise regulation of microtubule dynamics, and protein disaggregation^1^. Out of 7 clades of AAA+ proteins, distinguished based on variations in the conserved motifs within the AAA+ domain and distinct cellular functions, the best characterized is the classic clade 3^2^. Within this clade distinct subfamilies demonstrate specialized molecular functions, such as FtsH (membrane protein homeostasis), katanin (microtubule severing), Pex1/Pex6 (peroxisome biogenesis), and ClpA/B nucleotide-binding domain 1, or NBD1 (protein degradation/disaggregation)^3,4^. A subgroup of double-ring machines, such as ClpA/B, include a second NBD, which is part of clade 5 (HCLR) along with HslU/ClpX and Lon (protein degradation). Each NBD harbors specialized Walker A (WA) and B (WB) sequence motifs which are responsible for nucleotide binding and hydrolysis. Nucleotide-binding is a stringent requirement for oligomerization into the functional hexameric structure, which, remarkably, adopts a non-planar organization, as revealed by cryo-electron microscopy (cryo-EM) ^5–8^. The hallmark of the AAA+ machine action, conserved from fungi to humans^9^, is the propagation of the allosteric signal generated by ATP hydrolysis, through coordinated conformational changes, to the set of flexible loops protruding into the central channel that engage the substrate and impart mechanical force onto it (Fig. 1). Despite the recent intense focus on structural aspects of AAA+ machines, the detailed allosteric communication is still insufficiently understood.

**FIG. 1.**
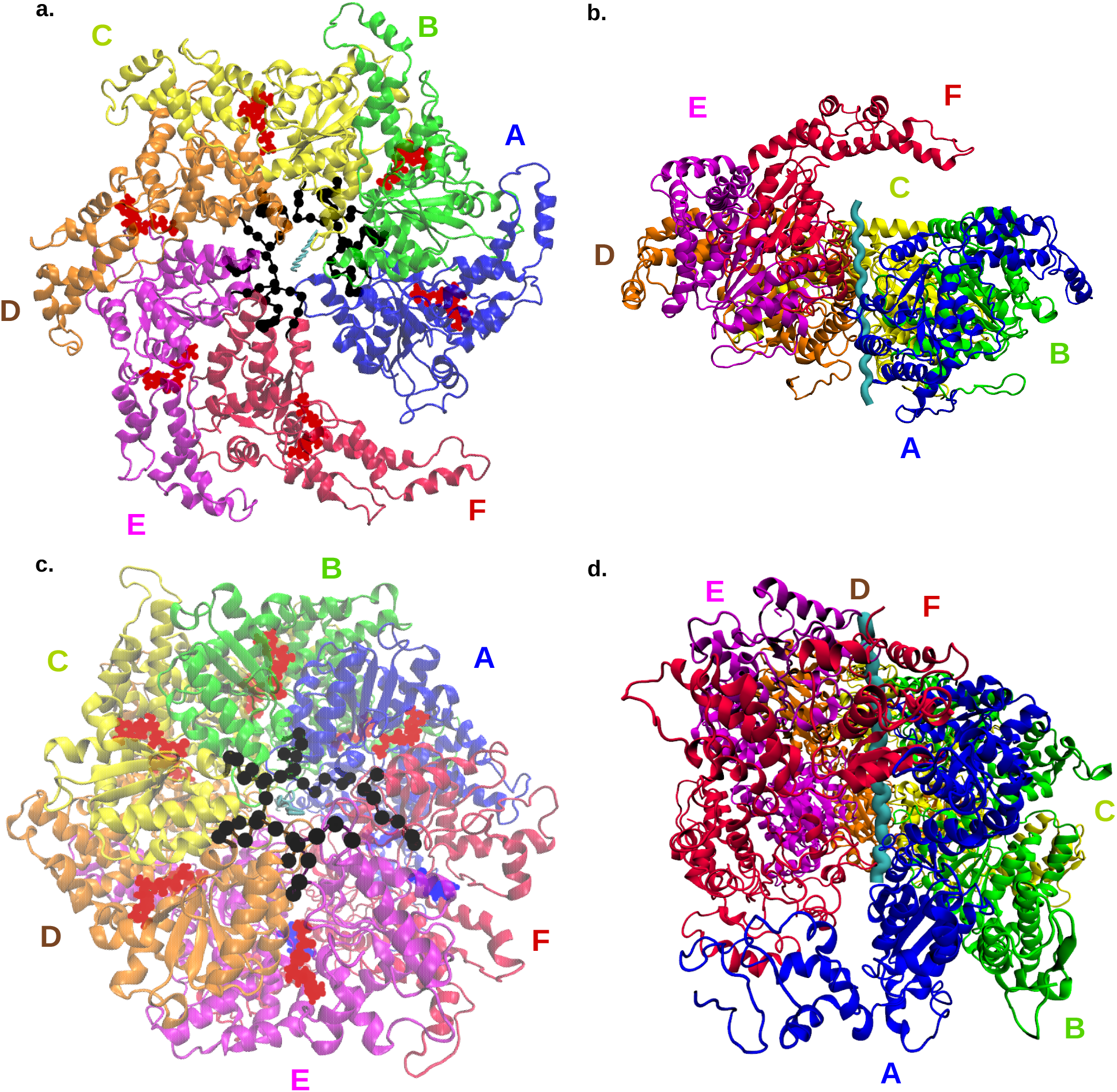
Hexameric structures for (a) top view of spiral conformation of katanin (PDB: 6UGD). Pore loop 1 is shown with black beads. Substrate, E14 is shown in cyan at the central pore, and ATP occupying the nucleotide-binding pocket in all chains is shown in red color. (b) side view of the conformation of katanin showing the right hand spiral around the substrate (shown in cyan) (c) top view of ring conformation of ClpB (PDB: 6OAX). Pore loop 1 is shown with black beads. The substrate, A26, is shown in cyan within the central pore, and AGS (a slowly hydrolyzable ATP analog) and ADP molecules occupying the nucleotide-binding pockets in all chains are shown in red and blue, respectively. (b) side view of the conformation of ClpB showing the right-hand spiral around the substrate (shown in cyan

Katanin, named after the Japanese sword katana, has a catalytic subunit (p60) and a regulatory subunit (p80)^10^. The p60 catalytic subunit in katanin has the AAA+ ATPase motor and is responsible for the severing activity of microtubules independent of the regulatory subunit, whereas the regulatory p80 subunit manages the association with the centrosome and improves microtubule binding. Pore loops PL1, PL2, and PL3 are responsible for the binding of the C-terminal tails (CTTs) of the tubulin monomers. The motor domain also has a helical bundle domain (HBD) comprising four alpha helices, which is unique to the severing enzymes. In lower-order oligomers, a hot spot analysis indicated that the HBD tip was a significant allosteric region^11^. In the ”spiral” (pre-hydrolysis) conformation, the protomers are arranged in a right-handed spiral with each protomer having a 5°A rise, and with a 40°A gap between the boundary protomers (Fig. 1(a, b)). Upon hydrolysis, the protomer A was found to lack the nucleotide and become flexible, thus closing the gap between the boundary protomers and resulting in a ”ring” conformation^6,12^.

ClpB, a member of the Clp (Caseinolytic peptidase) family, plays a crucial role in bacterial survival under thermal stress by solubilizing and unfolding toxic protein aggregates^13^. It collaborates with the DnaK/DnaJ chaperone system to rescue aggregated proteins, promote disaggregation, and restore their cellular function^13^. Unlike other Clp proteins, ClpB lacks an associated peptidase compartment and operates independently to remodel substrates (Fig. 1(c,d)). ClpB is a class I AAA+ ATPase that comprises an N-terminal domain (NTD), two AAA+ nucleotide-binding domains (NBD1 and NBD2), and a middle domain, connected by flexible linkers. The NTD is involved in substrate recognition, while the middle domain regulates activity and mediates inter-protomer communication^14^. NBD1 and NBD2 is further subdivided into a large (L) and small (S) subdomains that coordinate ATP-driven conformational changes and inter-domain communication^15^. Two sets of flexible pore loops are found in each NBD. PL1 in NBD1 is critical for substrate recognition and stabilization, while PL3 in NBD2 drives substrate translocation, with conserved tyrosine residues playing a key role in gripping substrates. Noncanonical pore loops, PL2 in NBD1 and PL4 in NBD2, assist PL1 and PL3 and by enhancing substrate engagement. Transitions between the spiral (pre-hydrolysis) and ring (post-hydrolysis) states of ClpB^8^ provide the essential power stroke that mediates substrate unfolding and threading through the central pore, coordinated by allosteric coupling between the nucleotide-binding sites and flexible pore loops. Notably, optical tweezers and single-molecule FRET experiments highlighted rapid translocation of unfolded substrates on sub-ms timescales^16,17^. Differential response of PL1 compared with PL2 and PL3 to mutations that abolish ATP hydrolysis supports a specialized role of the two NBDs^18,19^.

Experimental insights have provided a picture of how katanin’s mechanochemical cycle is organized through asymmetric inter-protomer interactions and allosteric coupling between functional sites^6,12^. Molecular dynamics (MD) simulations and cryo-EM analyses revealed that the spiral to ring transition is not a single rigid rearrangement but a nucleotide dependent power stroke driven by one or two gating protomers that coordinate ATP hydrolysis^11,12,20^. By contrast, research has revealed that ClpB’s allosteric control is achieved through coordinated inter-domain and inter-protomer communication. Despite sharing a conserved AAA+ domain, katanin and ClpB have distinct quaternary architectures to perform their specialized mechanical roles. Remarkably, their remodeling action is performed on substrates that range from microtubules at the *µ*m length scale, in the case of katanin, to protein aggregates at the nm scale, in the case of ClpB^21^. Currently, we lack a framework that incorporates these protein-specific adaptations into the canonical AAA+ network of WA and WB motifs, arginine fingers and flexible pore loops. Katanin leverages the specific arrangement of AAA+ domains as well as the flexible loops at its central pore for its function whereas in ClpB the interdomain communication (NBD1-NBD2) modulates the ATPase activity and hence translocating the substrate. Yet the molecular pathway by which these domains interact with the AAA+ network remains undefined. Our goal in this work is to map and compare allosteric signal transmission pathways in katanin and ClpB, thereby highlighting their shared mechanistic behavior as well as protein-specific functions. To achieve our goal, we rely on data from long MD simulations and employ a combined approach established in our recent work focused on the allosteric communication in spastin, the other major microtubule severing protein, and the changes induced by a major disease-related mutation in the transmission of these allosteric signals^22,23^. We started from the determination of the metastable states from the MD simulations of monomers to establish the major states and the transitions between them that characterize the binding of ligands to the monomeric forms of katanin and ClpB. Next we employed machine learning approaches to extract the secondary structure regions from both monomers and hexamers that control the response of these AAA+ machines to perturbations. Based on our previous spastin work^22,23^, we posit that the resulting regions are the major allosteric elements of katanin and ClpB. Another approach, crucial for elucidating allosteric mechanisms, is the mapping of the allosteric community network that has been used for the identification of conformationally important residues, including remote positions coupled to the active site, and had successfully rationalized experimental findings^22,24–27^. Using the community network analysis, we determined the most likely pathways of allosteric communication in our AAA+ systems as well as the type of wiring that enables each of them to perform their cellular functions.

## II. METHODS

### A. Molecular Dynamics Simulations

In our simulations, initial configurations of the AAA+ hexamers were prepared using high-resolution cryo-EM structures of the spiral conformation of C. elegans katanin (Protein Data Bank ID: 6UGD) and ring conformation for E. coli ClpB (6OAX), both representing pre-hydrolysis states, were obtained in the presence of nucleotides (ATP in katanin/ AGS and ADP combination in ClpB) and minimal substrates (14-residue polyglutamate (E14) for katanin/26-residue poly-alanine (A26) for ClpB)^8,12^. The katanin cryo-EM structure (6UGD) had two regions with missing residues: 183-187, and 324-331, which were modeled using the Modeler program^28^. The cryo-EM structure assigns residues 399 to 436 as a single long helix^12^, however, the per-residue helix probability across residues 417-424 is very low (0.03 in the online entry^29^), increasing to 0.27 only at residue 425. We therefore assigned these as separate secondary structure elements H10, L16, and H11, which shifted the assignment such that the helix H12 in the original structure corresponds to H13 in our work (see Table S1 from the supplementary material). The cryo-EM structure of ClpB (6OAX) lacks coordinates for the N-terminal domain (residues 1–160) and the regulatory M-domain (residues 409–524). To focus on the AAA+ core responsible for substrate translocation, the N-terminal domain was omitted from the simulations. The missing M-domain was replaced with a five-glycine linker, which preserves the continuity of the polypeptide chain while minimizing structural restraints between NBD1 and NBD2. We also performed simulations of the monomeric form of katanin and ClpB using protomer A (residues 156–472 for katanin and residues 161–853 for ClpB) of the corresponding hexameric conformation^11^. The sequence ranges for the secondary structures in each system are listed in the supplementary material Tables S1 for ClpB and S2 for katanin. For both the hexamer and monomer systems, we considered 4 setups: (i) CPX; both nucleotides and substrate present (ii) SUB; only substrate (iii) ATP; only nucleotides (iv) APO; no ligands present - for a total of 16 configurations^22^. For each setup of monomeric and hexameric systems, we performed all-atom molecular dynamics simulations using GROMACS (v 2020)^30^ and the GROMOS 54a7 force field^31^. The automated topology builder server was used to generate force field parameters for ATP^32^. Each system was placed in the center of an orthogonal water box, solvated with the SPC solvent model, and neutralized with NaCl ions^33^. Periodic boundary conditions (PBC) were applied in three-dimensions. The systems were minimized using the steepest descent algorithm and the Verlet cutoff-scheme for 50,000 steps with a criteria of the maximum force value smaller than 23.9006 kcal/mol/°A (1,000 kJ/mol/nm) to remove steric clashes^34^. The systems were then equilibrated for 500 ps in the NVT ensemble to bring the temperature to 300 K by using the velocity-rescaling thermostat and the leap-frog integrator algorithm^35^. Next, a second equilibration step was performed for 500 ps in the NPT ensemble to maintain the system at a pressure of 1.0 bar by using the Parrinello-Rahman pressure coupling scheme^36^, again with the leapfrog integrator algorithm. For katanin, we performed five production trajectories for both the monomer (for a minimum of 70 ns) and the hexamer (for a minimum of 200 ns). For ClpB, we performed three production trajectories for the monomer (for a minimum of 150 ns) and five for the hexamer (for a minimum of 150 ns).

#### 1. Feature-Based Simulation Convergence

To check the convergence of our simulations, we monitored the backbone RMSD along with a feature-based analysis using the solvent-accessible surface area (SASA), as described below, to evaluate when the system had sampled a sufficient amount of the conformational space^37^. We found that the SASA of each system reliably differentiates the 4 ligand states in katanin and ClpB, making it a robust indicator of dynamic changes throughout our simulations^22^. SASA was calculated using GROMACS^30,38^. We studied the distribution of SASA over 25 ns intervals until a non-significant *p*-value (*>* 0.001) was achieved. Specifically, a two-sample Kolmogorov-Smirnov test (ks 2samp from Python’s SciPy v1.12.0) demonstrated increasing similarity between consecutive final time intervals^39^. For katanin, convergence was achieved at 200 ns for the ATP-bound hexamer setups (ATP and CPX), at 300 ns for the APO setup, and at 450 ns for the SUB setup. For the monomer, the CPX setup converged at 70 ns, while the remaining setups converged at 100 ns. For ClpB, both the hexamer and monomer configurations demonstrated convergence at 150 ns. The convergence results for each of the 16 configurations are provided in Figures S2 and S3 from the supplementary material.

### B. Machine Learning

To identify allosteric changes due to ligand binding, we employed our previous methodology that successfully highlighted experimentally known allosteric regions in spastin^22^. Following our previous paper, the XGBoost classifier was used^22^. We then extracted RSA, COULOMB and VDW descriptors from each of our simulations. These descriptors, shown in Table 1, were selected because their distributions exhibited observable shifts between the different ligand states. We then employed the machine learning classifier XGBoost along with the explanatory algorithm SHAP (SHapley Additive exPlanations) to determine the most significant differences between two protein-ligand states (Figure S7-S8)^40,41^. RSA, COULOMB and VDW descriptors effectively captured the structural rearrangements, electrostatic and non-electrostatic/dispersive environment changes upon allosteric perturbations. Our predictive models analyze each protein-ligand state pair of one protein for one descriptor at a time. Each feature in the model represents a descriptor listed in Table 1 for a secondary-structure element in the protein across the sampled trajectories (Tables S1-S2). The simulations were sampled every 200 ps, resulting in approximately 10,000 frames per transition. Monomer transitions all start from the APO state and transition to binding one ligand (ATP or SUB) or both ligands (CPX). On the other hand, since the hexameric state of the protein forms in the presence of the ligands, the hexamer transitions start from the CPX state and transition to states characterized by the loss of one or both ligands (APO).

**TABLE 1.** Descriptors extracted at the level of secondary structure for katanin and ClpB.

| Descriptor | Description |
| --- | --- |
| RSA | Ratio of solvent accessibility of each residue to the maximum possible accessibility, which was calculated for each amino acid “X” by its SASA measurement within a Gly–X–Gly tripeptide. The ratio per residue is averaged over the secondary structure<sup>42</sup>. |
| COULOMB | Coulombic energy associated with the interaction between a secondary structure and the remainder of the protein (kJ/mol). |
| VDW | van der Waals energy associated with the interaction between a secondary structure and the remainder of the protein (kJ/mol). |

Finally, the descriptor features for secondary structures located within 3°A of the ATP (or substrate) binding site were removed for ligand transitions involving ATP (or the substrate) to avoid biasing the binding pocket in a model^22^.

### C. Markov State Models

Markov State Models (MSMs) were constructed from all the monomer states of our protein monomer simulation frames^43,44^ to identify the effect that ligand binding has on protein conformation. Here, we used the Pyemma (v 2.5.12) python package to build the MSM with a feature subspace from SI Table of features^45^. We selected the feature subspace by ranking their ability to capture the most kinetic variance from the original simulation data after reducing the dimensionality with the Variational Approach for Markov Processes (VAMP)^46^. We found that COULOMB gave the highest VAMP-2 score for katanin, while RSA gave the highest VAMP-2 score for ClpB. (Figure S4-S5). We employed TICA reduction followed by k-means clustering of 200 cluster centers^47–49^. For both katanin and ClpB we selected 5 macrostates based on implied timescale plots, which depict the slowest dynamic processes and 5 ns lag time was chosen depending on which lag time produced the highest VAMP-2 score (Figure S4-S5). The Chapman-Kolmogorov (CK) test was used to check our final model’s validity (Figures S4-S5). To classify the binding-order preference of each system, we generated bar plots showing the amount of each simulated ligand state that made up each macrostate.

### D. Sequence Alignment

To determine the common and protein-specific allosteric elements of the proteins tested in our study, we built a sequence alignment of katanin and ClpB. Moreover, to compare with our previous work^22^, we included spastin in the alignment. To this end, we used the sequences corresponding to the PDB entries of katanin (6UGD), ClpB (6OAX), and Spastin (6P07) chains A and we employed UniProt’s Align with 5 iterations to build the alignment.

### E. Graph Networks and Path Analysis

Following our previous study^22^, we built the dynamic networks of the monomer and the hexamer using the Python library Network X, such that nodes located at C*_α_* positions represent amino acids. Edges, connecting pairs of residues located within 10°A of each other in the starting structure for the simulation, are weighted by the strength of the dynamic cross-correlation values of residue pairs averaged over the five trajectories of a given ligand state^22,50–52^.

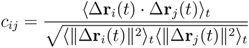

where **r***_i_*(*t*) and **r***_j_*(*t*) are the position vectors of C*_α_* atoms of residues *i* and *j*, ⟨·⟩*_t_* denotes the time ensemble average, and Δ**r***_i_*(*t*) = **r***_i_*(*t*) − ⟨**r***_i_*(*t*)⟩*_t_* and Δ**r***_j_*(*t*) = **r***_j_*(*t*) − ⟨**r***_j_*(*t*)⟩*_t_* are the displacement vectors from their respective mean positions. The dynamic network incorporates the simulations holistically. A *c_ij_* cutoff was chosen by evaluating the ratio, R, of the number of residues in the largest cluster in the network to the total number of residues in the system. Clusters consist of groups of residues connected by edges whose *c_ij_* values satisfy the selected cutoff threshold. Increasing the cutoff progressively removes weaker correlations, causing the network to fragment into smaller clusters. Cutoff values were systematically evaluated over the range 0.4 - 0.8 and the optimum threshold was defined as the highest value at which R reached 0.9, the point beyond which further change in cutoff produced no change in R, which indicates that the network had reached its most stable, meaningful connectivity while minimizing the inclusion of weaker correlations. For katanin, a cutoff of 0.6 was chosen since that allowed an R of over 0.9 for both APO and CPX setups. The cutoff for all states of ClpB is 0.5. We identified the nodes that are mediators of long-range communication in our protein systems as the residues with the top betweenness centrality. This network parameter is defined as the number of shortest paths, built between all pairs of nodes, that pass through a given node^53–57^. (Supplemental Tables S3-S5 for katanin and S15-S17 for ClpB). We carried out an in-depth path analysis on the dynamic networks to describe the flux, or the propagation of allosteric signals within the system. For katanin, to describe the communication within the NBD, we collected paths from the ATP binding pocket (sources: WA - P235 and T240) to the CTT binding channel (sinks: PL1 - W266 and E271, PL2 - H307 and R312). To describe the inter-domain communication, we collected paths from the CTT binding pore (sources: PL1 - W266 and E271, PL2 - H307 and R312) to the HBD tip (sinks: T418 and L426) (previously identified to be a particularly an important hotspot in katanin oligomers)^6,12,22,58^. For ClpB, for intra-NBD communication, we collected paths from the WA motif of NBD1 (WA 1: G206 and T213) to Tyr-containing PL1 (Y251 and R258); and from the WA motif of NBD2 (WA 2: G605 and T612) to another Tyr-containing PL3 (Y653 and G660). To describe inter-domain communication, we collected paths from PL1 to WA2 and from PL3 to WA1. Moreover, we also collected paths from PL1 to R819 on the C-terminal. The R819A mutation was found to inhibit the oligomerization of ClpB.^859^ For each pair of sites, optimal and suboptimal paths were determined using the methods in our previous paper (Tables S6-S14, S18-S26)^22^. The optimal paths, determined with the Dijkstra method in NetworkX as the sum of the weights of the involved edges, were extracted to identify the shortest path for the allosteric signal to propagate^51,52,60^. The suboptimal paths (restricted to a total of 20,000 paths per collection), which are slightly longer than the optimal paths, were calculated using the Yen algorithm^61^, and used to identify residues frequently sampled by allosteric signals. For each collection of paths, we determined the relative degeneracy of each node, defined as the number of suboptimal paths traversing that node divided by the total number of paths in the collection. Next, we retained only the positions with high degeneracy (*>* 0.1) and their values were averaged within each secondary structure element to identify regions that act as key controllers of the communication within the protein (Tables S27-S46).

## III. RESULTS AND DISCUSSION

### A. Metastable States Explain Ligand-Induced Structural Ensembles

We probed the conformational space of katanin and ClpB monomers by building Markov State Models (MSMs) of the configurations sampled in the Molecular Dynamics (MD) simulations. To this end, we considered four setups (see Methods): (i) CPX; both nucleotides and polypeptide substrate present (ii) SUB; only substrate (iii) ATP; only nucleotides (iv) APO; no ligands present^22^. The selection of the MSM states was made based on the descriptors with the highest VAMP-2 score (COULOMB for katanin and RSA for ClpB), as detailed in the Methods and represented in Figures S4 and S5 from the supplementary material. For both katanin and ClpB, our analysis identified 5 conformational ensembles, or metastates, and revealed common and divergent features in the conformational landscapes. Although both proteins exhibit ligand-dependent changes in the metastate populations and transition pathways, they differ in the degree of conformational overlap among ligand states and in the connectivity between metastates (Fig. 2).

**FIG. 2.**
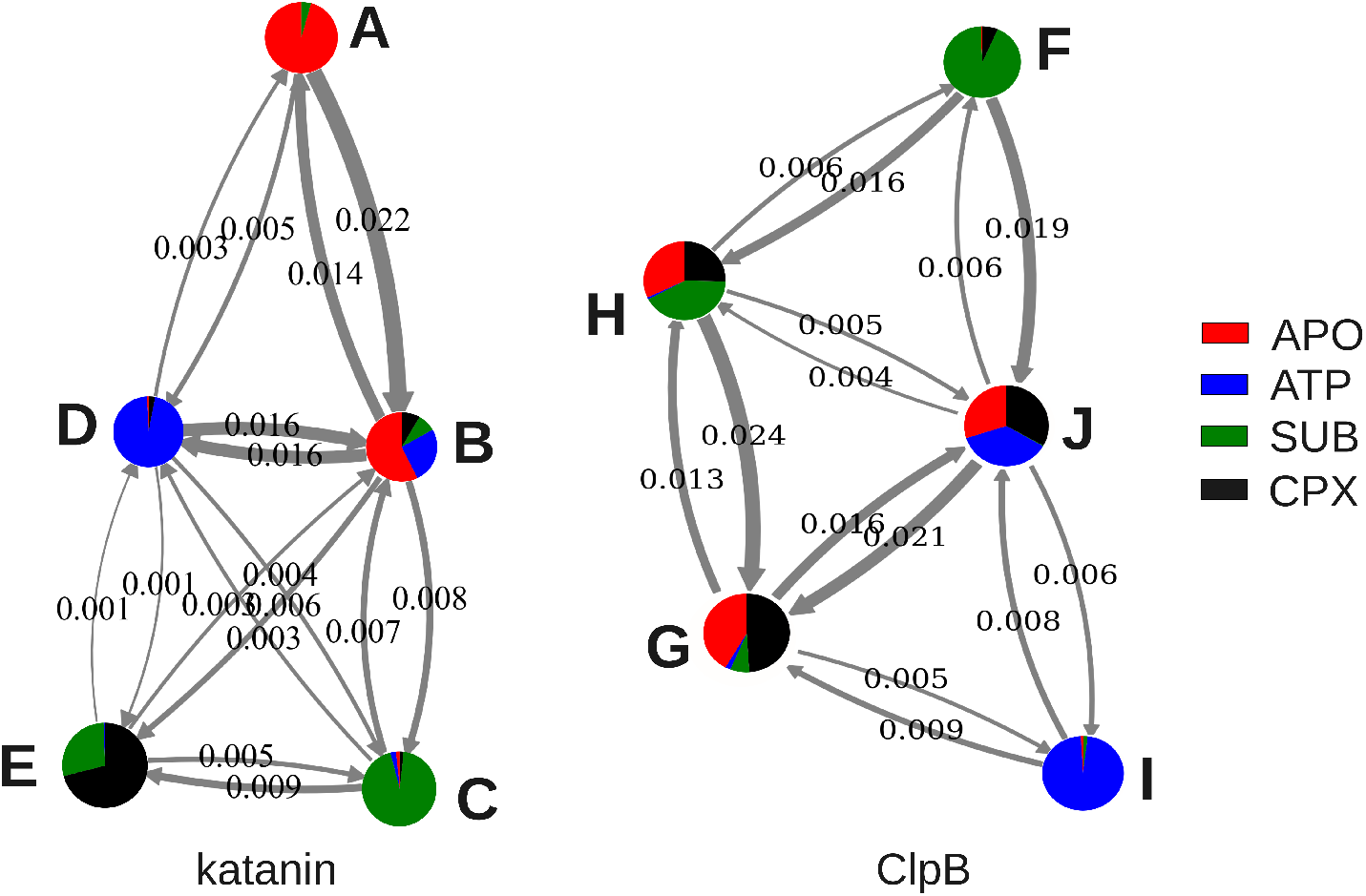
Markov State Models of katanin and ClpB monomers. Each metastable state is labeled with a letter and includes a pie chart of frames sampled in each ligand state (APO, red; ATP, blue; SUB, green; CPX, black). Transition probabilities between each pair of states are indicated by numbers and relative line widths.

As shown in Figure 2, we identified several similarities between the two MSMs, such as the presence of one metastate populated nearly exclusively by frames sampled in a single ligand setup, i.e. ATP (D in katanin and I in ClpB) or SUB (C in katanin and F in ClpB). These results indicate that binding of a single ligand induces a distinct global reduction in the conformational variety of each protein. We also noted that conformations from the APO simulations were split among multiple metastates of each set of MSMs, underscoring the ability of the monomer to broadly sample the conformational space in the absence of ligand-induced constraints. Importantly, we observed that the most likely transition from each of the APO- or SUB-populated metastates is to the ATP-populated metastates. This finding is consistent with the role of the ATP binding in the ClpB structure stabilization, which enables high-affinity substrate binding and subsequent disaggregation activity^62^, and to the ATP-induced stable structure formation in the substrate binding pore loops of katanin priming them for sequential substrate (CTT) binding^12^. We note that in our previous study for spastin, another member of the clade 3 of AAA proteins and a microtubule severing machine, we found that the MSM built for the spastin monomers also supported a preferred route to achieving the CPX setup through the conformational selection resulting from substrate binding to the ATP setup^63^. Thus, we attribute these shared aspects of the MSMs to common trends within the clade 3 of AAA proteins.

We also found specific aspects of the MSMs of katanin and ClpB, which are likely attributes of their sub-families. Namely, katanin’s metastate B contained at least 10% of conformations from every ligand state (Fig. 2), whereas there is no corresponding state in ClpB. This correlates well with the fact that katanin, unlike ClpB, is more stable as a monomer than a hexamer and relies on the combined effect of ATP and substrate binding to form the hexamer^64^, which explains the interchangeable conformations found between any of the ligand states. This is crucial for katanin so that the monomers can accumulate on the microtubule surface before hexamerization is initiated at the spindle poles^11,64^. Another notable difference between the MSMs is that katanin’s ATP and SUB clusters (metastates D and C, respectively) can transition directly between each other while ClpB’s ATP and SUB clusters (metastates I and F, respectively) cannot.

### B. Machine Learning Pinpoints Allosteric Perturbations from Ligand-Binding

Allosteric effects induced by ligand-binding involve complex changes within the monomer and hexamer conformations that preclude a straightforward analysis of individual structural features at the microscopic level. To overcome this challenge, we employed a machine learning model that ranks all regions in each of the two proteins based on the most significant changes between the ligand-bound and the unbound states. This model used the XGBoost decision tree algorithm (see Methods) to classify the secondary structure elements according to the contributions of several biophysical properties that serve as allosteric descriptors. Relative solvent accessibility (RSA), electrostatic (COULOMB) and van der Waals (VDW) interactions were identified as suitable features based on the marked separation of their distributions in distinct protein-ligand states (Supplemental Fig. S1 and Methods). As in our previous studies of ClpP^65^ and spastin^22^, a binary classification approach identifies the top features in the transitions between states, i.e. APO → ATP/SUB/CPX in the monomer, and CPX → ATP/SUB/APO in the hexamer. On the basis of our previous findings for spastin^22^, for which there exists extensive experimental data regarding its allosteric regions^58,66^, we propose that the highest ranked secondary structure elements are the regions responsible for allosteric responses within ClpB and katanin (Figs. 2-3 and Supplemental Figs. S7-S16). The direction and magnitude of the change in the value of each descriptor between two ligand states of each protein can be evaluated by using a SHapley Additive exPlanations (SHAP) analysis of the values of each feature in binary classification models corresponding to two proteins states, e.g. APO and SUB (Supplemental Figs. S7-S8). The results are illustrated through SHAP beeswarm plots, which highlight the ranked-order top features that characterize the transitions between the two states. Points with positive SHAP values have a stronger influence on the predictions made by the XGBoost model in ligand binding, thus playing a larger role in determining the outcome. The raw descriptor value is illustrated through a color gradient that ranges from magenta (highest value) to blue (lowest value). According to the clade membership, we evaluate separately the clade 3 domains, NBD1 in ClpB (secondary structural elements L1-H12, as indicated in SI Table S2) and NBD in katanin (L1-L13, SI Table S1), and the clade 5 domain NDB2 in ClpB, (L13-B17, SI Table S2). We note that the SHAP plots of katanin and ClpB monomers displayed more varied key allosteric regions due to their smaller size compared to hexamers, whereas the SHAP plots for hexamers highlighted secondary structures important across multiple chains.

**FIG. 3.**
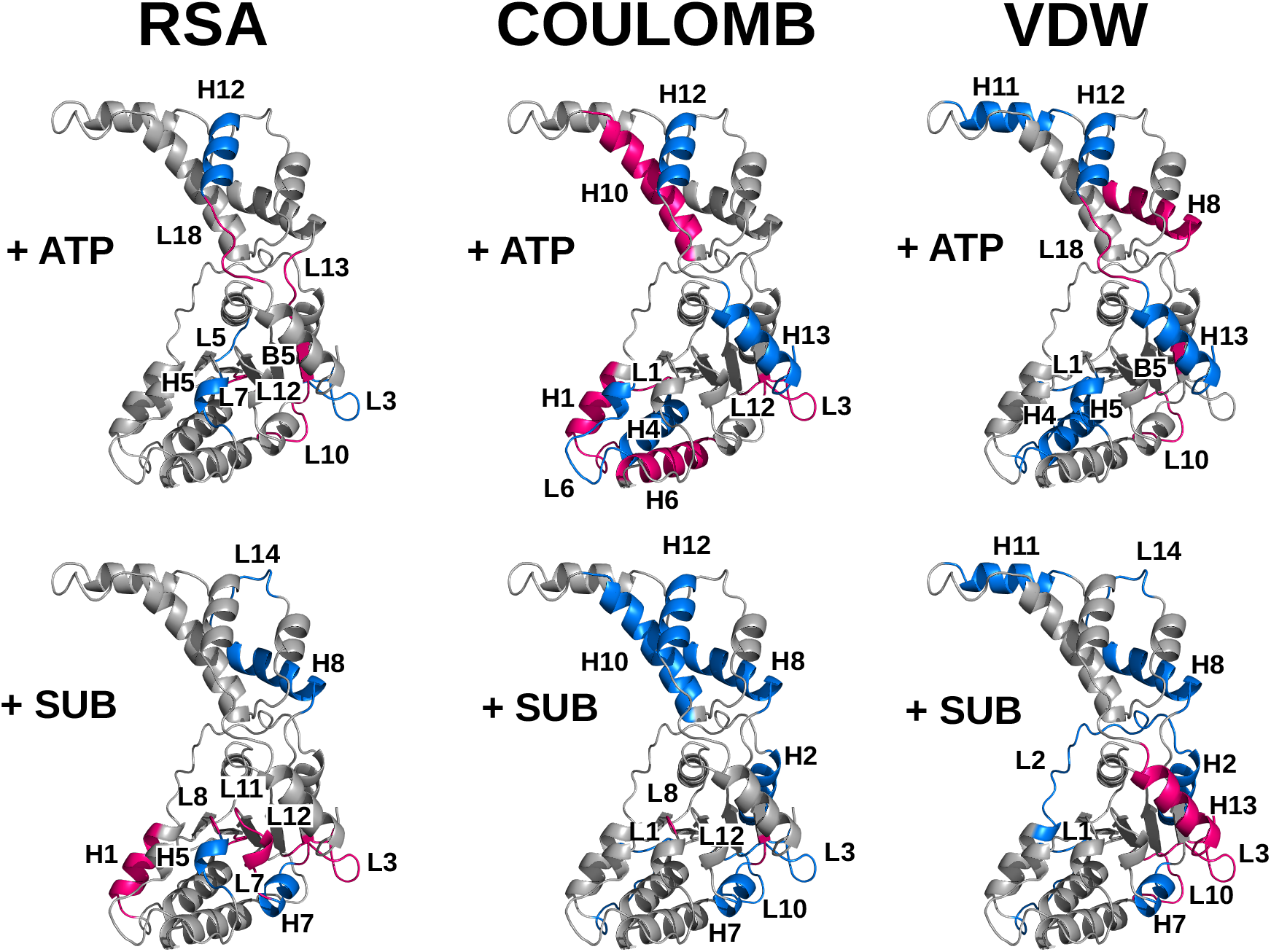
Key allosteric regions of the katanin monomer in ligand-binding transitions. Secondary structure elements of the katanin monomer with the largest contribution in the transition due to nucleotide (ATP, top panels) or substrate (SUB, bottom) binding. Structural regions are color-coded according to the increase (red) or decrease (blue) in RSA (left panels), COULOMB (middle panels), and VDW (right panels) feature values.

Across both the monomer and hexamer for katanin, the secondary structure elements that distinguish ligand states belong predominantly to the NBD rather than the HBD (Figure S7). This result is consistent with the role of the NBD in transducing the allosteric signal from the ATP and substrate binding, and the primary role of the HBD in maintaining the overall shape of the protein. The regions that appeared as top features in both the monomer and hexamer were L1, H1, L2, L3, H5, L10, L11, L12, and B5 from NBD, and L18, and H13 from HBD (Figs. 3, and S12-13, and S15 in the supplementary material). Strikingly, the top features are not catalytic motifs but rather regions involved in the interprotomer communication and hexamer assembly. This strongly suggests that the effects of ATP and substrate binding are not limited to a local perturbation but they include signal propagation directly to the interfaces governing the geometry and stability of the hexamer. In what follows, we review the major functional and structural elements with information about additional regions provided in the Results part 1 of the supplementary material.

For L1, the fishhook linker that is a characteristic of katanin but it is absent in spastin, in the monomer substrate binding led to a decrease in Coulombic energy, while the presence of either ligand alone resulted in a reduction of the van der Waals energy, thus corresponding to an energetic stabilization of L1 in the bound-states (Figs. 3, and S9). In the hexamer, L1 from the internal chains is destabilized in the absence of the substrate alone or of both the ligands (Supplemental Figs. S11, and S12). Instead, L1 is stabilized by the presence of the ATP. This finding aligns well with the fact that in experiments a mutation of the L1 position 170, which is involved in the interaction network between the fishhook and PL1 adjacent elements, from Tyr to Ala reduces ATPase to background levels resulting in a loss of the severing function^12^.

H1, the helix immediately following the fishhook linker and an important oligomerization element, showed a conserved but context-dependent role in the assembly. In the monomer, ATP binding destabilized H1 (Fig. 3). In the hexamer, however, H1 destabilization occurs only upon substrate removal (Supplemental Fig. S11). These findings suggest that ATP engagement in the katanin monomer allosterically triggers a conformational change at the oligomerization interface, promoting the assembly of the hexamer, which is subsequently stabilized by the binding of the substrate.

L11, or the PL3 loop, is involved in the communication between the substrate binding region and the ATP binding region of severing proteins. We found that, in the monomer, substrate binding transiently increased its solvent exposure, while simultaneous nucleotide binding stabilized the local VDW packing (Fig. 3, and S9), suggesting that substrate engagement introduces flexibility that is dampened by ATP. In the hexamer, (Supplemental Fig. S11) internal chains displayed reduced RSA and Coulombic energy and unfavorable VDW interactions when the substrate was absent, reflecting the ligand-dependent constraint of this loop. The terminal chain F showed an elevated sensitivity to the ligands across both RSA and COULOMB, consistent with its end position in the assembly^12,20^.

For H13 (CT-Hlx), which is essential for severing protein oligomerization, in the monomer (Fig. 3), ATP binding stabilized its electrostatic interactions while substrate binding alone introduced local VDW strain, suggesting that the nucleotide binding promotes an interaction-competent conformation whereas substrate engagement introduces local flexibility. In the hexamer, ATP removal resulted in an increase in RSA, indicating a relaxed hexameric arrangement (Supplemental Fig. S15), whereas, the removal of the substrate and ATP led to an increase in the VDW energy, reflecting a loss of favorable interactions and reduced local stabilization in the ligand free state. Our findings suggest that, for the katanin hexamer, the presence of the ATP supports the global compaction of the structure. Moreover, taking into account the behavior of the monomer, our results strongly support the idea that the binding of the substrate is favored only following katanin’s oligomerization as it results in the local stabilization of the hexamer through the formation of stabilizing VDW contacts. In turn, binding of the substrate to the monomer is disfavored as it leads to energetic instability.

L12 (E354, K355), located adjacent to the Arg Finger motifs (R351, R352), showed a distinct behavior between the monomer and the hexamer. In the monomer (Fig. 3, and S7), L12 was more exposed and electrostatically unstable in the presence of ligands. In the hexamer, L12 is located at the inter-protomer interface where it forms NBD to HBD interactions at the convex interface of each protomer (E354 of each i-th protomer makes a salt bridge with R406 of the neighboring i-1 protomer), and it is stabilized both electrostatically and by becoming buried. The SHAP plot (Supplemental Fig. S15) shows that, in this case, the absence of ligands leads to the destabilization of the inter-protomer interface, meaning that the presence of the ligands locally stabilizes L12 in the hexamer. The fact that ligand binding makes this region more flexible in the monomer might indicate that the formation of the inter-protomer NBD to HBD interactions upon hexamerization is achieved through an adaptive matching of flexible interfaces in the partner protomers, rather than through rigid interfaces docking.

In the ClpB monomer, allosteric regions are primarily observed in the NBD2, where they undergo major structural changes upon ligand binding, as indicated by RSA shifts (Supplemental Fig. S8). This is consistent with our above MSM observation that the NBD2 serves as the primary region of ligand-dependent conformational changes (Supplemental Fig. S6). Decreases in Coulomb and VDW energies observed in the ATP state of the ClpB monomer reveal the stabilization effects in the terminal loops (L29 and L32), helix H24, and β-strand B17 (Fig. 4) and highlight the interdomain allosteric coupling unique to ClpB. Consistent with these changes, the pore loops PL1 (L6) and PL3 (L19) (Supplemental Table S2) are buried in the ATP state, highlighting the strong coupling between nucleotide binding and pore-loop dynamics. Coulomb and VDW profiles also indicate that the formation of the CPX state lowers the energies of several helices (H4, H11, H14, H19, H21) and loops (L16, L22, L23, L24, L32) (Supplemental Fig. S10), including H4 adjacent to PL1 and H19 containing WB2, thereby stabilizing many ATP binding site and pore regions compared to the SUB or ATP states (Supplemental Fig. S8).

**FIG. 4.**
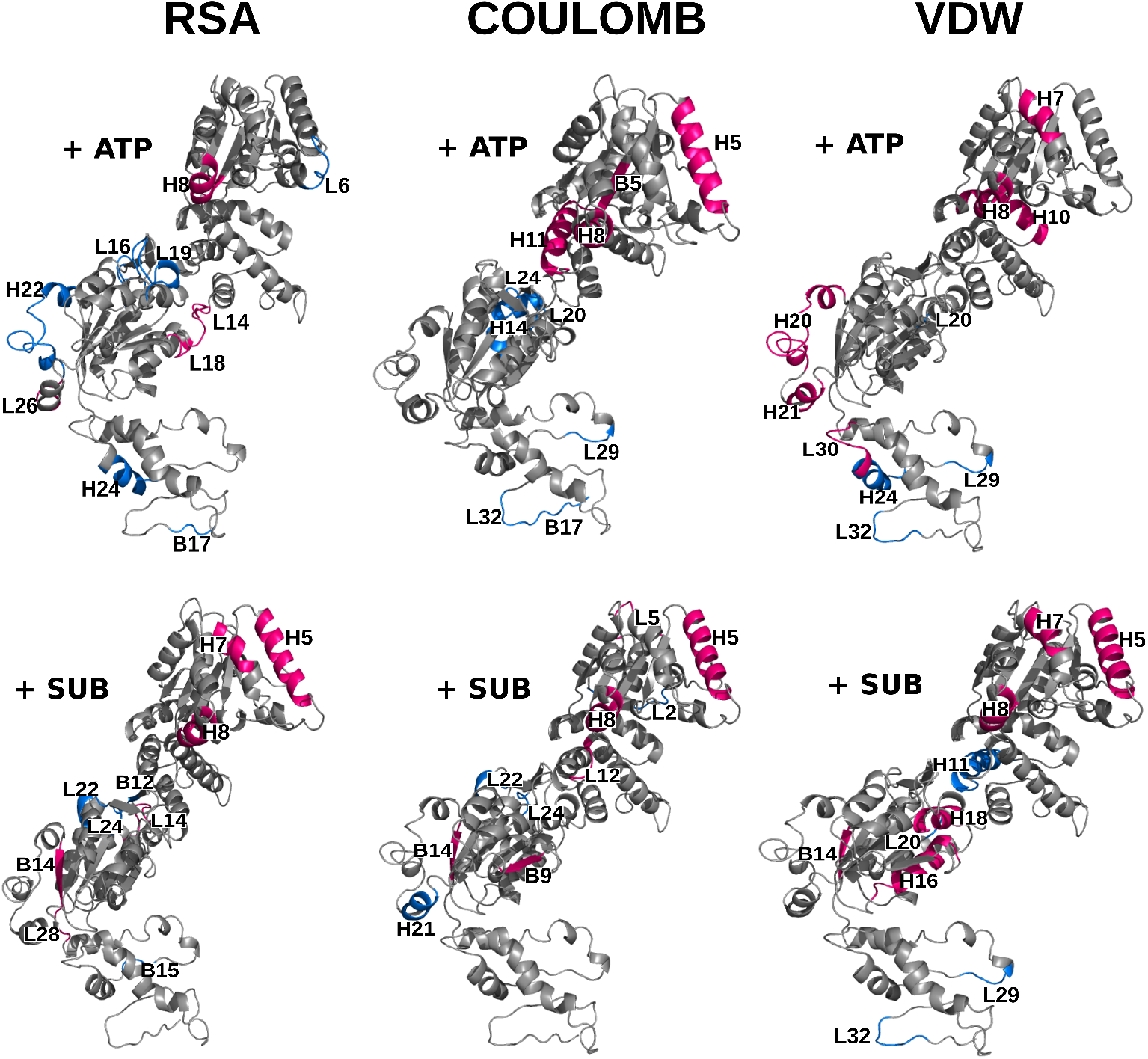
Key allosteric regions of the ClpB monomer in ligand-binding transitions. Secondary structure elements of the katanin monomer with the largest contribution in the transition due to nucleotide (ATP, top panels) or substrate (SUB, bottom) binding. Structural regions are color-coded according to the increase (red) or decrease (blue) in RSA (left panels), COULOMB (middle panels), and VDW (right panels) feature values.

Our results suggest that the binding of either ligand to the monomer increases the energy of many helices (H5, H7, H8, H10, H11) in the NBD1 compared to the NBD2, based on the Coulomb and VDW energy profiles (Supplemental Fig. S8), suggesting an energy imbalance between the two domains. This imbalance leads to instability around the ATP binding sites and pore regions in NBD1, which is mitigated upon complex formation. For example, H11 exhibits destabilized Coulomb interactions in the ATP state, but becomes stabilized in the ATP and substrate complex.

Upon substrate loss in the ClpB hexamer, VDW, Coulomb and RSA shifts indicate that many helices (H5, H8, H11, H12, H13, H14, H15, H17, H21) and loops (L9, L10, L12, L17, L21, L25) in the hexamer are destabilized, resulting in the exposure of the central pore and the weakening of the inter-protomer interactions that support substrate grip^20^ (Supplemental Figs. S11-S12). Destabilization in L10 (adjacent to Arg finger 1) and L17 (WA2 adjacent loop) indicated that the interactions with the substrate contribute to the stabilization of the ATPase and interface region of the hexamer (Supplemental Fig. S14). However, in the monomer, we observed increased VDW energy in the WA2 region (H16) upon substrate addition, suggesting that substrate binding alone is insufficient to stabilize the WA2 region and that both nucleotide binding and hexamerization are required (Fig. 4).

ATP loss in the ClpB hexamer stabilizes PL1 (L6) and H7 (adjacent to PL2), indicating that ATPase activity contributes to the stabilization of the pore environment in the hexamer^67^. Conversely, many helices (H13, H14, and H15) and the loop L32 in NBD2 are destabilized upon ATP removal (Supplemental Fig. S12). This observation is consistent with the monomer findings, where the L32 and H14 regions are stabilized in the ATP state, based on Coulomb and VDW interactions (Supplemental Fig. S8), suggesting a conserved ATP-dependent response across many regions of ClpB.

Loss of both ligands from the hexamer led to the stabilization of β-strands adjacent to the Arg Finger 1 (B6) and the WA2 (B9), as well as loops L20 (adjacent to WB2) and L8 (contains PL2), highlighting the role of the ATPase region in coupling substrate processing to hexamer stability (Supplemental Fig. S16). We observed that any perturbation leads to the stabilization of Coulomb interactions in L25 and B5 across multiple chains, highlighting ClpB’s unique inter-protomer coupling network that facilitates substrate engagement^20^.

### C. Comparing Evolutionarily Significant Residues Among ATPases

Our previous study^22^ found multiple sequence alignments to be a powerful tool for tracking functionally relevant positions. Those alignments showed that the conserved residues were evolutionarily significant for the meiotic clade, while divergent positions were indicative of spastin-specific or functional positions. When determining the importance of specific residues, the challenge we face is that the level of experimental support from mutational assays for katanin and ClpB is not as extensive as it is for spastin. This makes it difficult to narrow down which residues should be analyzed in detail. Therefore, we focused on the set of positions that we found to be highly relevant for spastin in our previous study^22,58,66^. Following this logic, we created a sequence alignment of the NBD of three AAA+ ATPases: *C .elegans* katanin (Accession ID: P34808), NBD1 of *E.coli* ClpB (E. coli, P63284), and *D.Melanogaster* spastin (A0A0B4LHJ5). Since both katanin and spastin are severing enzymes, we found that their sequences were more closely related to each other, 45.75% identity, than with ClpB, whereas ClpB was 23.11% identical to katanin and 22.87% identical to spastin (Fig. 5).

**FIG. 5.**
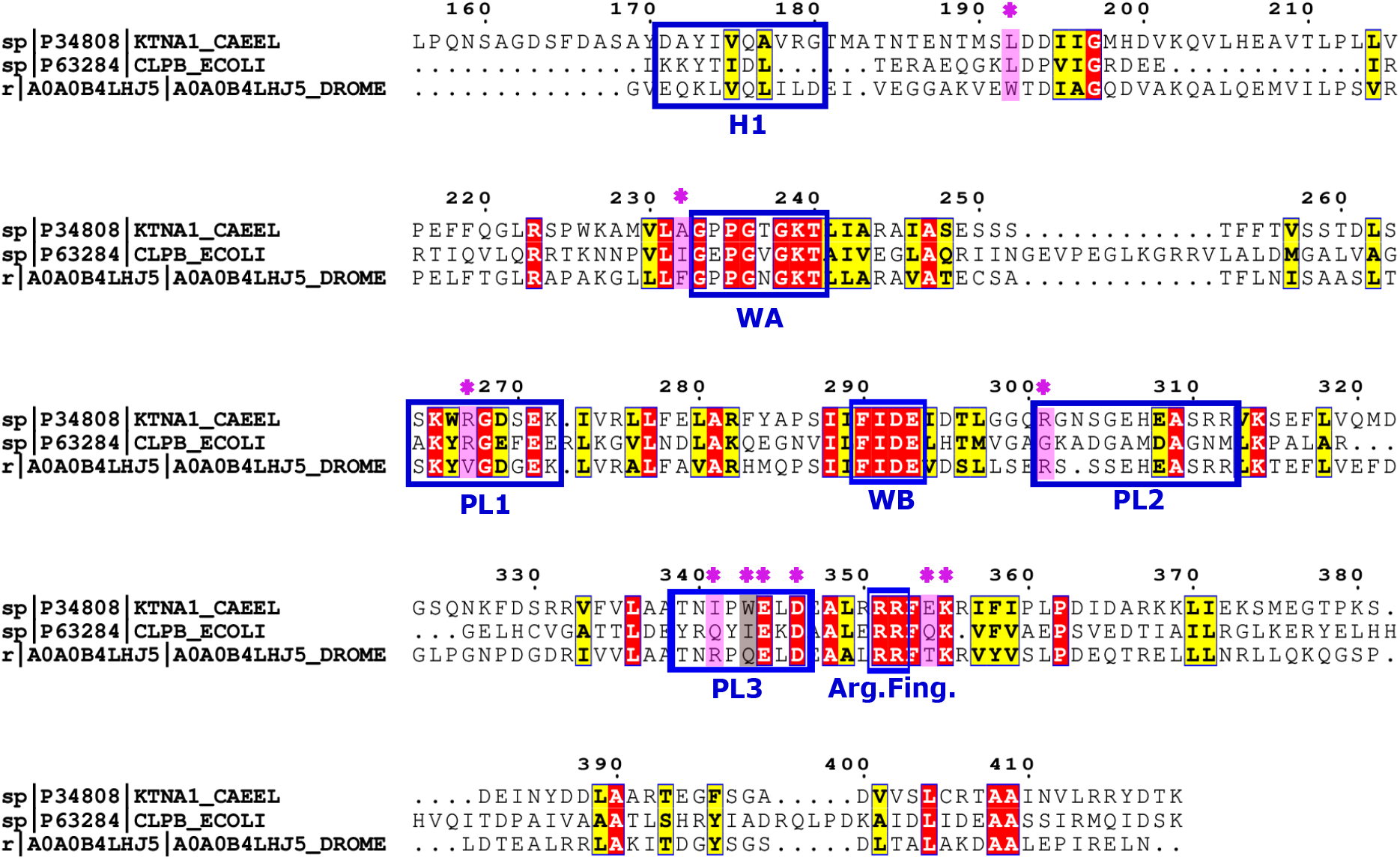
Sequence alignment of AAA+ domains of *C .elegans* katanin (P34808), NBD1 of *E .coli* ClpB (P63284), and *D .Melanogaster* spastin (A0A0B4LHJ5). Key functional motifs are indicated in blue boxes: helix H1, WA, WB, pore loops (PL1, PL2, PL3), and the arginine fingers (Arg. Fing.). Strictly conserved residues are highlighted in red, and similar residues across the three sequences are indicated in yellow. Residues with asterisks are discussed in the text.

Since spastin is associated with HSP (hereditary spastic paraplegia), we investigated whether some of the mutational sites implicated in the disease are conserved in katanin and ClpB. A key residue for spastin is its allosteric center R591^66^, which is perfectly conserved in the spastin family. The corresponding residue in katanin, R301, also remains conserved as an arginine however it is a glycine in ClpB, G287. This substitution may be related to the distinct functional roles of severing versus disaggregase proteins.

The residues that were found to be invariant in spastin (*>* 90% identity), but not in the meiotic clade in our previous study^22^ including W482, F522, V557, R630, and Q632 (shown with asterisk in Fig. 5) did not follow the same trend in katanin and ClpB (corresponding to positions L192, A232, R267, I341, and W343 in katanin and L177, I205, R252, Q321, and I323 ClpB). Residue F522 was found to be very important for spastin in terms of conservation, and especially as a structural lynchpin in the protein structure graphs (PSGs) likely due to its bulky side chain. This position is an Ala in katanin and an Ile in ClpB, which, based on the very poor conservation, indicates its uniqueness to nematoda. Residues R630 and Q632 are located in the allosterically active PL3 in spastin, which is part of a larger region that also contains highly conserved residues such as E633/E344/E324 (spastin/katanin/ClpB) and D635/D346/D326. The PL3 loop is highly conserved among spastin and katanin; all 6 residues are conserved, including N340 and P342, whose mutations are associated with HSP in spastin. However, the loop is not as highly conserved in ClpB (only 2 out of 8 residues are conserved), which is to be expected as the role of PL3 in ClpB differs from its role in severing proteins.

The most prominent region in the SHAP results for katanin was L12, which is composed of residues E354 and K355. The lysine is perfectly conserved across all three AAA+ proteins, whereas the glutamate is replaced by threonine (T) in spastin (L11) and a glutamine (Q) in ClpB (S6). This substitution suggests an evolutionary adjustment in charge polarity that modulates inter-protomer interface stability.

Overall, this analysis shows that the AAA+ ATPases have highly conserved ATP-associated regions, as expected based on their action being dependent on ATP hydrolysis, whereas the substrate interaction regions depend on the protein under study. Also, the allosteric center in spastin R591 remains conserved in katanin (R301), but switches to a Gly in ClpB (G238), which indicates that it plays a different functional role in ClpB.

### D. Impact of the Binding Partners on Dynamic Networks

Allosteric communication within a protein structure is highly sensitive to perturbations due to ligand binding. The effect of such perturbations is noticeable on relatively short timescales, which are accessible on the typical ns-microns MD simulation duration, even as large-scale conformational rearrangements of the protein occur on longer timescales of ms-s^19^. To probe the effect of ligand binding on the allosteric communication of katanin and ClpB, we developed coarse-grained representations of the allosteric network, in the presence (CPX) or absence (APO) of the binding partners, with C*_α_* positions indicating the residue locations and edges indicating inter-residue couplings (see Methods). Here, the strength of the inter-residue couplings is quantified using the corresponding terms of the Dynamic Cross-Correlation Matrix (DCCM) extracted from our MD simulations^22,54,55^. The importance of specific positions for the allosteric communication is evaluated through their relative contribution to the allosteric paths that connect pairs of amino acids, as measured by the betweenness centrality, C*_B_* (see Methods). Using this metric, we ranked the top 10% residues in katanin and 5% in ClpB, which we refer to as the highly central residues; thereby identifying positions that are essential for passing information to all of the nodes in the network. We compare and contrast the patterns of highly central residues in the monomer and in the hexamer (Supplemental Figs. S17-S18, Tables S3-S5, S15-S17). Following our above approach for the most important features, we determined how frequently each functional region has positions listed as highly central residues for each setup of both katanin and ClpB. (Fig. 6)

**FIG. 6.**
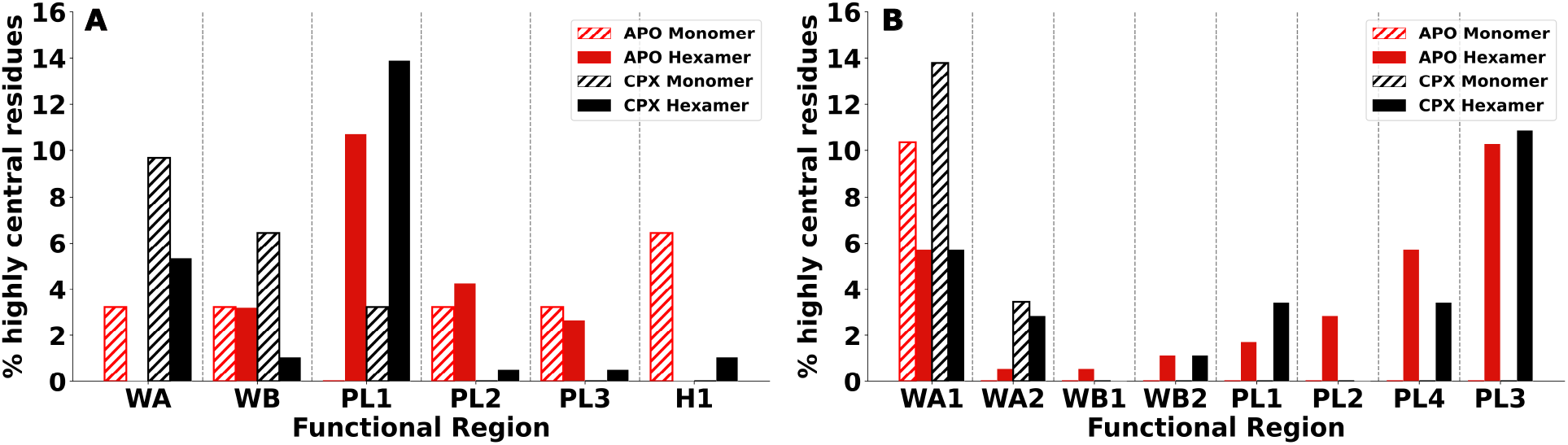
Distribution of highly central residues among functionally-important regions. The percentages of highly central residues that belong to each functional region are shown for (A) katanin and (B) ClpB in APO (red) and CPX (black) setups. Monomer and hexamer configurations are indicated by dashed and solid bars, respectively.

In the katanin monomer (Supplemental Fig. S17 a,b), the highly central residues showed distinct patterns between the APO and CPX setups, suggesting a redistribution of allosteric pathways upon the binding of ligands. Residues V175 and V178 from H1 appeared as highly central in APO, but were absent in the CPX. K239 and F290, located in the WA and WB motifs respectively, were highly central in both APO and CPX. Importantly, CPX had additional highly central WA residues, T237 and G238, suggesting that the ATP binding pocket becomes more organized in the presence of ligands. This is further supported by the fact that D292, the WB partner of K239 in a functionally important salt bridge, appeared as highly central only in CPX. Notably, S263 from PL1 and residues adjacent to this pore loop (S258, L262, and I273) were highly central in the CPX only, indicating the activation of PL1 in the presence of ligands. By contrast, PL2 only appeared highly central in APO through its position S310. Similarly, residues from PL3 and its adjacent region (A337, A338 and T339) were highly central in APO only, pointing to their possible scaffolding or pre-organization role. Taken together, this redistribution tracks the conformational dynamics previously reported for the katanin monomer^11^, in which in the APO setup HBD shows a dominant motion at the hinge region, whereas NBD and HBD move in a highly correlated, tandem fashion when both ligands are present. Because the monomer hinge (P234) sits at the edge of the nucleotide-binding pocket, inside the WA motif, the redistribution of the highly central residues to the WA and PL1 upon binding of ligands suggests that the reorganization of the catalytic core serves as the network-level signature of coordinated interdomain motions. Interestingly, while the high centrality of the WA and WB motifs agrees with our previous findings for spastin, the high centrality of the pore loops, and in particular of PL1, is unique to katanin^23^.

In the katanin hexamer (Supplemental Fig. S18a,b), WA was highly central only in the CPX setup (in chains B, C, E and F). Notably, K239 (from WA), which was highly central in both APO and CPX in the katanin monomer, showed up as highly central in chain F of the CPX setup in the hexamer. WB residues F290 and I291 appeared in chains A, C, D and F in APO, but were highly central only in the terminal chains (A and F) of the CPX setup. Residue R352, which is part of the Arg fingers, appeared in chain B of both APO and CPX setups, and both R351 and R352 were highly central in chain C of the CPX. These results strongly suggest that primarily WA and, to a lesser degree, WB motifs are important for the terminal protomers in the CPX, while the phosphate binding site (Arg fingers) is central for the interior protomers in the CPX state of the katanin hexamer.

PL1 showed virtually universal centrality across all chains of both setups (Fig. 6), with E271 appearing in chains B, C, and F in both setups and chain E of APO and chain D of CPX, and K272, present in chains B, E, and F for CPX and B, D and F for APO. This finding strongly suggests that PL1 is the only pore loop constitutively embedded in the allosteric network regardless of the ligand state. This is consistent with the principal component motions of the katanin spiral hexamer reported previously^20^, in which the dominant mode corresponds to the opening and closing of the central pore in both CPX and APO setups carried by the PL1 loops. The same study found this pore opening and closing behavior to be specific to katanin, since the PL1 loops in the spastin spiral instead undergo axial excursions. Because these PL1 dependent pore motions persist across both ligand setups, the ligand-independent centrality of PL1 in the hexamer reflects its function as a constant conduit for hexamer’s pore dynamics. This contrasts with the monomer results, where PL1 becomes central only upon ligand binding, indicating that oligomerization is responsible for the major role of PL1 in the katanin allosteric network, which aligns with the experimental finding that PL1 has a dual role in oligomerization and substrate recognition^12^. Spastin severs microtubules as a hexamer corresponding to only its AAA+ region that requires no accessory subunit^66,68^, and, as discussed in our previous work, the axial excursions of its PL1 loops, which are highly central in the hexameric state only, are likely associated with the severing action itself according to the death spiral model of severing^23^. The corresponding katanin hexamer structure solved by cryo-EM is similarly limited to only the catalytic p60 subunit (AAA+ region). However, unlike spastin, this assembly does not represent the fully active enzyme, since katanin’s microtubule affinity and severing activity are enhanced by the p80 subunit^69^. Thus, in light of the enhanced role of the PL1 loops in the allosteric network of CPX katanin, which is connected to its severing function, we propose that the difference in the pore loop motion between katanin and spastin hexamers from our study^20^ is due to the absence of the p80 subunit in katanin. Namely, we posit that the pore loops in both functionally active spastin and katanin likely execute the same type of axial excursions characteristic for severing. Importantly, these motions are very similar to those executed by the pore loops responsible for substrate processing in ClpB^20^, which strongly suggests that the axial excursions of the central pore loops are a characteristic of clade 3 AAA+ machines.

By contrast, PL2, which appeared highly central in chains B, C, and F of APO, only appeared as highly central in chain F of CPX, suggesting that ligand binding reorganizes PL2 communication specifically at the terminal protomer rather than globally across the hexamer. This result aligns well with the high centrality of PL2 in protomer F of the spastin hexamer^23^. PL3 appeared highly central in the APO setup for chains B (T343, D346) and E (T339, P342, L345), but was absent in CPX, consistent with a scaffolding role in the APO hexamer that is suppressed once the ligand-driven network is engaged.

Our results (Table S5) showed that Chain E in CPX was the only protomer characterized by simultaneous high centrality across the oligomerization interface residues I174 from H1, T339 from PL3, and C462, T465, and F469 from CT-Hlx, as well as positions from WA (P235, G238, L241), and PL1 (T260, S264, W266, D269, and K272). Chain E also showed the highest number of highly central residues in both APO and CPX setups. This distinct behavior of chain E is consistent with the structural observation from our previous work^11^, in which a DynaMut analysis of the cryo-EM E293Q construct predicted that reverting the WB glutamine to the wild-type glutamate is stabilizing in monomer E and destabilizing in monomer A, providing a clear evidence of chain E occupying a distinct structural and energetic environment.

Among the residues that were identified as stabilizing the intermediate state of the spiral-to-ring katanin hexamer transition in our previous well-tampered metadynamics study^63^, we found a clear division in their centrality patterns across the two setups. K239 from WA, Arg fingers residues R351, and R352 appear as highly central in CPX, and R356 is highly central in both APO and CPX, suggesting that their roles are reinforced upon ligand binding. R367 and D295 (next to the WB), by contrast, were predominantly central in APO, indicating that their contribution to the communication network is more important in the absence of ligands. R275 (following the PL1) showed broad centrality across chains in both APO and CPX, suggesting a ligand-independent allosteric role. The appearance of these residues as highly central is particularly noteworthy because the corresponding residues in spastin are associated with Hereditary Spastic Paraplegia mutation sites. This agreement between the network centrality, mechanistic studies of spiral-to-ring transition, and disease-associated mutations suggests that the communication analysis successfully captures functionally important allosteric hotspots that are critical for both normal activity as well as pathological dysfunction.

In ClpB, in each NBD, a larger fraction of high-centrality residues is found within the WA region compared to the WB region, which recalls our above findings for katanin. (Fig. 6 B) Notably, the fraction of high-centrality residues in monomers is nearly double than in hexamers, in both APO and CPX setups, reflecting that the role of WA1 in mediating allosteric coupling is sensitive to the oligomerization status of ClpB. These observations are consistent with previous studies which indicated that WA1 mutants promoted distinct hexamer formations compared to the wild-type^70,71^, highlighting the important role of WA1 in oligomerization. We also note that pore loop residues included high centrality residues only in hexamers, but not in monomers. In comparison to PL1 and PL2 in NBD1, PL3 and PL4 in NBD2 are more central in the ClpB hexamer communication network. Overall, these results indicate that pore loops become active in allosteric communications after oligomerization, and pore loops in NBD2 play more important roles in mediating communications than the loops in NBD1.

### E. Allosteric Communication Among Functional Regions

Detailed microscopic insight into the propagation of allosteric signals within the protein can be obtained by constructing network paths in the cross-correlation space between functionally-important regions^51^. For both katanin monomer and hexamer, we identified optimal and suboptimal paths (see Methods, Tables S6-S14) between (i) the ATP-binding pocket (WA) and the substrate-binding pore loops (PL1/PL2), and (ii) the substrate-binding pore loops (PL1/PL2) and the oligomerization sites (HBD tip). For the hexamer and monomer of ClpB, we identified optimal and suboptimal paths between (i) the ATP-binding pocket (WA1) and the substrate-binding pore loops (PL1) within NBD1, (ii) the ATP-binding pocket (WA2) and the substrate-binding pore loops (PL3) within NBD2, (iii) the ATP-binding pocket (WA2) in NBD2 and the substrate-binding pore loops (PL1) in NBD1, and (iv) the ATP-binding pocket (WA1) in NBD1 and the substrate-binding pore loops (PL3) in NBD2.

In the katanin monomer, optimal communication paths were consistently shorter in APO than in the CPX setup (Fig. 7), regardless of the destination (PL1, PL2, or the HBD tip), with the suboptimal path distributions following the same trend. This is the opposite of the results for spastin^22^ where the binding of ligands resulted in shorter path lengths. Together, these results indicate that ligand binding in the katanin monomer reorganizes the allosteric network such that the communication signal has more possible routes to reach the destination region, while the allosteric signals in the APO setup use a distinct path along the concave interface of the monomer. For the inter-domain communication from NBD to HBD (pore loops to HBD tip), a larger number of NBD secondary structure elements participated in the communication pathways (B1, L4, H3, B2, L6, B3, L8, H5, H6, B4, and L13), than HBD elements (L15 and H10), highlighting the dominant role of NBD in shaping long-range information transfer. While most highly degenerate regions were shared between the PL1 and PL2 derived paths, there were path specific distinctions as well. For example, H5 and H6 (regions flanking the PL2 in sequence) participated predominantly in the PL2-derived communication pathways. Particularly notable were the APO PL2-HBD pathways (Table S38), where loop L13 (the hinge at the C-terminal end of NBD) and H6 (next to PL2 in sequence) exhibited node degeneracy values approaching 1, indicating that these regions function as mandatory nodes, essential for ensuring inter-domain communication in the absence of ligands. A consistently prominent region was WB (with B3 and L8), which participated in most paths (Table S38) regardless of ligand state (except in APO PL2). This suggests a dual function for the WB: a canonical role in ATP hydrolysis and also a role as a central allosteric hub for the propagation of information among the functional regions.

**FIG. 7.**
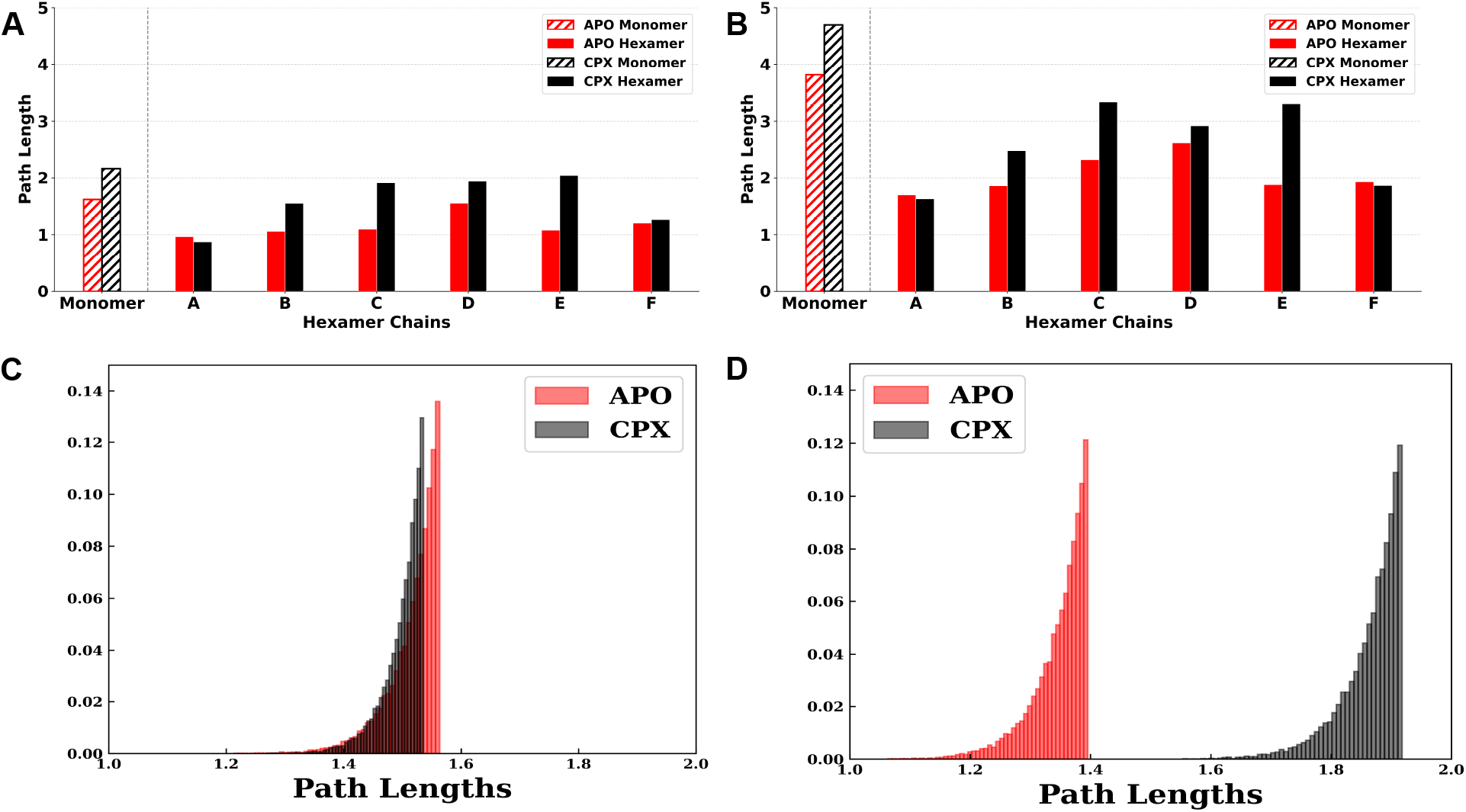
Katanin intra-protomer shortest path lengths and distributions in the APO and CPX setups. Shortest path lengths are shown for monomer (hashed) and individual protomers (chains A-F) of the hexamer in the APO (red) and CPX (black). (A) shortest paths within the NBD, from WA to PL2, and (B) shortest paths from NBD to HBD, from PL2 to HBD tip. (C) intra-protomer suboptimal path distributions in the hexamer for APO and CPX setups for chain F from WA to PL2, and (D) for chain B from WA to PL2.

Interestingly, the presence of neighboring protomers optimized the allosteric communications, as indicated by the shorter optimal and suboptimal paths for intra-protomer paths in the hexamer compared to the monomer (Fig. 7). Within the individual protomers, the monomer trends were mirrored; APO had shorter paths than CPX, with a key exception: the seam chains A and F. These chains, which possess only one neighbor and thus lack a stabilizing inter-protomer HBD-NBD contact on their exposed side, deviated significantly from the behavior of the internal protomers (B-E). In the seam chains, the allosteric signal paths were shorter than in the internal protomers and the clear distinction between the APO and CPX path lengths was lost as their distribution substantially overlapped. A similar trend is observed in the sub-optimal paths, where the path distributions for APO and CPX of the non-seam chains were well separated, whereas for the seam chains the distributions of APO and CPX overlapped (Fig. 7).

In the hexamer, the intra-protomer communication network was also found to be less redundant than in the monomer as shown by lower node degeneracy values (Tables S39-S45). There were certain outcomes unique to particular chains, such as the finding that the L7 loop (preceding WB in sequence) showed high degeneracy exclusively in chain E of the CPX setup. Critically this is the only chain in the setup where B3 (immediately following L7, that has a part of WB motif) was not degenerate. The same results also appeared in the inter-protomer CPX paths from chain E to D, where L7 again appeared as highly degenerate when B3 did not participate in the paths. This shows a compensatory behavior where L7 is recruited to maintain network integrity when B3 is disengaged from the path network. For Chain C, all intra-protomer communication paths involving PL1 were found to consistently go through L6 (immediately precedes PL1 in the sequence) of the adjacent chain B (Fig. 9). This indicates the specialized role of L6 as a conduit for PL1 mediated pathways. A similar pattern was observed for spastin where intra-protomer paths involving chain B included secondary structure elements from chain A for the signal propagation.

The inter-protomer optimal path lengths were shortest in the APO setup for the majority of internal protomers (see Figure S20), which agrees with our previous findings for spastin^23^. In the CPX setup, allosteric signals adopted flexible pathways, meaning a variety of positions showed up along the paths with low degeneracy. The only highly degenerate region for APO paths across all interfaces was L13, the inter-domain hinge, with this redundancy becoming more pronounced in the paths from F to A, which used a massive network of regions to bridge the terminal chains. Upon ligand binding, this L13-mediated pathway is largely bypassed, indicating that all the regions involved in allosteric path communication occur with low degeneracy in the CPX setup. Of all the inter-protomer optimal path distances, the interdomain route from chain D to chain C deviated most strongly between the APO and CPX setups. Figure 8 illustrates the origin of this change: it shows that the allosteric pathway connecting PL1 in chain D to the HBD tip in chain C shifts from the convex interface in the APO setup to the concave interface in the CPX setup. In the hexamer, the convex interface of each protomer interacts with both NBD and HBD of one neighboring protomer, whereas the concave interface only interacts with the NBD interface of the other neighbor. Rather than rerouting the signal within a single interface, adding ligands resulted in a complete shift in the participating interfaces.

**FIG. 8.**
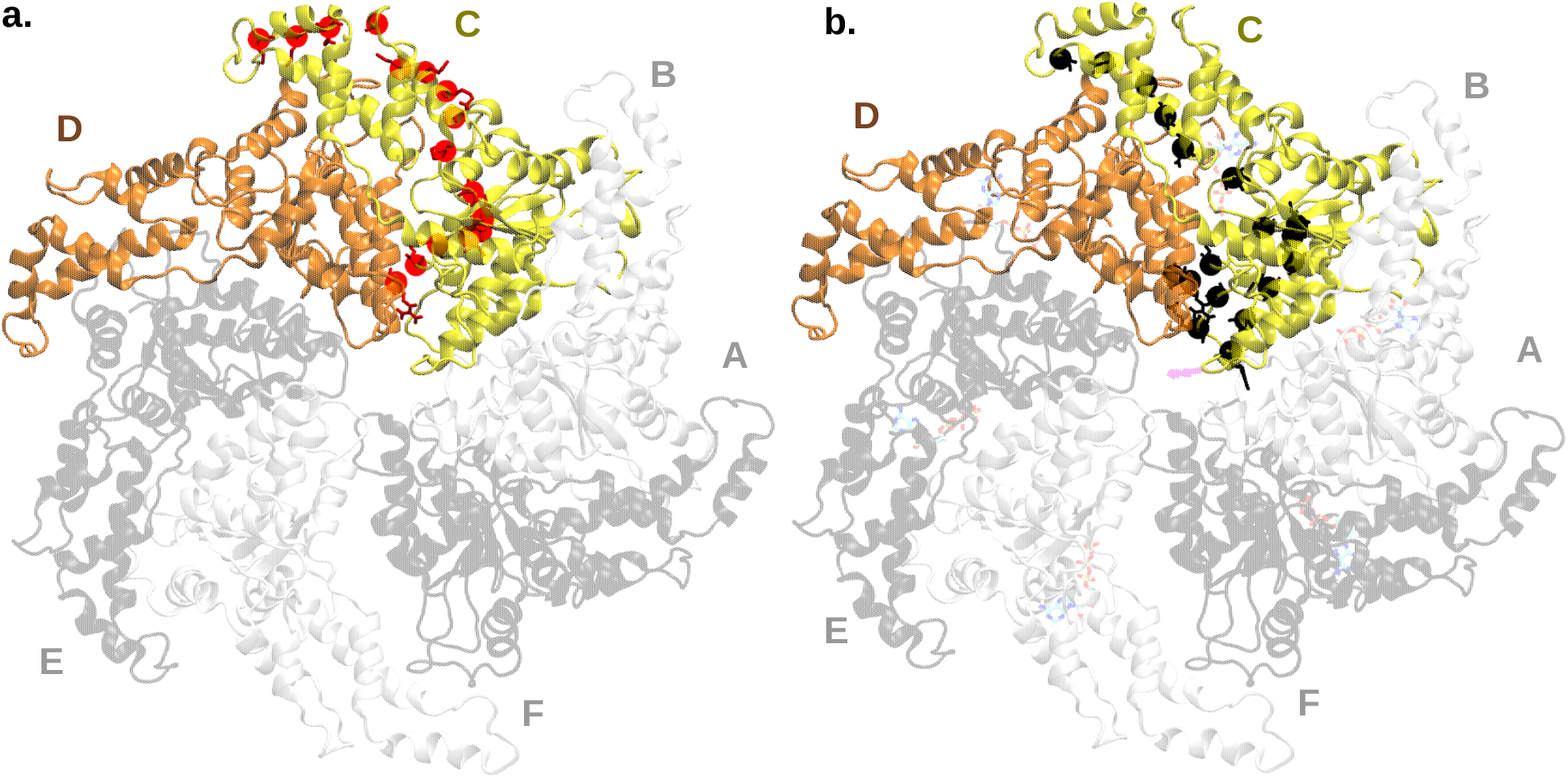
Inter-protomer pathway for chain D to C from PL1 to HBD tip in the (a.) APO, and (b.) CPX setups for katanin hexamer. The positions participating in the shortest paths are shown in red for APO and are present at the convex interface. The positions for CPX setup are present at the concave interface and are shown in black. Chain C and chain D are shown in colors yellow and blue, respectively, and the remaining protomers are colored in gray for clarity.

**FIG. 9.**
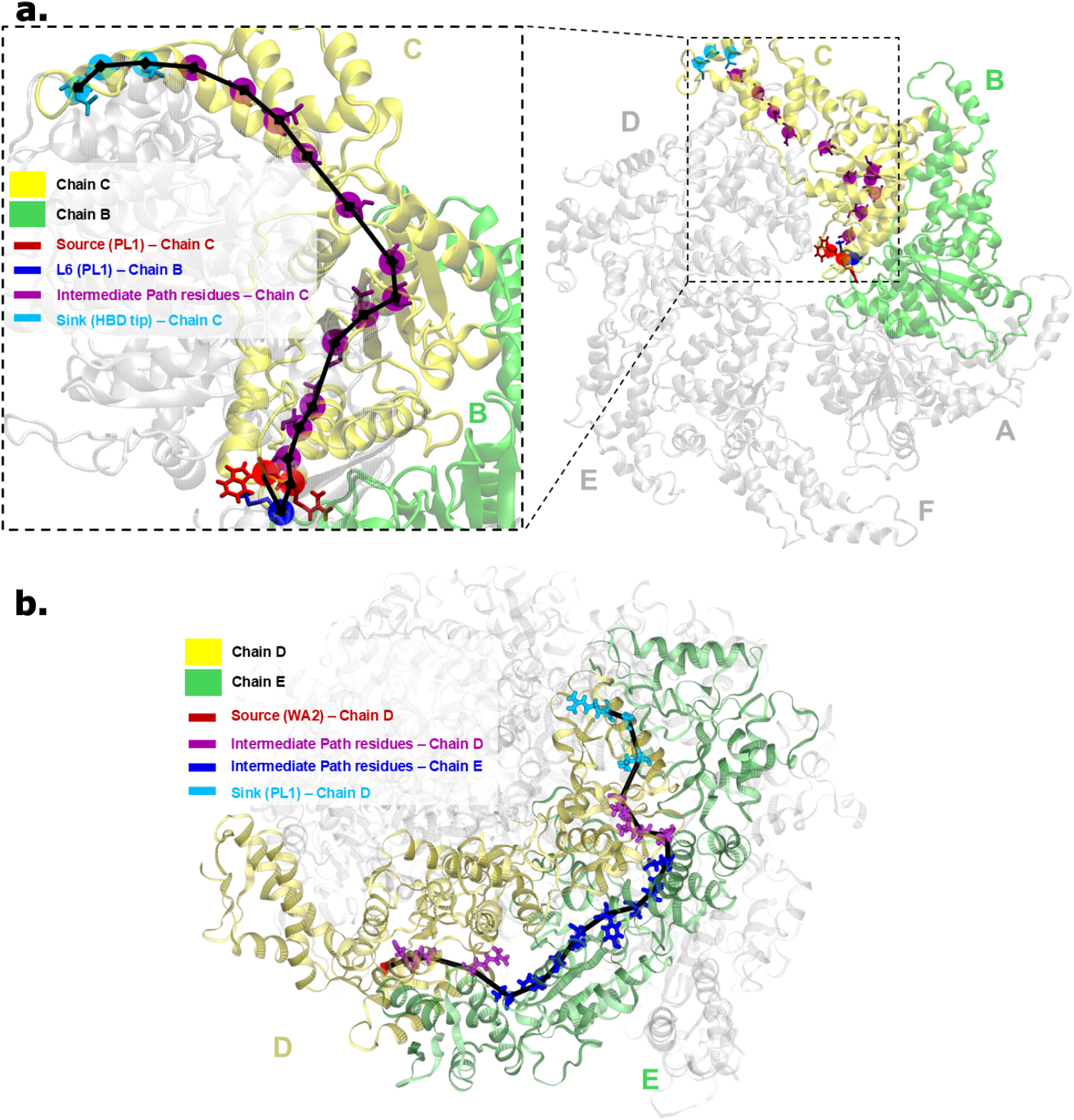
Long-range inter-domain allosteric communication in protomers of katanin and ClpB hexamers. (a.) Inter-domain pathway between PL1 and the HBD tip in the CPX setup for chain C of the katanin hexamer. The pathway (black line) begins at the source PL1 (red), in chain C, relays through L6 of the adjacent chain B (green), and propagates through intermediate residues spanning NBD and HBD (purple) before terminating at the sink, HBD tip (cyan), in chain C. The left panel provides a magnified view of residues involved in the signal propagation. (b.) Inter-domain pathway between PL1 in NBD1 and WA2 in NBD2 in the CPX setup for chain D of the ClpB hexamer. The pathway (black line) begins at the source WA2 (red), in chain D, passes through intermediate residues of chain D (yellow), traverses to residues of chain E (green) before returning to chain D, and terminating at the source PL1 (cyan), in chain D. Non-participating protomers are shown in gray.

In the ClpB monomer, for the PL1–WA1 network, we noted that the optimal path length and the average suboptimal path length were shorter in the APO than in the CPX setup (See Table S18), consistent with observations for katanin. By contrast, the opposite observation is noted for the PL3-WA2 network, where the path lengths of the CPX setup are shorter than those in the APO setup. We note that, in the APO setup, path lengths of both PL1-WA1 and PL3-WA2 networks are similar. These comparisons suggest that the two NBDs of the ClpB monomer respond differently to binding partners, and that the binding partners weaken the coupling between the catalytic site in NBD1 and PL1, but they strengthen the coupling between the catalytic site in NBD2 and PL3.

In the ClpB hexamer, intra-protomer optimal and suboptimal paths are longer, in PL1–WA1 and PL3-WA2 networks, compared with the monomer (See Tables S18-S20), which is the opposite trend compared with katanin. Comparing the APO setup with the CPX setup, in line with katanin, the paths in the CPX setup are longer than those in the APO. Notably, the path lengths in ClpB CPX setup are also longer than the APO setup in seam protomers A and F (See Tables S19-S20), while in katanin seam protomers are exceptions showing very similar lengths in the APO and CPX paths. The increase in path lengths from monomer to hexamer suggests that ClpB pore loops are regulated cooperatively in the hexamer, thus weakening allosteric communications with the nucleotide-binding site within the same NBD. Analysis of residue degeneracy within functionally-important secondary structure elements involved in these two networks revealed that the PL1 region consistently appeared as a highly degenerated source in all protomers except protomer F (See Table S28). By contrast, in the PL3–WA2 networks, no clear source region was consistently identified (See Table S29). This difference suggests that PL3 distributes couplings more broadly across the network, while PL1 communication is more localized to the source.

Inter-protomer networks within the same NBD ring indicate that the PL3–WA2 network has shorter path lengths than PL1–WA1, except for the seam protomer F in the APO setup (See Tables S23-S24), suggesting that PL3–WA2-mediated communications are stronger across adjacent protomers, similar to the intra-protomer case.

In view of the tandem-NBD architecture in ClpB, we extended our suboptimal path analysis to PL1-WA2 communications and PL3-WA1 communications to explore the inter-NBD allosteric regulations. In the monomeric ClpB, the PL3–WA1 network indicated shorter paths than the PL1-WA1 network in both APO and CPX setups (See Table S18), which we attribute to the shorter distance along the amino acid sequence between PL3 and WA1. We also found that the L12 loop at the end of NBD1 and at the NBD1-NBD2 concave was highly degenerated in PL1-WA2 and PL3-WA1 communications in both APO and CPX setups (See Table S27), indicating that it is a key region inter-NBD couplings in ClpB monomer.

In the inter-NBD communications within the same protomer in the ClpB hexamer, the NBD1–NBD2 linker region, comprising H12, H13, L14, and H14, was found as a degenerated region in all inter-NBD communications. By contrast, the L12 region only frequently shows high degeneracy in the PL3-WA2 network. More surprisingly, inter-NBD communications of nucleotide-binding sites and pore loops in the same protomer can also traverse into neighboring protomers, indicated by the appearance of degenerated secondary structure elements in the networks (see Tables S30 and S31). In the PL1-WA2 network, high-degeneracy secondary structure elements of neighboring protomers were found in all inner protomers (protomers B,C,D,E), but not in seam protomers A and F. In the PL3-WA1 network, high-degeneracy secondary structure elements of neighboring protomers were identified in protomers A, C, and E. To identify important regions that transmit inter-protomer couplings in inter-NBD networks, we identified residue pairs in optimal and suboptimal paths that are located in different protomers and mapped these inter-protomer residue pairs onto corresponding secondary structures. These secondary-structure pairs are termed as transition pairs. The percentage of transition-pair occurrence relative to the total number of paths in networks (20,000) was calculated (see Tables S32-S33). Transition pairs associated with pore loop regions, that L6–L6 of PL1 in addition to L19–L19 and L19–H17 of PL3, were frequently found and the percentages of occurrence are more than 90%, except for the PL3-WA2 network in the CPX setup. Besides, transition pairs H2–H12 and L3–H12 appeared in all intra-protomer PL1–WA2 and PL3–WA1 networks. Structurally, H2 and L3 are connected secondary structures located in the NBD1 large subdomain (NBD1L), while H12 resides within the NBD1 small subdomain (NBD1S). Our previous clustering analysis also found strong correlated motions between NBD1L in one protomer and NBD1S in the neighboring protomer^20^. Here, we identified the important secondary structure elements in neighboring protomer pairs that transmit these inter-protomer couplings by network analysis. As an example, the optimal path of PL1-WA2 communication of protomer D in the CPX setup is visualized to gain insights into the inter-protomer communication (Figure 9 b.). In the communications between PL1 and WA2 of protomer D, the optimal path traversed to NBD2 of protomer E from WA2 of protomer D and went from NBD2 to NBD1 in protomer E. At the end, the optimal path transmitted from NBD1 of protomer E to that of protomer D, and ended in the PL1 of protomer D.

In the ClpB inter-NBD communications across protomers, in which the source and sink were set to two neighboring protomers, PL1–WA2 couplings have shorter average path lengths than their intra-protomer counterparts, except for protomers B and D in the APO setup and protomer B in the CPX setup (see Supplemental Tables S21 and S25). By contrast, inter-protomer PL3-WA2 couplings reveal longer paths than in the corresponding intra-protomer case, except for protomer C of the PL3-WA2 network in the CPX setup and protomer C of the PL3-WA1 network in the CPX setup (see Supplemental Tables S20, S22, S24, and S26). The decrease in the length of PL1-WA2 paths from the intra-protomer network to the inter-protomer network implies that in the inter-NBD regulations between nucleotide-binding sites and PL1 regions, allosteric signals from neighboring protomers are stronger than the signals within the same protomer.

## IV. CONCLUSIONS

Our comparative study provides a multi-scale perspective, from the tertiary/quaternary level to the residue level, of how ATP and substrate binding regulate the allosteric communication in katanin and in ClpB. At the tertiary/quaternary scale, Markov State Models reveal that, in the absence of the nucleotide or substrate ligands, both katanin and ClpB conformations sample multiple metastates, while transitions preferentially proceed toward the ATP-populated state, suggesting that the ATP state serves as the preferred intermediate during complex formation. Diverging behavior of the two AAA+ machines is highlighted through ligand-specific metastable states of ClpB, with ATP and SUB conformations occupying distinct metastates that do not transition directly between each other. The MSM findings also highlight divergent allosteric responses of the two proteins to ligand binding. ClpB samples distinct conformational landscapes depending on the bound ligand, with structural variations primarily observed in the terminal region of the NBD2 domain, which supports the distinct role of the two NBDs suggested by earlier studies^19,20^. By contrast, katanin samples more similar conformational landscapes upon binding either ligand, with most differences concentrated in its HBD domain. Importantly, our previous MSMs for the spastin monomer also had the most notable differences within its HBD^22^, thus suggesting that the HBD changes are characteristic of the metastable states in severing proteins. Despite this similarity, spastin follows closely the behavior seen in ClpB: it adopts distinct conformational landscapes depending on the type of ligand bound. We note that neither NBD2 nor the HBD domains belong to the clade 3, thus indicating AAA+ protein-specific properties.

Our machine learning approach affords a global evaluation of the contribution of each secondary structure element to conformational transitions promoted by ligand binding. SHAP analysis revealed that the dominant allosteric regions in both katanin and ClpB are concentrated in NBDs around the nucleotide-binding site and the pore-loop regions, however in ClpB these are primarily located within NBD2. Comparison of katanin and ClpB mechanisms highlights that both proteins sense the ligand binding through similar elements yet differ in whether that signal propagates over a long distance throughout the protein assembly. In the monomers, substrate binding increased the solvent accessibility of the same region, L11 in katanin and the corresponding H8 region in ClpB, indicating a shared response within the NBD. In the hexamers, ligand removal destabilizes the inter-protomer interface. They differ in how far this destabilization reaches: in ClpB it spreads to many helices and loops, opens the central pore, and weakens the contacts that hold the substrate, whereas in katanin it is confined to the inter-protomer interface region, L12, and does not affect the pore or substrate grip. They also differ in the key determinants of the ligand states, which, in katanin, are non-catalytic and interface regions, whereas in ClpB they are the functional and pore loop regions themselves.

Our allosteric network path analysis reveals that the NBD in katanin is the primary domain sensing and propagating the ligand-induced perturbations in the protein. Rather than remaining localized at the nucleotide binding pocket, ligand-induced perturbations propagate through the NBD interface elements involved in the inter-protomer communication, supporting a model in which nucleotide binding primes the protein for oligomerization and coordinated hexamer assembly. Importantly, the network analysis also highlights residues previously implicated in stabilizing the spiral-to-ring transition intermediate, which also correspond to HSP mutation sites in spastin. The important role of these experimentally relevant positions suggests that the identified communication pathways capture key determinants of both functional conformational transitions and disease-associated perturbations. Finally, we see a differential response of the seam versus non-seam chains, to ligand removal across multiple regions and descriptors providing a region-level energetic basis for structural asymmetry previously resolved by cryo-EM, indicating that the open-spiral/closed-ring geometry of the hexamer is encoded not only structurally but also energetically. Importantly, the lengthening of both the intra-protomer and the inter-protomer allosteric paths upon the binding of both ligands likely provides the katanin hexamer with increased resilience against loss of function as applied perturbations (other ligands, mutations, changes in solvent) would more easily be dissipated by passing through a variety of alternative paths, as required in other proteins^23,26^.

The double-ring structure of ClpB highlights specific inter-NBD interactions, revealed by the cross-domain suboptimal path networks, between PL1 and WA2 and between PL3 and WA1, within the same protomer. Our node-degeneracy analysis indicated that the linker between NBD1 and NBD2, comprising secondary structures H13, L14, and H14, is an active allosteric hotspot mediating inter-NBD couplings. We note that the important role of the NBD1-NBD2 linker in interdomain communications was also recognized in other double-ring AAA+ proteins, such as the Rix7 protein^72^ and the p97 protein^73,74^. Moreover, we found that inter-NBD communication within ClpB surprisingly traversed through neighboring protomers. Such an engagement of the neighboring protomer in transmitting inter-NBD communications was also noted in the p97 protein^73^. In p97, inter-NBD interactions transit from NBD2 of one protomer to the NBD1-NBD2 linker of its neighboring protomer, and then back to NBD1 of the original protomer, in a zigzag pattern. While the inter-NBD communication in ClpB also includes this type of pattern, we note that its bridge between neighboring protomers comprises a broader region formed by the pore loops and H2, L3, and H12 in NBD1 in addition to the NBD1-NBD2 linker itself. Thus, the interprotomer crosstalk involved in the inter-NBD communication within a protomer indicates the functional cooperativity of hexameric ClpB.

## V. SUPPLEMENTARY MATERIAL

The supplementary material contains additional Results with information on Machine Learning and allosteric communication in katanin and ClpB. Tables S1 and S2 provide secondary-structure assignments and visualization for katanin and ClpB monomers. Tables S3-S17, S38-S46 present betweenness centrality analyses, optimal intra- and inter-protomer communication pathways, and node degeneracy analyses for katanin monomer and hexamer networks. Tables S18-S36 provide corresponding betweenness-centrality analyses, optimal communication pathways, and node degeneracy analyses for ClpB monomer and hexamer networks. Figures S1-S3 present structural descriptor distributions for katanin monomer and feature-convergence analyses for katanin and ClpB hexamers and monomers. Figures S4-S6 provide Markov State Model validation and metastable-state characterization of katanin and ClpB monomers. Figures S7-S16 show SHAP analyses and structural mapping of ligand-dependent allosteric responses in katanin and ClpB monomeric and hexameric states. Figures S17 and S18 identify highly central residues from betweenness-centrality analysis in katanin monomer and hexamer networks. Figures S19 and S20 compare intra- and inter-protomer communication path lengths in katanin APO and CPX networks.

## ACKNOWLEDGMENTS

We thank Amanda C. Macke for her help with setting up the GROMACS simulations for the katanin simulations. G.S. would like to thank Gilad Haran for stimulating and insightful discussions. This research was funded by the National Science Foundation (NSF) through grants MCB-1817948 and MCB-2527485 (both to R.I.D.), and MCB-2136816 (to G.S.). This work used supercomputer center resources within the ACCESS system, supported by the National Science Foundation under the Office of Advanced Cyberinfrastructure awards #2138259, #2138286, #2138307, #2137603 and #2138296, through allocation BIO210094 (to R.I.D.) and TG–MCB170020 (to G.S.).

## AUTHOR DECLARATIONS

### Author Contributions

Maryum Irshad: Data curation (equal); Formal analysis (equal); Investigation (equal); Methodology (equal); Software (equal); Validation (equal); Visualization (equal); Writing – original draft (equal). Krishan Walpalage: Data curation (equal); Formal analysis (equal); Investigation (equal); Methodology (equal); Software (equal); Validation (equal); Visualization (equal); Writing – original draft (equal). Zhaocheng Zhang: Data curation (equal); Formal analysis (equal); Investigation (equal); Methodology (equal); Software (equal); Validation (equal); Visualization (equal); Writing – original draft (equal). Maria S. Gillen: Data curation (equal); Formal analysis (equal); Investigation (equal); Methodology (equal); Software (equal); Validation (equal); Visualization (equal); Writing – original draft (equal). Ruxandra I. Dima: Conceptualization (lead); Data curation (equal); Formal analysis (equal); Funding acquisition (lead); Investigation (equal); Methodology (equal); Project administration (lead); Resources (lead); Supervision (lead); Validation (equal); Writing – original draft (equal). George Stan: Conceptualization (lead); Data curation (equal); Formal analysis (equal); Funding acquisition (lead); Investigation (equal); Methodology (equal); Project administration (lead); Resources (lead); Supervision (lead); Validation (equal); Writing – original draft (equal).

## DATA AVAILABILITY

The data that support the findings of this study are available from the corresponding author upon reasonable request.

