## Supplemental information for "Probing Mechanisms of Allosteric Regulation in AAA+ ATPases for Microtubule Severing and Protein Disaggregation"

### Results

#### Machine Learning Pinpoints Allosteric Perturbations from Ligand-Binding

In addition to the important functional regions discussed in the main text, regions including L2, L3, H5, L10, B5 and L18 in katanin reveal how ligand-driven structural changes differ between the monomer and hexamer (Figures S7, S9, S11, S13 and S15). L2 changes between the monomer and the hexamer. Its VDW interactions are stabilized by substrate alone, whereas in the hexamer, L2 showed more favorable electrostatic interactions when substrate is removed. As a loop positioned at the concave interface, L2 participates in both NBD-NBD and NBD-HBD contacts with neighboring protomers in the hexamer, indicating that the same region experiences a fundamentally different structural state in two oligomeric states. In the monomer, L3 showed decreased RSA in the presence of the ATP or of both ligands, a more buried conformation, whereas with the substrate alone the RSA increased. Electrostatic interactions involving L3 were most stable in the presence of the substrate alone, while VDW interactions were specifically destabilized in the substrate only state, suggesting that substrate engagement favors electrostatic ordering but introduces local packing strain in the absence of the ATP. In the hexamer, L3 became prominent only in the absence of substrate, where both RSA and VDW energy decreased, reflecting a more compact and favorably packed structure under substrate free conditions. H5, the helix next to the Walker B motif, exhibited a more buried conformation in the monomer as the RSA decreased for all three states. The VDW interactions were also stabilized in the presence of both ligands. In the hexamer, in the absence of substrate, the COULOMB interactions were destabilized whereas the VDW interactions became more stable in the absence of ATP. L10 showed an increase in RSA in the monomer in the presence of the ATP or of both ligands whereas COULOMB decreased in the presence of the substrate or of both ligands and VDW increased for all states. In the hexamer, both COULOMB and VDW decreased in the absence of ATP, whereas absence of substrate destabilized the VDW. For B5 in the monomer, RSA and VDW increased in the presence of ATP whereas in the hexamer the electrostatic interactions were stabilized in the absence of either the ATP or both ligands. L18 became more exposed and showed an increase in VDW in the presence of ATP in the monomer. While it showed the same RSA trend in the hexamer for the seam chains, the RSA increases for the internal chains. The VDW interactions had the same trend in the absence of substrate, where it stabilized for the seam chains and destabilized for the internal chains. Overall in the katanin monomer, L2, L3, H5, L10, B5, and L18 respond to ligand presence, mainly ATP, through changes in burial and packing within a single subunit. In the hexamer, the same regions respond instead to ligand absence, and their behavior is shaped by inter-protomer interface contacts and by seam versus internal chain positions.

In ClpB, additional changes were observed in the Arg Finger 2 region (H22) upon ATP binding (Table S2), where the region becomes buried but is packed into a less favorable local environment, based on RSA and VDW changes (Figure S8). No such changes were observed in the SUB state, highlighting ClpB's ligand-dependent conformational landscape. Additional responses in the pore region were also observed during CPX formation. The H5 (adjacent to PL1) and H18 (adjacent to PL3) helices become more exposed during CPX formation, modulating the signaling to the pore loops (Figure S10). Moreover, the L24 and the terminal loops (L29, L32) exhibited lower Coulombic and VDW energies in the SUB or NUC states, confirming the region's affinity for either ligand (Figure 4). Substrate engagement produced additional responses within the ATPase and interdomain regions. Upon substrate engagement, L12, which neighbors WA1, and H16, which contains WA2 (Table S2), shows destabilized Coulomb or VDW interactions, highlighting substrate-induced allosteric changes in the ATPase region (Figure S8). Binding of both ATP and substrate to the monomer increases VDW energy (Figure S10) in the interdomain linker region (L13 and H13), suggesting the need for hexamerization for stability. The loop L14 (adjacent to H13) becomes more exposed upon addition of both ligands. Upon ATP removal, differential Coulombic responses were observed in katanin and ClpB hexamers, especially around H6 in katanin and the corresponding region H7 in ClpB, where the region became stabilized in ClpB, but not in katanin (Figure S11 and S12). Interestingly H7 is destabilized in all ligand-bound monomer states (Figure S8). ATP loss causes the L1 linker to become buried and stabilize VDW interactions, suggesting that steric rearrangements around the pore opening are coupled to perturbations at ATPase sites. RSA plots indicated that structural changes induced by ATP loss are primarily observed in the seam protomers (A and F), suggesting that ATP-induced conformational changes are mostly local in ClpB (Figure S12). Additional responses were observed following substrate removal from the hexamer. The loop adjacent to Arg Finger 1 (L10) became more exposed with destabilized VDW interactions due to substrate loss and causing instability at the E-F interprotomer interface (Figure S14). The VDW profile primarily shows weakened interactions within the seam protomers (A and F), with increased VDW energies in several helices (H5, H13, H14, H15, H21) indicating overall destabilization upon substrate removal (Figure S12). Conversely, decreased

Coulomb and VDW energies around PL2 (L8) reveal local stabilization within the pore-loop environment despite the global shift toward destabilization.

### Tables

| | | | | | |
|---|---|---|---|---|---|
| L1 | 156 - 170 | L7 | 285 - 287 | L13 | 359 - 362 |
| H1 | 171 - 180 | **B3 (WB)** | 288 - 291 | H8 | 363 - 375 |
| L2 | 181 - 198 | **L8 (WB)** | 292 - 293 | L14 | 376 - 384 |
| H2 | 199 - 215 | H5 | 294 - 299 | H9 | 385 - 393 |
| L3 | 216 - 227 | **L9 (PL2)** | 300 - 308 | L15 | 394 - 398 |
| B1 | 228 - 232 | **H6 (PL2)** | 309 - 322 | H10 | 399 - 416 |
| **L4 (WA)** | 233 - 239 | L10 | 323 - 333 | L16 | 417 - 424 |
| **H3 (WA)** | 240 - 249 | **B4 (PL3)** | 334 - 339 | H11 | 425 - 436 |
| L5 | 250 - 253 | **L11 (PL3)** | 340 - 346 | L17 | 437 - 439 |
| B2 | 254 - 257 | **H7 (Arg)** | 347 - 353 | H12 | 440 - 449 |
| **L6 (PL1)** | 258 - 268 | L12 | 354 - 355 | L18 | 450 - 455 |
| **H4 (PL1)** | 269 - 284 | B5 | 356 - 358 | **H13 (CT)** | 456 - 472 |

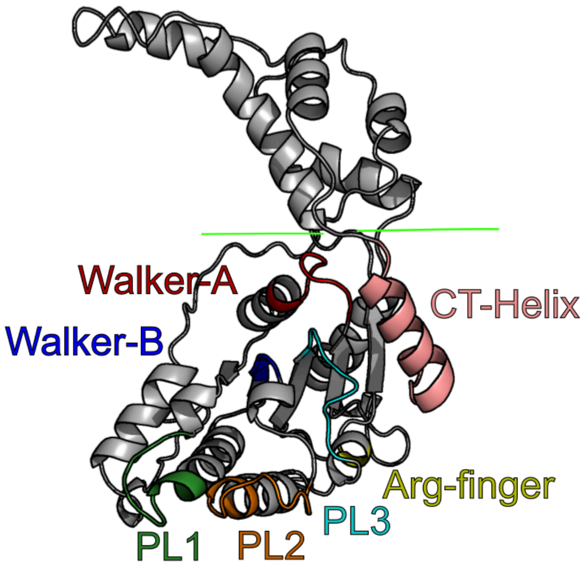


Table S1. Secondary Structure labels for katanin (PDB:6UGD). Helices (H) consist of at least 4 consecutive residues, loops (L) consist of at least 2 consecutive residues, and beta strands (B) consist of at least 3 consecutive residues. Functional regions are colored in the table and figure of the katanin monomer. The green line represents the NBD-HBD border. The cryo-EM structure assigns residues 399 to 436 as a single long helix. However, the per-residue helix probability across residues 417-424 is very low (0.03 in the online entry) increasing to 0.27 only at residue 425. We therefore assigned these as separate secondary structure elements H10, L16, and H11, which shifted the assignment such that the helix H12 in the original structure corresponds to H13 in our work.

| | | | | | |
|---|---|---|---|---|---|
| L1 | 161-164 | L11 | 339-346 | H20 | 685-693 |
| B1 | 165-167 | H10 | 347-362 | L22 | 694-697 |
| H1 | 168-174 | B7 | 363-365 | B11 | 698-700 |
| L2 | 175-182 | H11 | 366-381 | L23 | 701-705 |
| H2 | 183-194 | L12 | 382-386 | B12 | 706-708 |
| L3 | 195-201 | H12 | 387-408 | L24 | 709-712 |
| B2 | 202-206 | L13 | 525-531 | B13 | 713-717 |
| L4 | 207-212 | H13 | 532-544 | L25 | 718-721 |
| H3 | 213-226 | L14 | 545-554 | H21 | 722-728 |
| L5 | 227-235 | H14 | 555-567 | L26 | 729-733 |
| B3 | 236-240 | L15 | 568-573 | H22 | 734-756 |
| H4 | 241-246 | H15 | 574-589 | L27 | 757-758 |
| L6 | 247-253 | L16 | 590-598 | B14 | 759-762 |
| H5 | 254-268 | B8 | 599-604 | L28 | 763-766 |
| L7 | 269-272 | L17 | 605-609 | H23 | 767-788 |
| B4 | 273-278 | H16 | 610-624 | L29 | 789-792 |
| H6 | 279-284 | L18 | 625-628 | B15 | 793-795 |
| L8 | 285-298 | B9 | 629-633 | H24 | 796-805 |
| H7 | 299-305 | H17 | 634-648 | L30 | 806-813 |
| L9 | 306-310 | L19 | 649-660 | H25 | 814-835 |
| B5 | 311-313 | H18 | 661-669 | L31 | 836-842 |
| H8 | 314-323 | L20 | 670-672 | B16 | 843-847 |
| L10 | 324-325 | B10 | 673-677 | L32 | 848-853 |
| H9 | 326-333 | H19 | 678-682 | B17 | 854-857 |
| B6 | 334-338 | L21 | 683-684 | | |

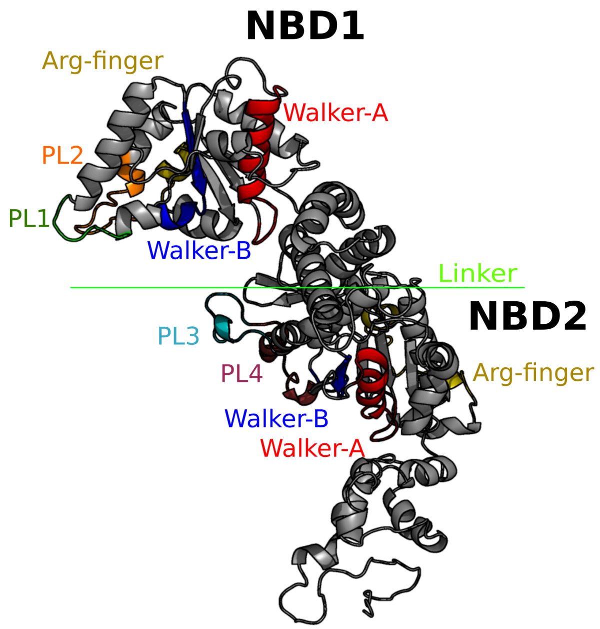

Table S2. Secondary Structure labels for ClpB (PDB:6OAX). Helices (H) consist of at least 4 consecutive residues, loops (L) consist of at least 2 consecutive residues, and beta strands (B) consist of at least 3 consecutive residues. Functional regions are colored in table and figure of ClpB monomer. The green line represents a five-glycine linker replacing the M-domain.

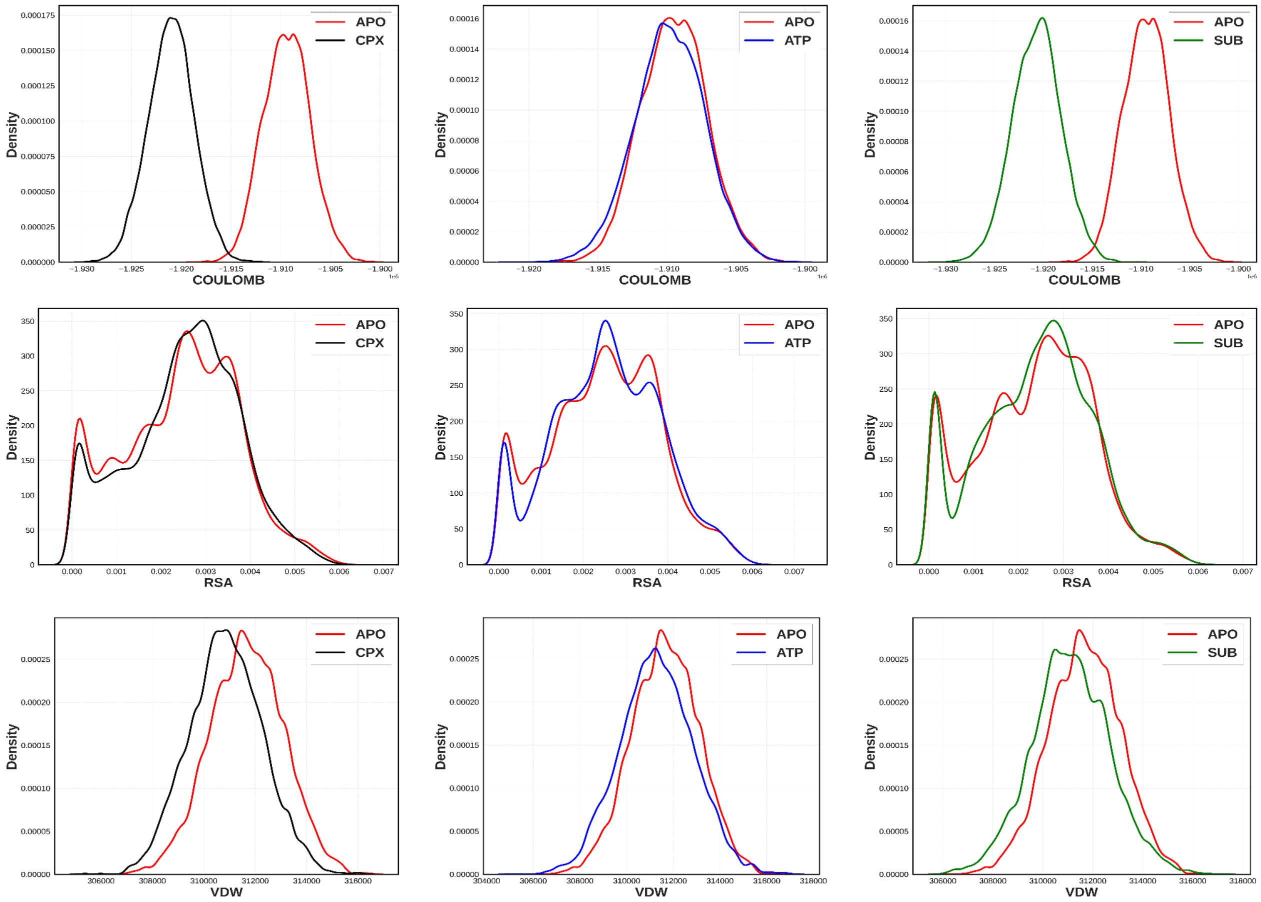


Figure S1. Distribution of structural descriptors across ligand states in katanin monomers. Kernel density estimates of Coulomb energy (top), relative solvent accessibility (RSA, middle), and van der Waals (VDW) energy (bottom) across APO vs. CPX (left), APO vs. ATP (center), and APO vs. SUB (right) conditions. Separation between distributions indicates the discriminatory power of each descriptor in distinguishing ligand-bound from apo state

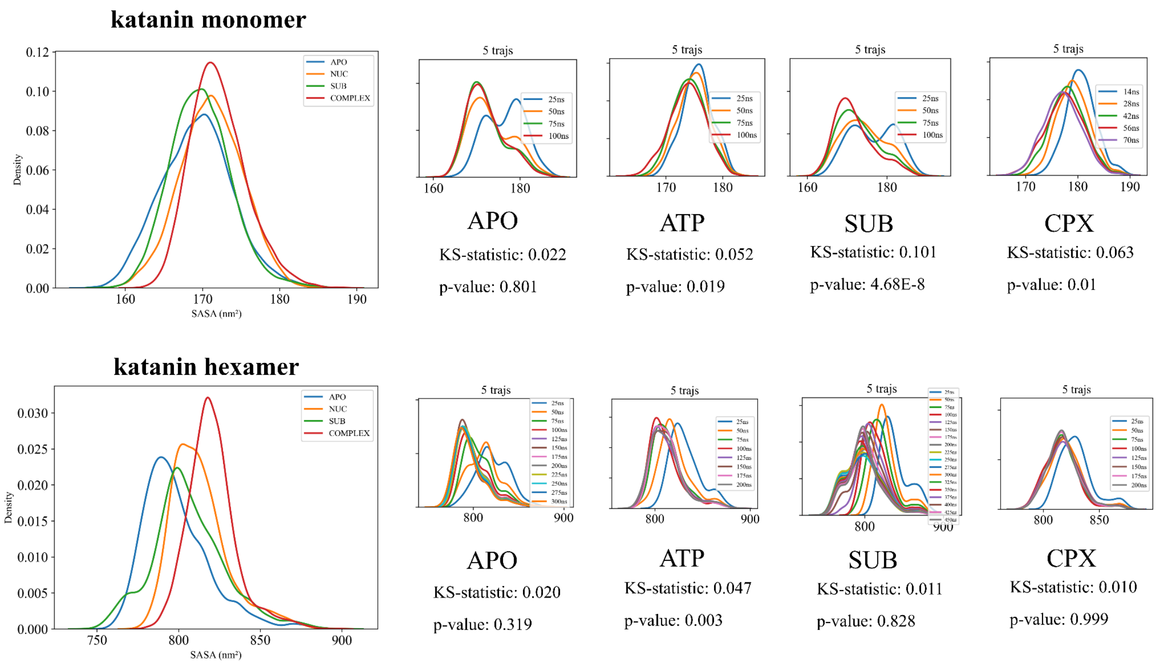


Figure S2. SASA feature convergence for katanin monomer and hexamer simulations.

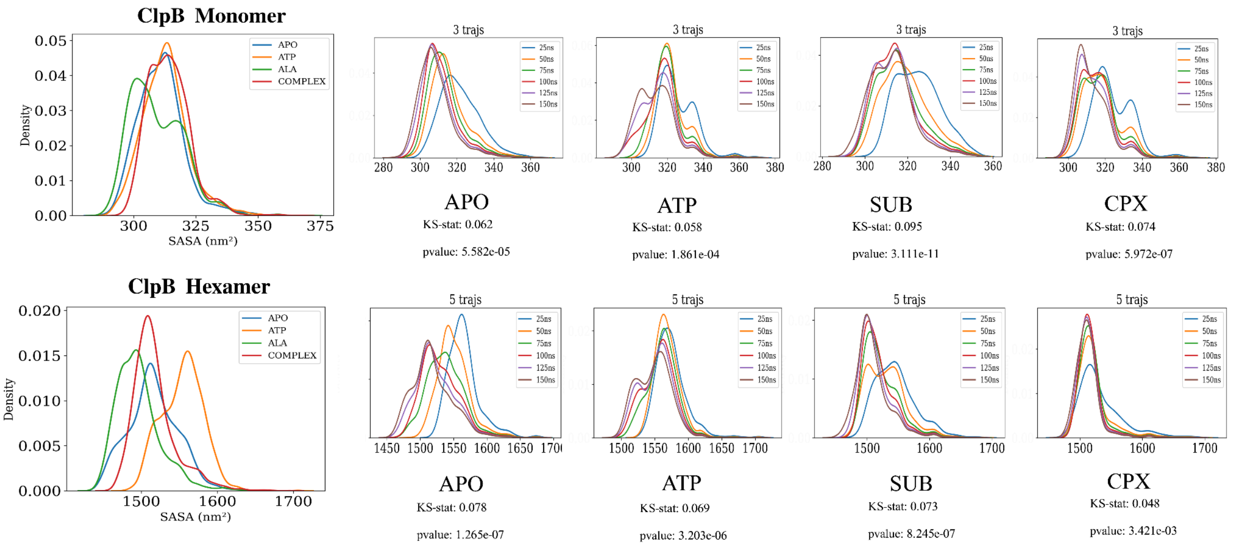


Figure S3. SASA feature convergence for ClpB monomer and hexamer simulations.

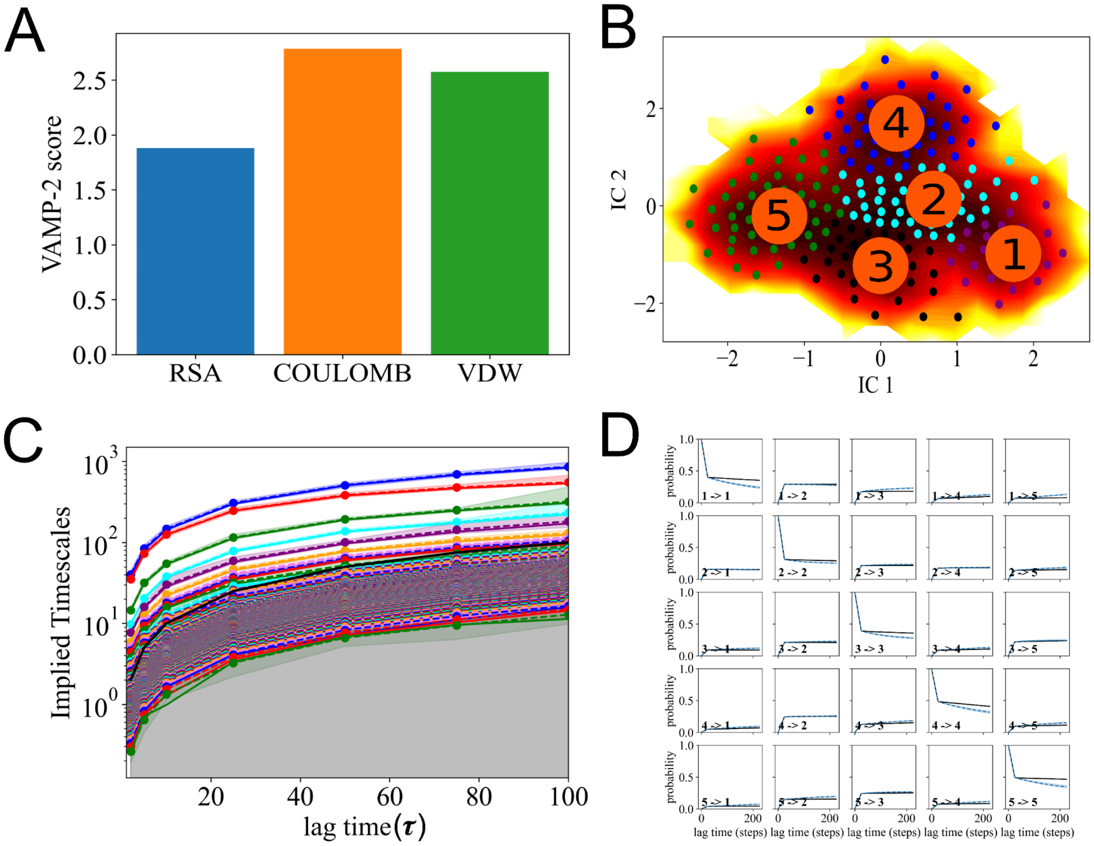


Figure S4. MSM outputs and validation for katanin. (A) VAMP-2 score for reducing each feature subspace. (B) 5 macrostate clustered defined in MSM in TICA (IC) component space. (C) Implied timescales plot for each microstate shown in B vs lag time where $\tau$ = simulation frames at every 200ps. (D) Chapman-Kolmogorov (CK) test for 5 macrostates.

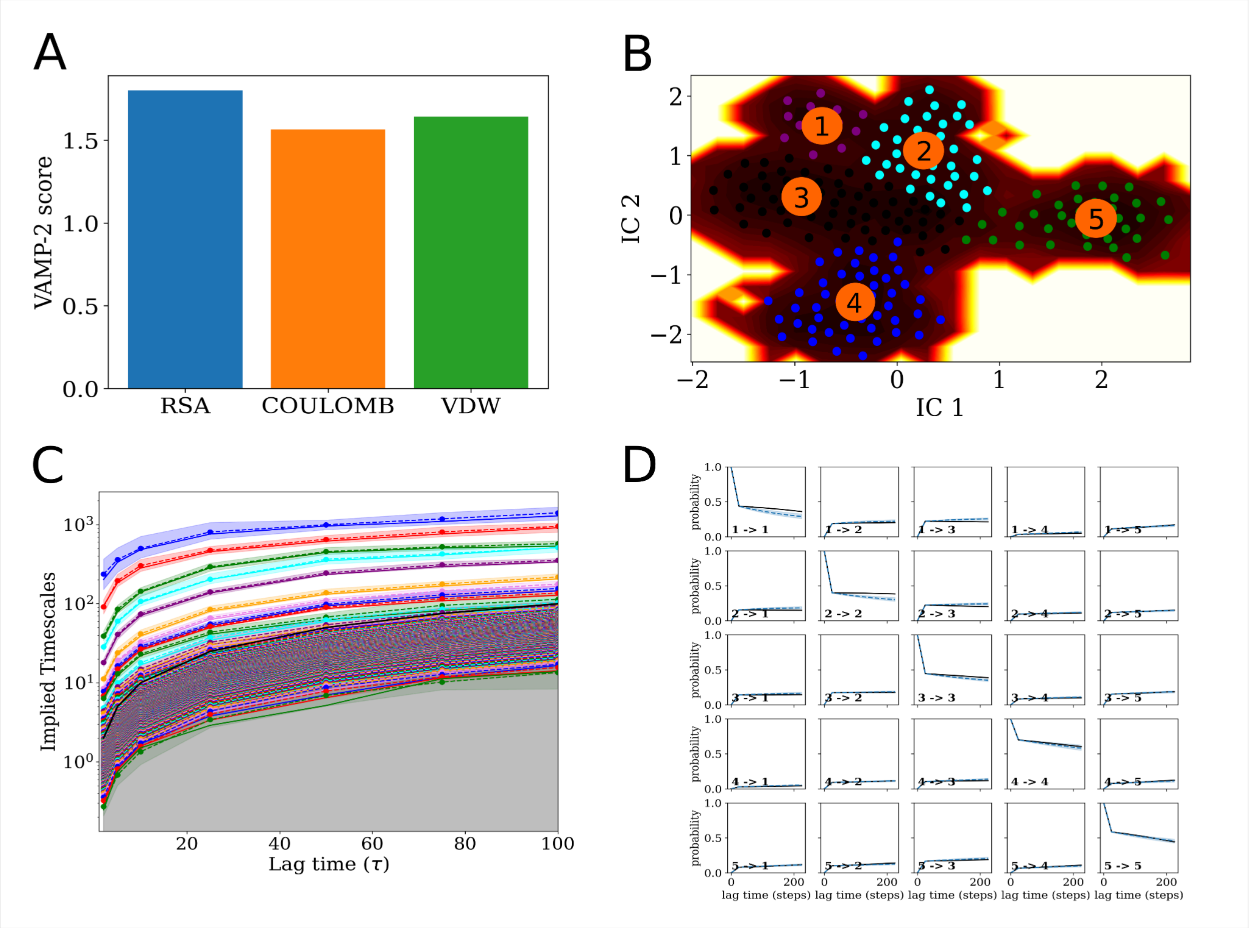


Figure S5. MSM outputs and validation for ClpB. (A) VAMP-2 score for reducing each feature subspace. (B) 5 macrostate clustered defined in MSM in TICA (IC) component space. (C) Implied timescales plot for each microstate shown in B vs lag time where $\tau$ = simulation frames at every 200ps. (D) Chapman-Kolmogorov (CK) test for 5 macrostates.

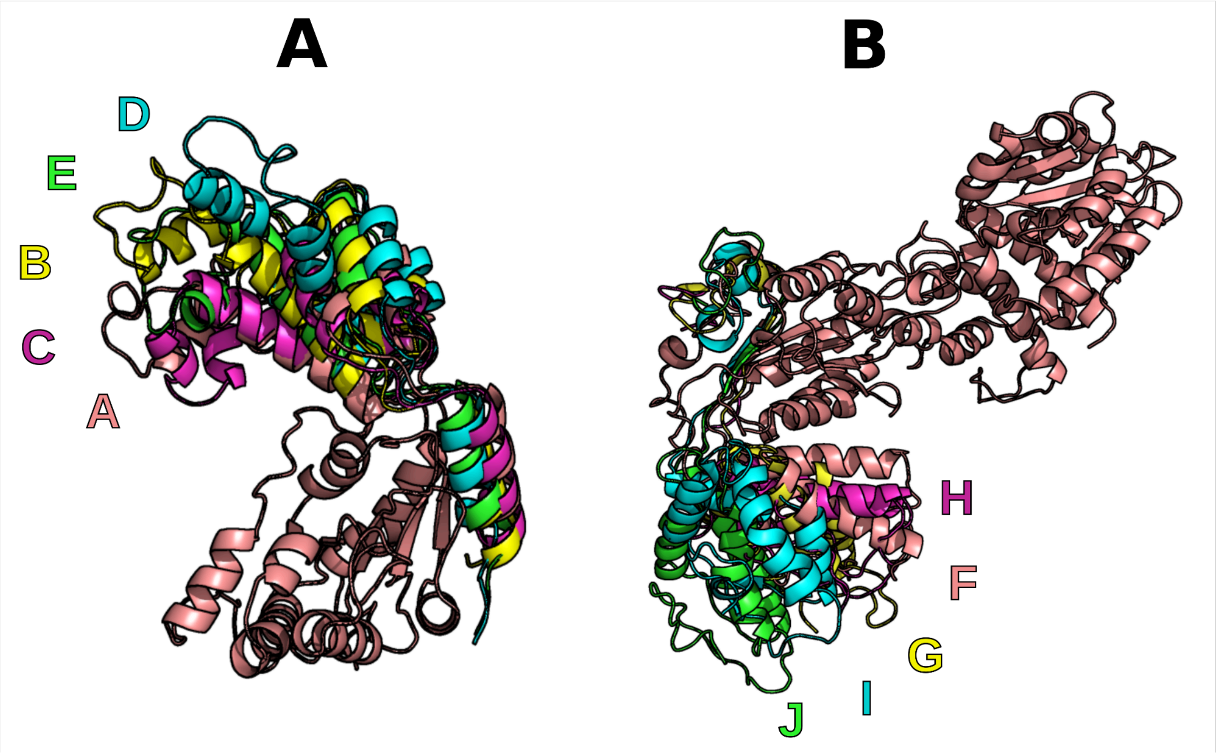


Figure S6. (A) MSM metastable states A–E for katanin. The full monomer is shown only for metastate A, while the HBD is displayed for other metastates and aligned to A. (B): MSM metastable states F–J for ClpB. The full monomer is shown only for metastate F, while the terminal region of NBD2 is displayed for other metastates and aligned to F.

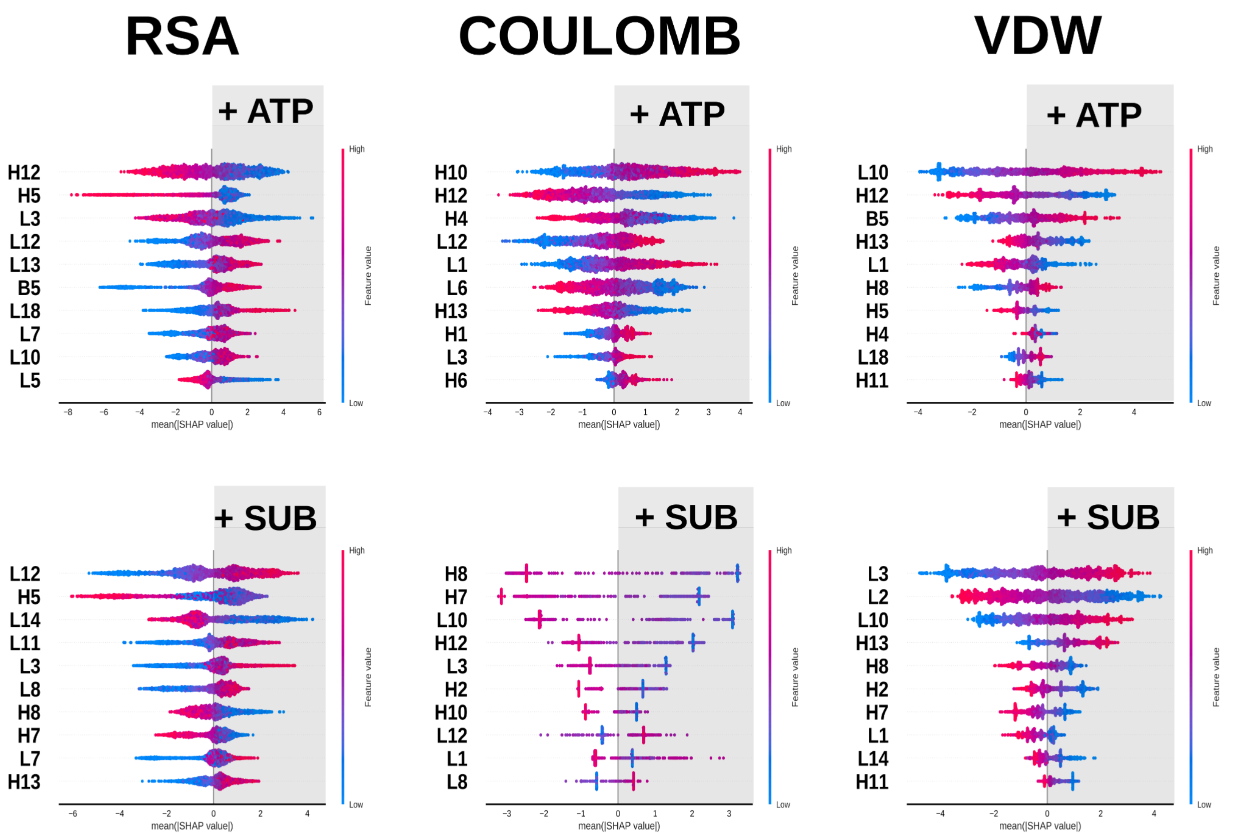


Figure S7. SHAP beeswarm plots for the transitions (addition of SUB or ATP) in the katanin monomer. Feature points with positive SHAP values indicate the corresponding feature value increases (magenta) or decreases (blue) of key regions upon the addition of ligands.

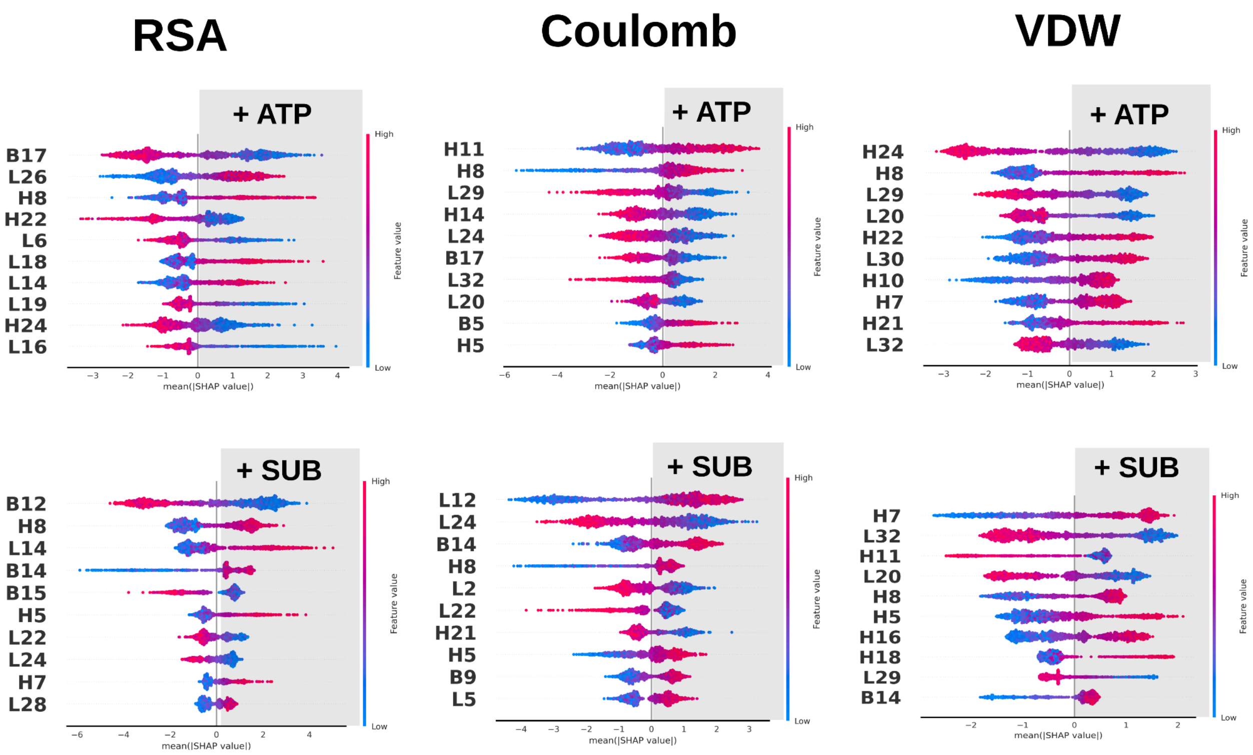


Figure S8. SHAP beeswarm plots for the transitions (addition of SUB or ATP) in the ClpB monomer. Feature points with positive SHAP values indicate the corresponding feature value increases (magenta) or decreases (blue) of key regions upon the addition of either ligand.

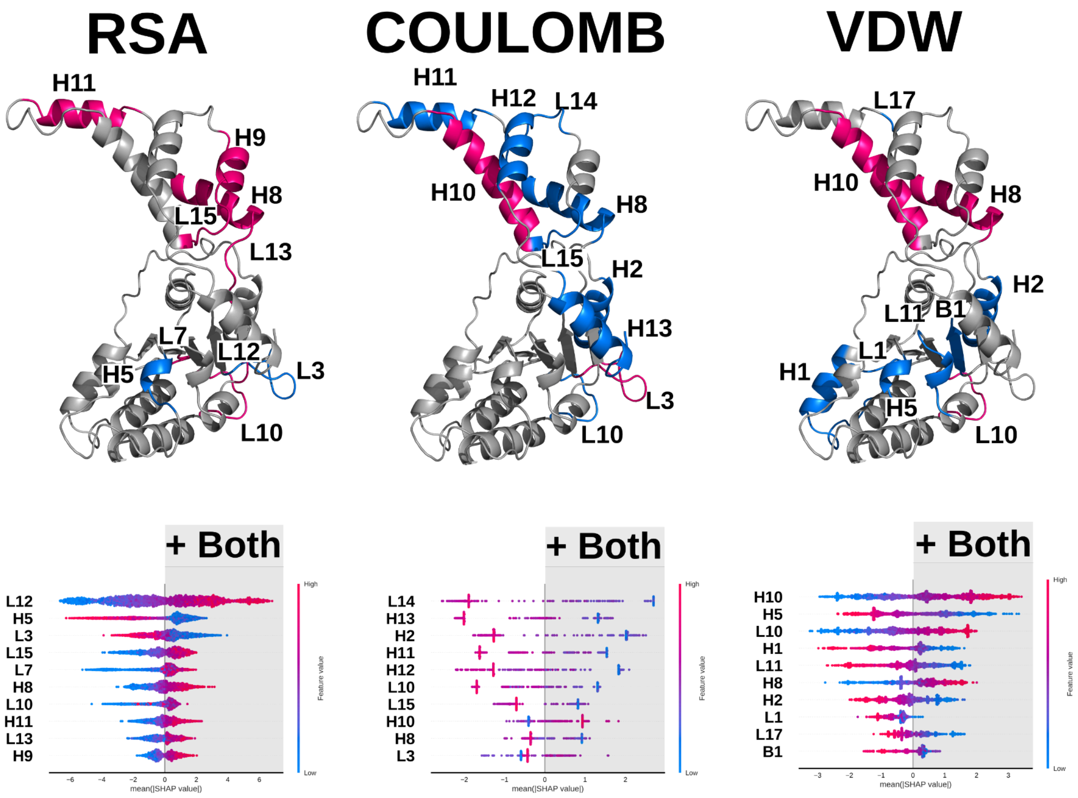


Figure S9. Representative illustrations showing key regions in the katanin monomer associated with the binding of both ligands (SUB and ATP). Regions highlighted in pink indicate an increase in feature values, while those in blue indicate a decrease. The bottom panels display the corresponding SHAP beeswarm plots, where positive SHAP values reflect the same directional changes of these key regions.

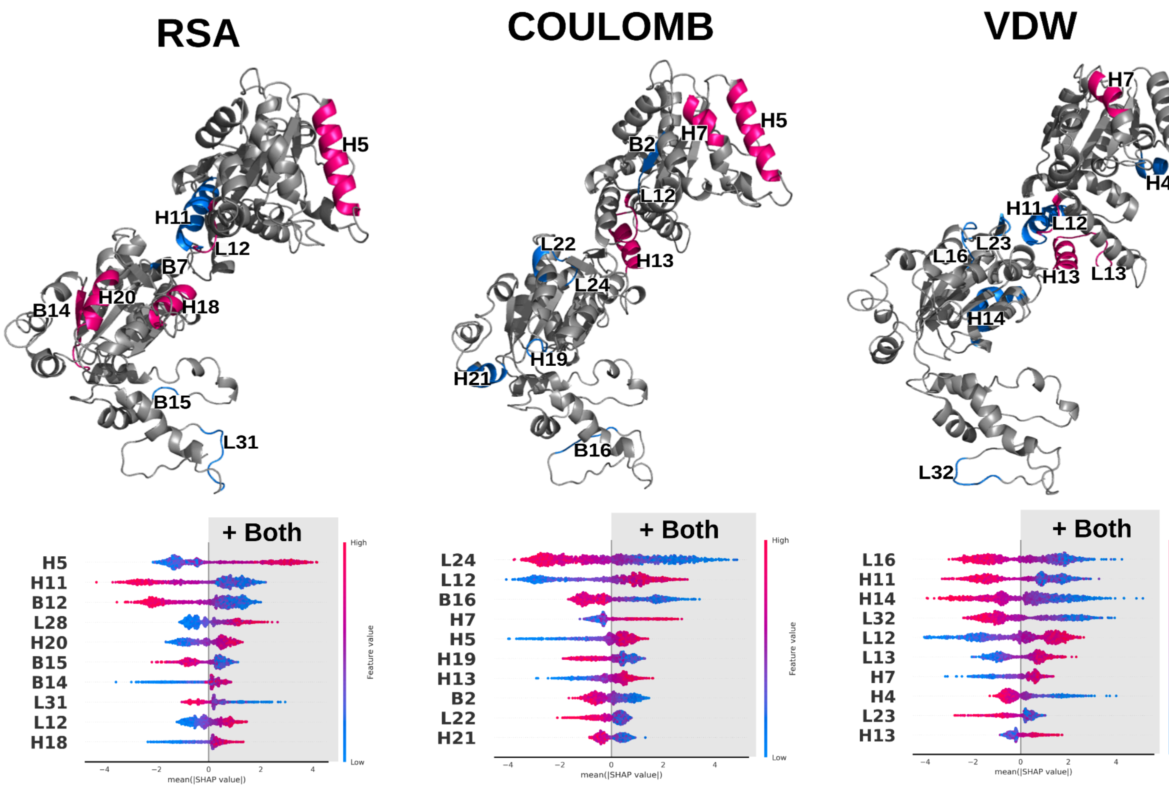


Figure S10. Representative illustrations showing key regions in the ClpB monomer associated with the binding of both ligands (SUB and ATP). Regions highlighted in pink indicate an increase in feature values, while those in blue indicate a decrease. The bottom panels display the corresponding SHAP beeswarm plots, where positive SHAP values reflect the same directional changes of these key regions.

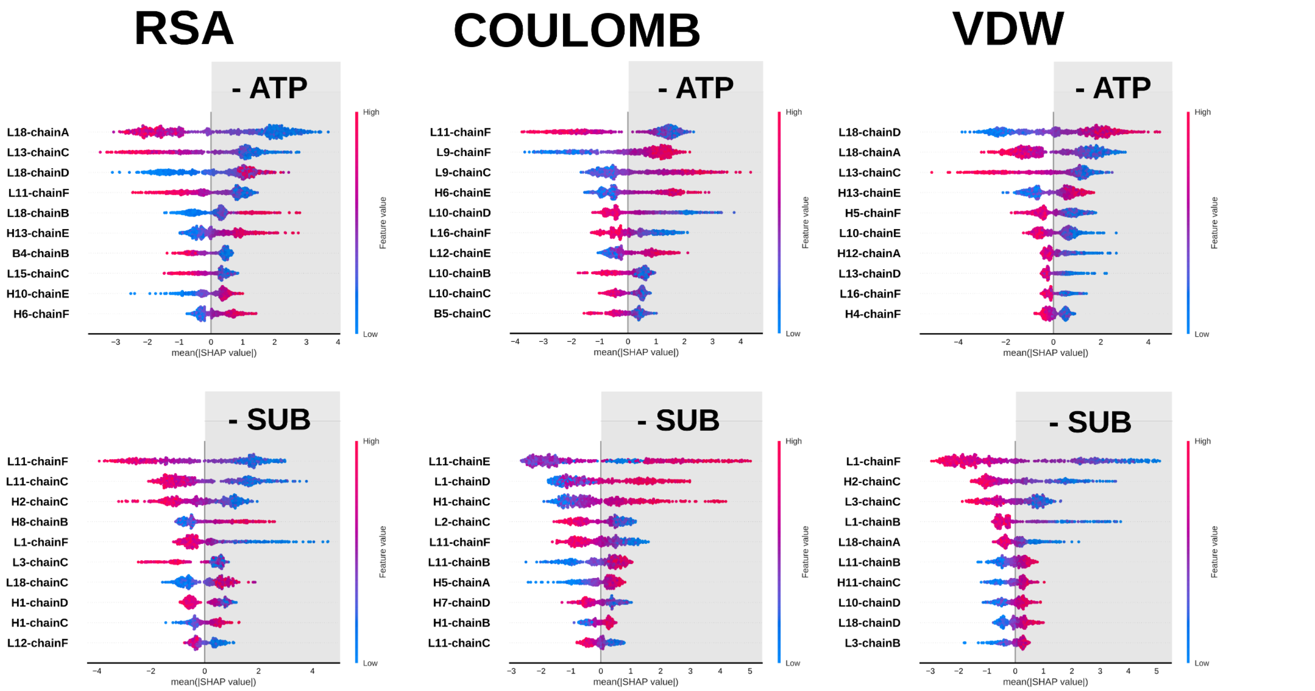


Figure S11. SHAP beeswarm plots for the transitions (removal of SUB or ATP) from the katanin hexamer. Feature points with positive SHAP values indicate the corresponding feature value increases (magenta) or decreases (blue) of key regions upon the removal of either ligand.

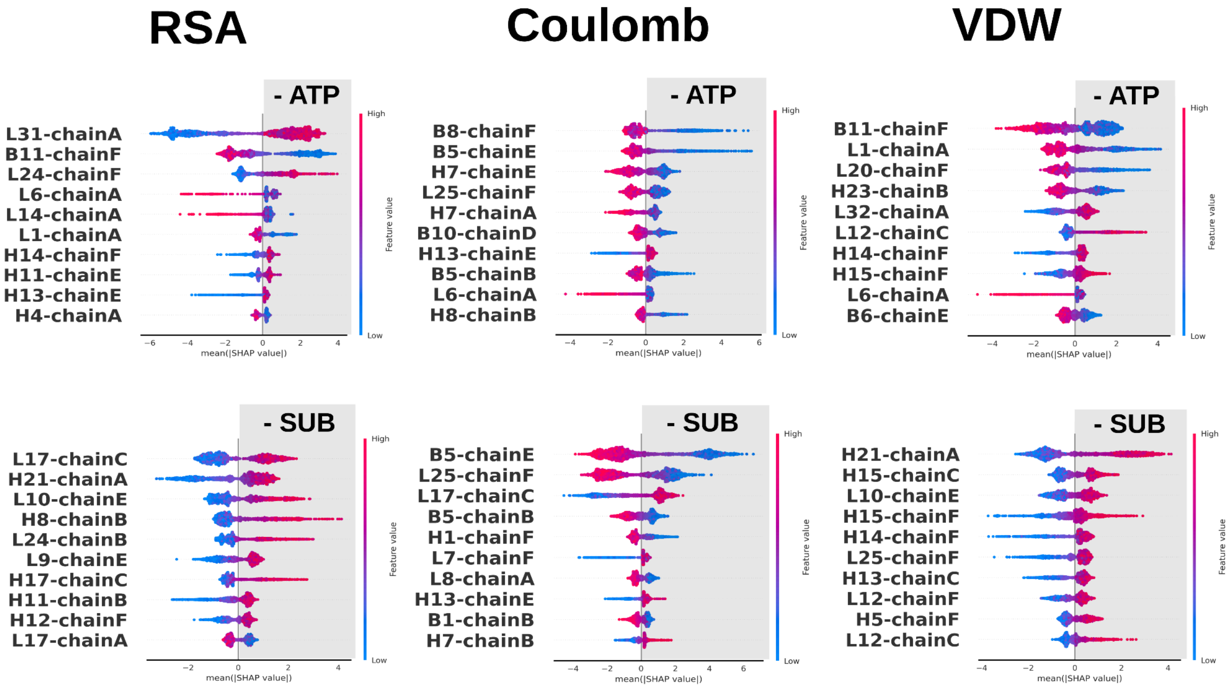


Figure S12. SHAP beeswarm plots for the transitions (removal of SUB or ATP) from the ClpB hexamer. Feature points with positive SHAP values indicate the corresponding feature value increases (magenta) or decreases (blue) of key regions upon the removal of either ligand.

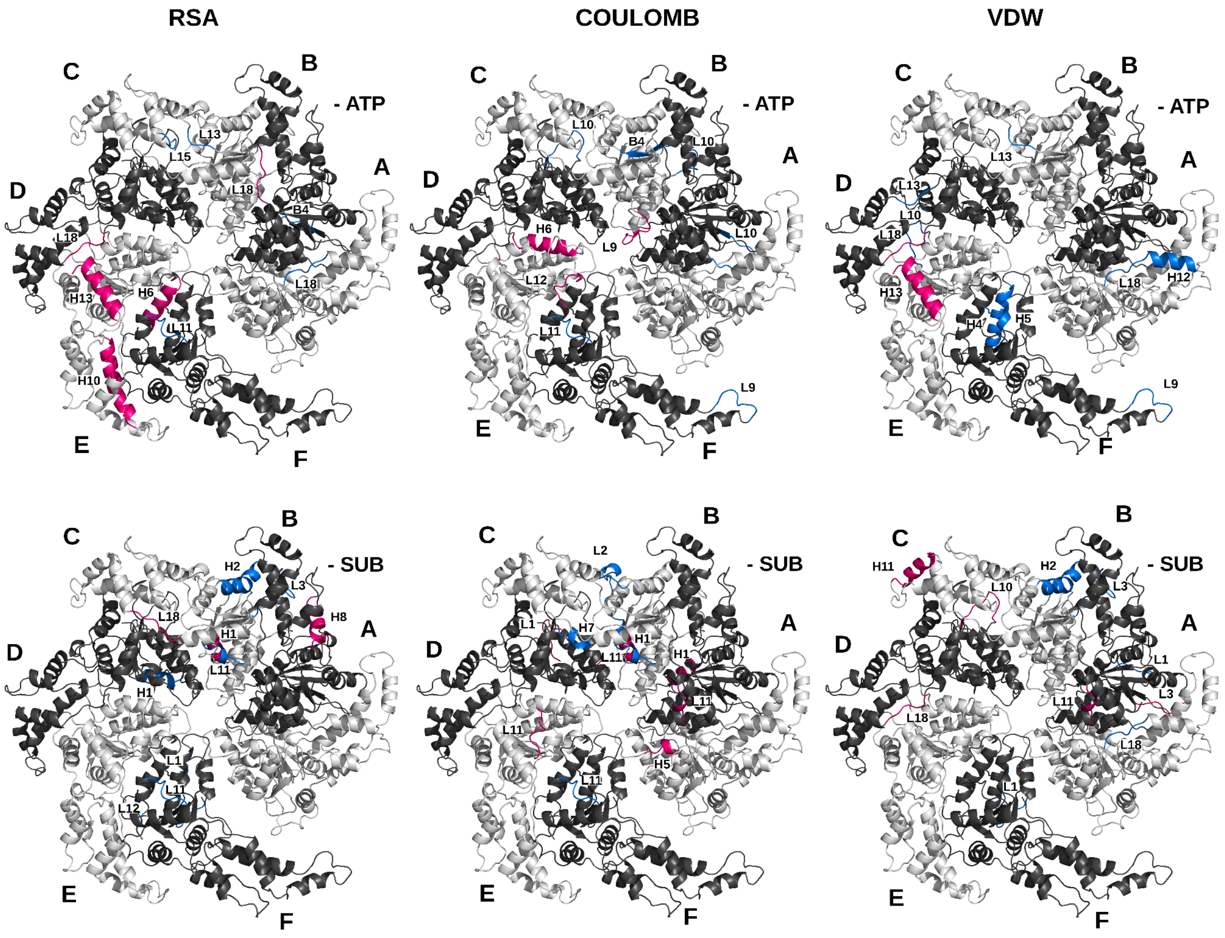


Figure S13. Key regions associated with each transition (removal of ATP or SUB) from the katanin hexamer. Regions colored in pink indicate an increase in feature values, while regions in blue indicate a decrease in feature values.

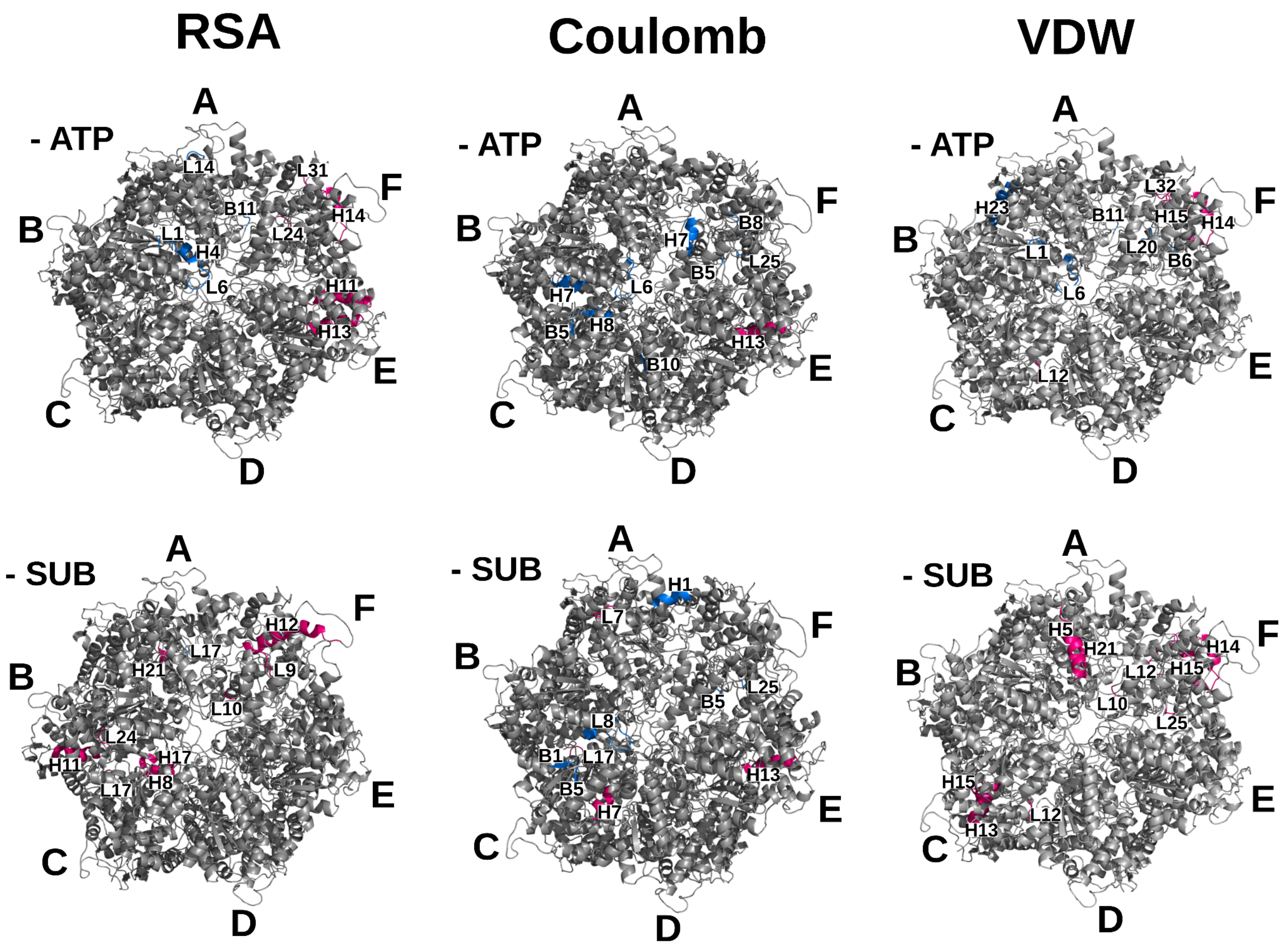


Figure S14. Key regions associated with each transition (removal of ATP or SUB) from the ClpB hexamer. Regions colored in pink indicate an increase in feature values, while regions in blue indicate a decrease in feature values.

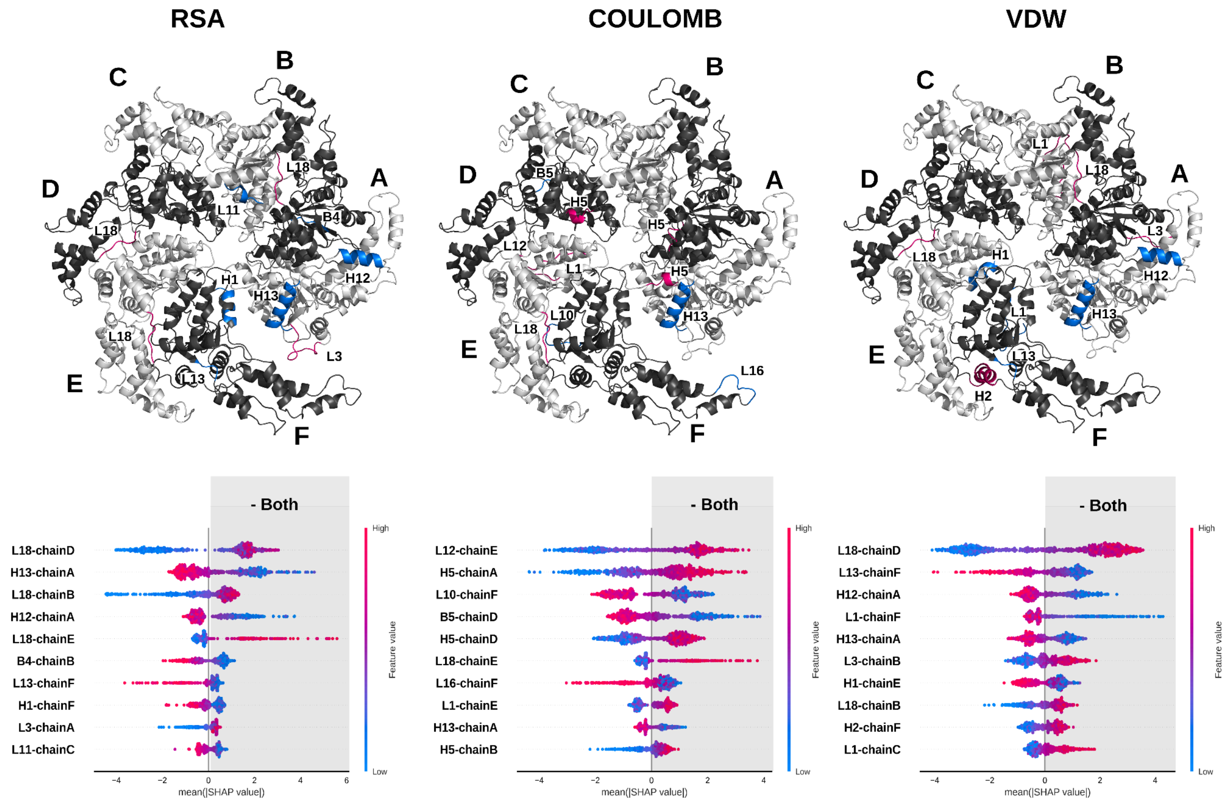


Figure S15. Key regions in the katanin hexamer associated with the removal of both ligands (SUB and ATP). Regions highlighted in pink indicate an increase in feature values, while those in blue indicate a decrease. The bottom panels display the corresponding SHAP beeswarm plots, where positive SHAP values reflect the same directional changes of these key regions.

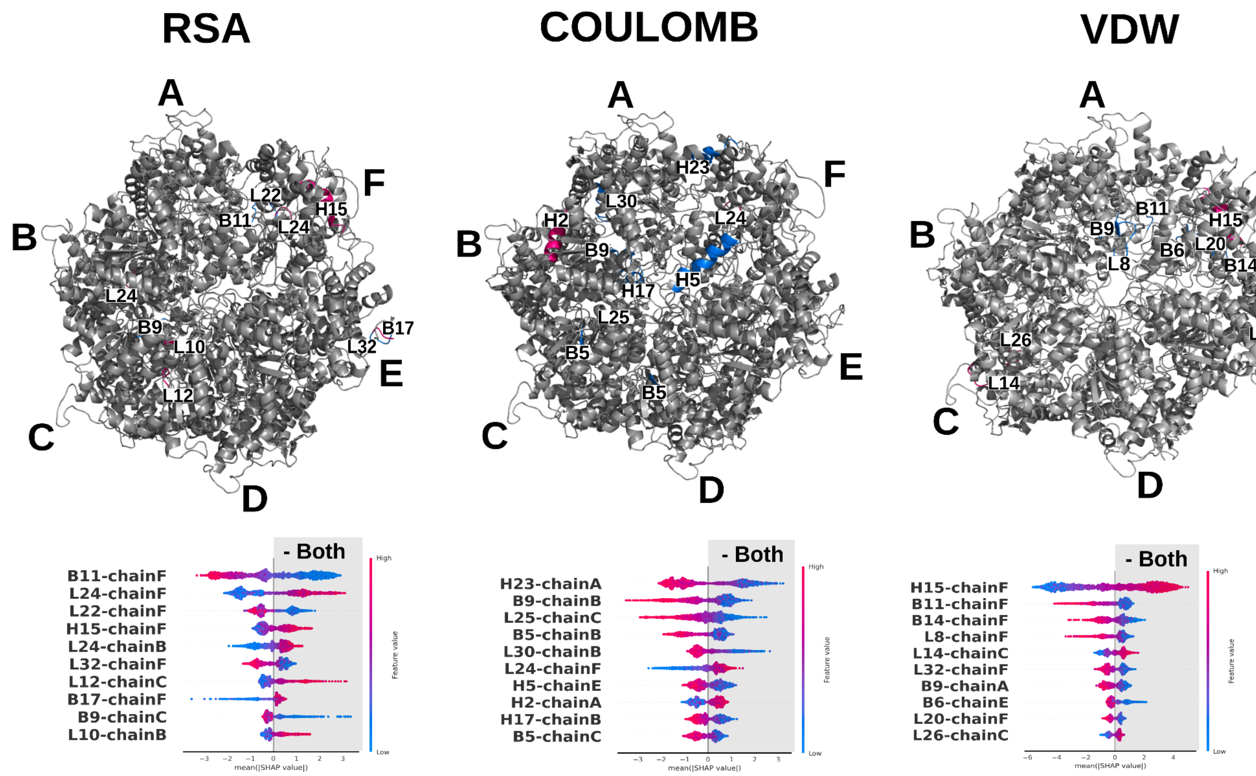


Figure S16. Key regions in the ClpB hexamer associated with the removal of both ligands (SUB and ATP). Regions highlighted in pink indicate an increase in feature values, while those in blue indicate a decrease. The bottom panels display the corresponding SHAP beeswarm plots, where positive SHAP values reflect the same directional changes of these key regions.

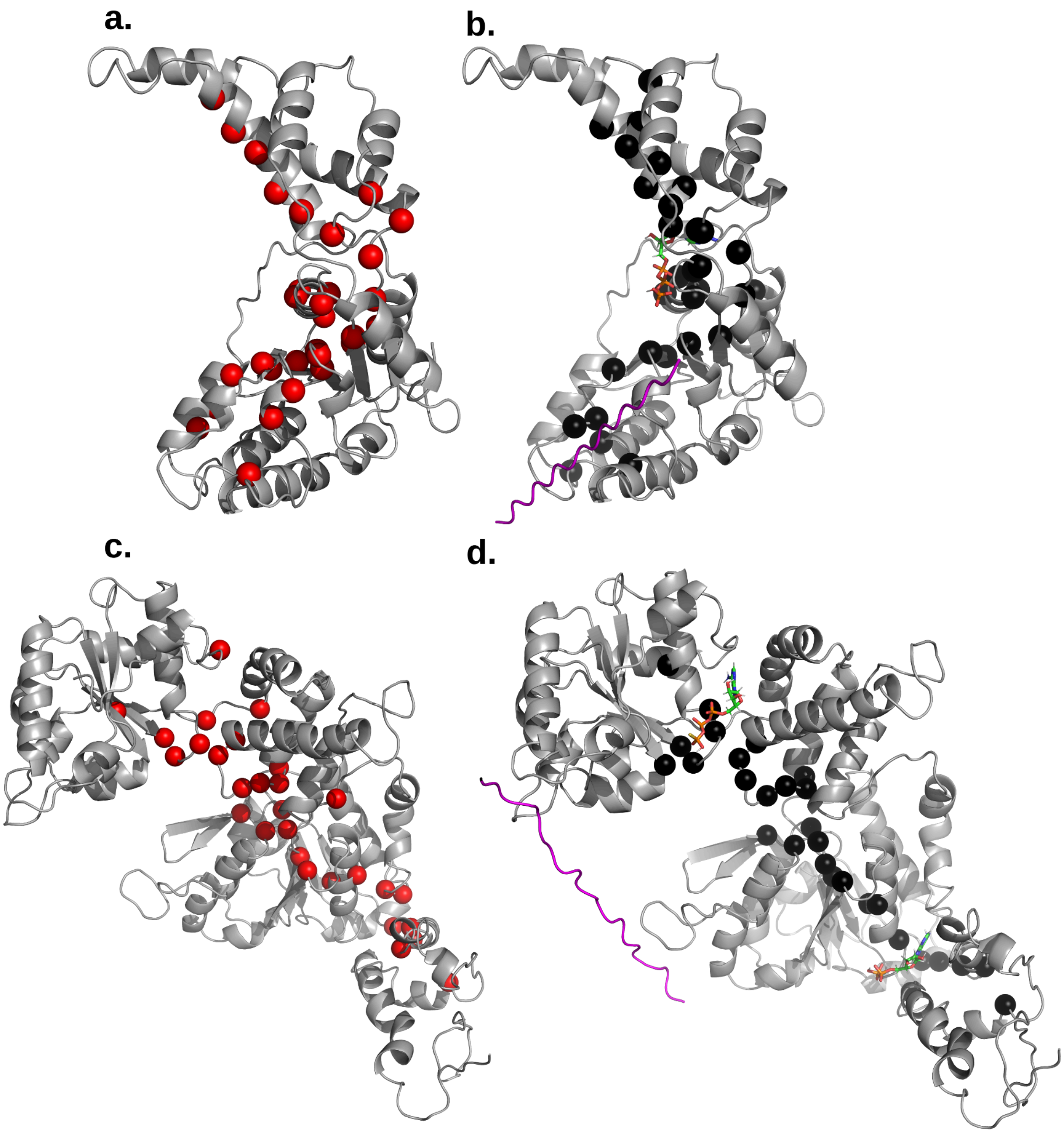


Figure S17. The top 10% betweenness centrality residues for monomeric structures of (a.) katanin APO, top left, and (b.) katanin CPX, top right (listed in Table S3). The top 5% betweenness centrality residues for monomeric structures of (c.) ClpB APO, bottom left, and (d.) ClpB CPX, bottom right (listed in Table S15). The CPX setup indicates the substrate (magenta), ATP (lines), and the central residues (black). In the APO setup the residues are colored in red.

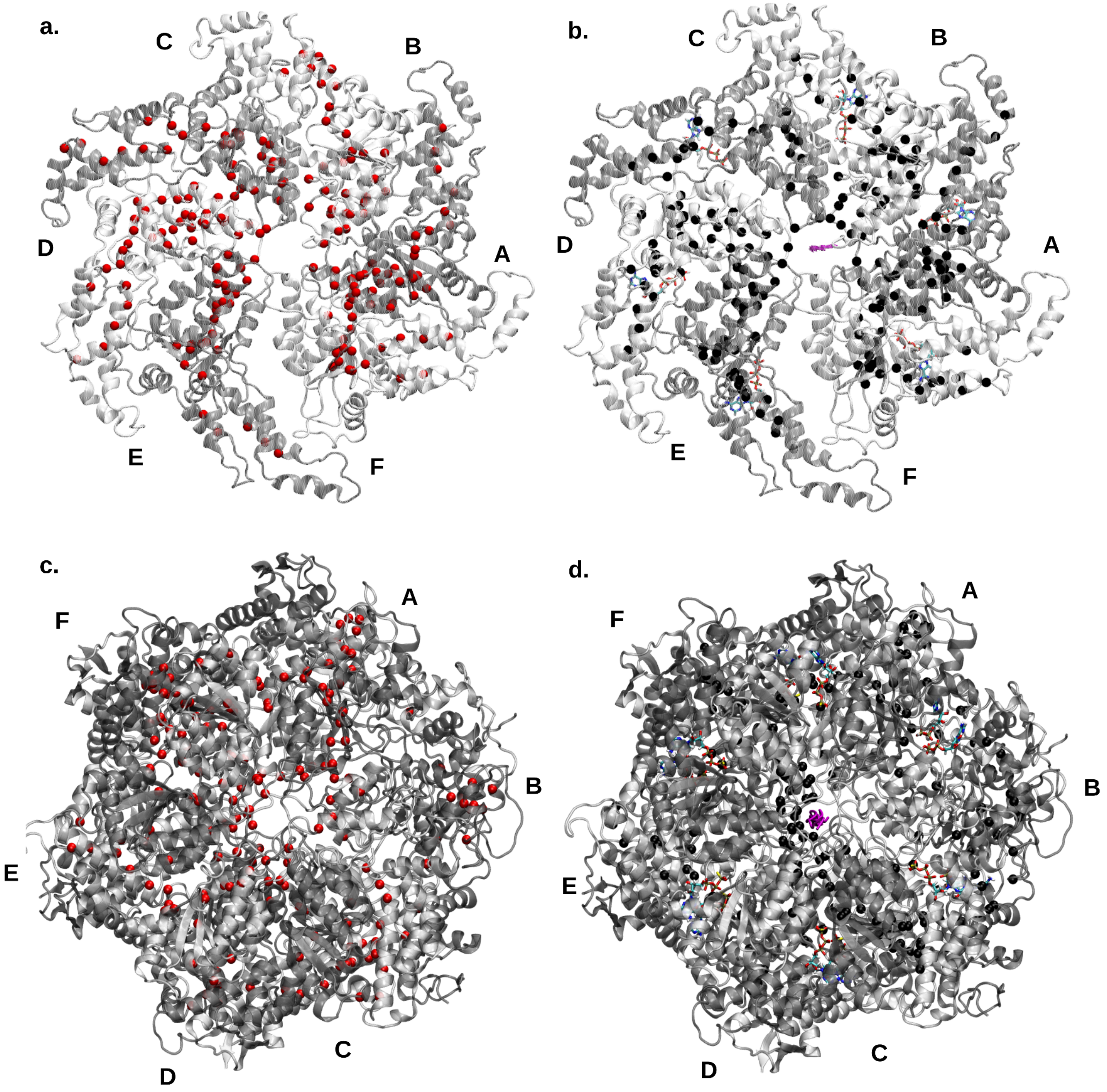


Figure S18. The positions of top 10% betweenness centrality residues for hexameric structures of (a.) Katanin APO, top left, and (b.) Katanin CPX, top right (listed in Tables S4-S5). The positions of top 5% betweenness centrality residues for hexameric structures of (c.) ClpB APO, bottom left, and (d.) ClpB CPX, bottom right (listed in Tables S16-S17). The CPX setup depicts the substrate (magenta), ATP (lines), and the central residues (black). In the APO setup the residues are colored in red.

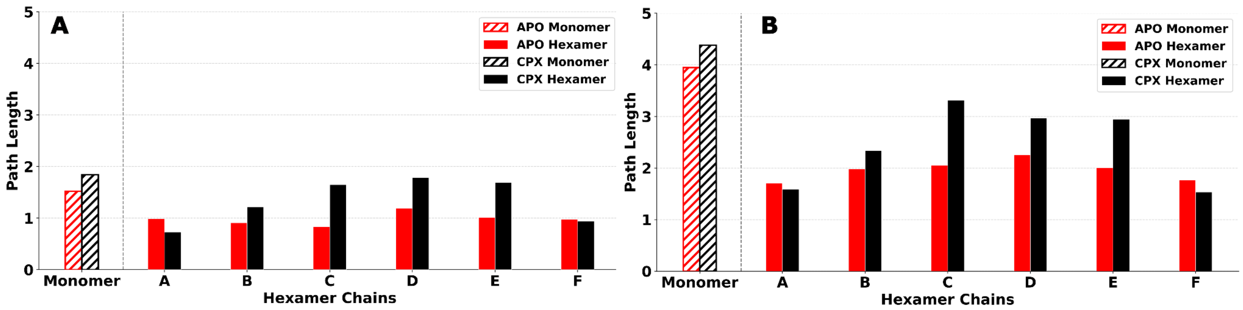


Figure S19. Katanin intra-protomer shortest path lengths in the APO and CPX setups. Shortest path lengths are shown for katanin monomer (hashed) and individual protomers (chains A-F) of the hexamer in the APO (red) and CPX (black). (A) shortest paths within the NBD, from WA to PL1, and (B) shortest paths from NBD to HBD, from PL1 to HBD tip.

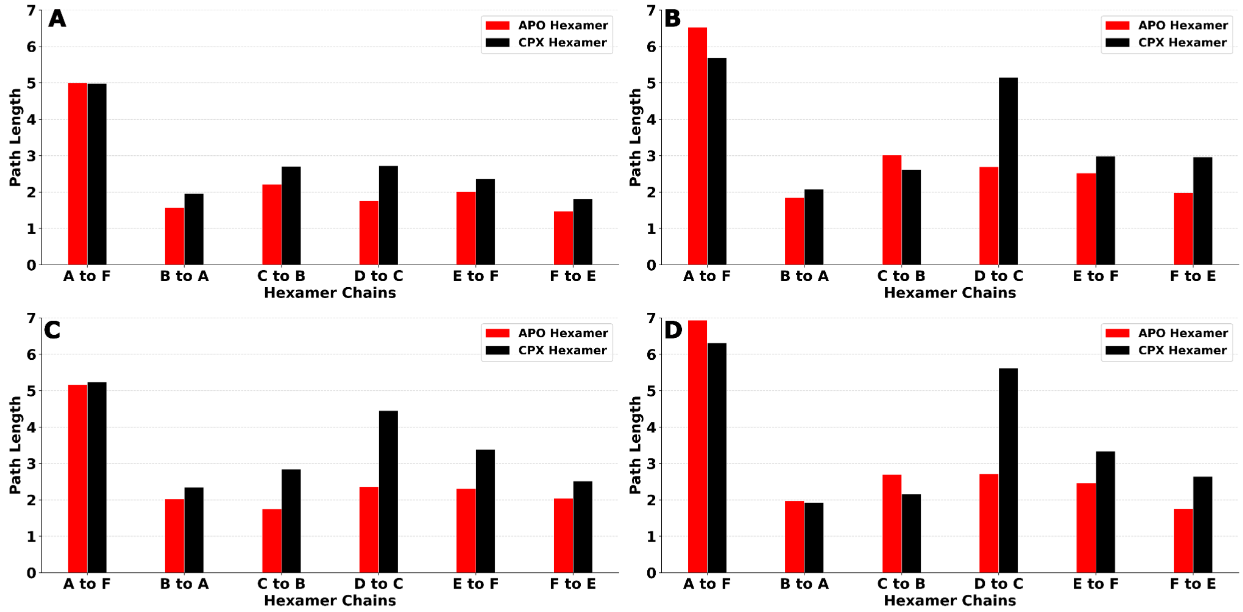


Figure S20. Katanin inter-protomer shortest path lengths in the APO and CPX setups. Shortest path lengths are shown for katanin monomer (hashed) and individual protomers (chains A-F) of the hexamer in the APO (red) and CPX (black). (A) shortest paths within the NBD, from WA to PL1, (B) shortest paths from NBD to HBD, from PL1 to HBD tip, (C) shortest paths within the NBD, from WA to PL2, and (D) shortest paths from NBD to HBD, from PL2 to HBD tip.

| Setup | Top 10% $C_B$ Katanin - Monomer |
|---|---|
| APO | V175, V178, T181, T210, V215, L231, A232, K239, I242, R244, S248, S250, F255, T256, F290, D295, L297, S310, A337, A338, T339, I359, L361, D363, R367, S397, D400, S403, T407, I410, L413 |
| CPX | Y170, K202, H206, L231, T237, G238, K239, L241, I242, R244, S258, L262, S263, I273, F290, D292, V313, A338, I359, L361, S397, G398, A399, D400, V401, S403, L404, T407, A408, I410, V438 |

Table S3. The highly central (top 10%) residues for the katanin monomer network. These positions are shown in Fig. S17.

| Chain | Top 10% $C_B$ Katanin – Hexamer APO |
|---|---|
| A | L231, A232, V257, S258, T260, D261, I273, I288, F290, I291, V335, A337, A338, I359, L361, R367, I371, S397, D400, L404, A408 |
| B | E271, K272, I273, V274, R275, L276, L277, F278, L280, A281, H307, S310, R311, V313, K314, S315, F317, L318, M321, V333, T343, D346, R352, K355, I357, I359, P360, L361, P362, A366, R367, I371, C405, A408, V412 |
| C | L231, S258, T260, L262, S264, D269, S270, E271, I273, R275, F278, I289, F290, I291, E308, R311, R312, K314, E316, F317, M321, F334, L336, A337, I359, L361, P362, A366, R367, K369, I371, E372, M375, C405 |
| D | V230, L231, A232, V257, S258, T260, D261, L262, K265, D269, K272, V274, R275, L276, L277, F278, L280, I289, F290, V313, L336, I359, L361, P362, R367, S397, V401, C405, A408, E436, V438 |
| E | A232, S263, K265, R267, G268, D269, E271, K272, V274, R275, L276, L277, F278, L280, I289, D295, V313, S315, T339, P342, L345, A348, L349, R350, F353, K355, R356, I357, F358, I359, P360, L361, P362, R367, S397, D400, L404, A408, V438 |
| F | Y170, L231, F255, G268, D269, E271, K272, I273, V274, L276, L277, I289, I291, I294, L297, R311, R312, V313, F334, L336, A338, I359, L361, R367, I371, C405, A409, V412 |

Table S4. The highly central (top 10%) residues for the katanin hexamer network of the APO setup. These positions are shown in Fig. S18.

| Chain | Top 10% $C_B$ Katanin – Hexamer CPX |
|---|---|
| **A** | V230, L231, V257, S259, T260, D261, S264, I273, L276, L277, A281, I289, I291, F334, L336, A337, I359, L361, P362, I364, R367, I371, I384, Y386, A390, C405 |
| **B** | L231, A232, G233, P235, G236, T237, K265, R267, D269, E271, K272, I273, R275, L277, F278, A281, I288, I289, S315, E316, L318, L336, R352, K355, I357, F358, I359, P362, D400, L404, E436, V438, F443 |
| **C** | Y170, I174, S224, A228, V230, L231, A232, T237, A247, T260, S263, S264, K265, W266, R267, D269, E271, K272, I273, L277, A281, I288, I289, F334, L336, R351, R352, F353, R356, I359, P362, R367, D400, L404, A408 |
| **D** | I174, L231, K265, R267, E271, K272, L276, L277, F278, L280, A281, I289, V333, L336, I359, L361, P362, F396, S397, D400, L404, A446, S451 |
| **E** | I174, V209, M229, V230, L231, P235, G238, L241, R244, A247, T260, S264, W266, D269, K272, I273, R275, L276, L277, F278, A281, S287, I294, T296, V333, V335, T339, R356, I359, P362, G398, A399, V401, L404, A408, C462, T465, F469 |
| **F** | Y170, L231, T237, G238, K239, L241, A243, V257, R267, D269, E271, K272, I273, V274, R275, L276, L277, A281, S287, I289, F290, R312, E316, F334, L336, F358, G398, D400, V401, L404, C405, A409 |

Table S5. The highly central (top 10%) residues for the katanin hexamer network of the CPX setup. These positions are shown in Fig. S18.

| Katanin - Monomer | | |
|---|---|---|
| **Setup** | **APO** | **CPX** |
| **WA to PL1** (1.52/1.84) Avg. Path (2.01/2.76) | T240 - F290 - T256 - V178 - I273 - E271 | T240 - R244 - F290 - S258 - D261 - S264 - W266 |
| **WA to PL2** (1.61/2.16) Avg. Path (2.44/2.86) | T240 - D292 - I294 - L297 - S310 - H307 | T240 - R244 - F290 - S258 - T260 - S263 - S310 - H307 |
| **PL1 to HBD tip** (3.95/4.38) Avg. Paths (4.25/4.81) | W266 - S264 - T260 - S258 - D292 - A338 - L231 - I359 - L361 - S397 - D400 - S403 - T407 - I410 - L413 - T418 | W266 - S264 - D261 - S258 - F290 - R244 - L241 - T237 - A399 - S403 - T407 - I410 - R414 - Y416 - T418 |

| | | |
|---|---|---|
| **PL2 to HBD tip** (3.81/4.70) Avg. Paths (4.18/5.04) | H307 - S310 - L297 - D295 - T339 - A232 - I359 - L361 - S397 - D400 - S403 - T407 - I410 - L413 - T418 | H307 - S310 - S263 - T260 - S258 - F290 - R244 - L241 - T237 - A399 - S403 - T407 - I410 - R414 - Y416 - T418 |

Table S6. Optimal paths for the katanin monomer for the ATP binding pocket (Walker A - P235/T240) to the CTT binding channel (PL1 - W266/E271, PL2 - H307/R312) and from the CTT binding channel (PL1 - W266/E271, PL2 - H307/R312) to the HBD tip (T418/L426). The path lengths and average lengths from suboptimal paths are provided under the setup name in order of (APO/CPX).

| **Katanin – Hexamer Intra-Protomer Paths – Walker A to PL1** | | |
|---|---|---|
| **Setup** | **APO** | **CPX** |
| **Chain A** (0.99/0.73) Avg. Path (1.20/0.92) | A:T240 – A:F290 – A:F255 – A:L277 – A:I273 – A:E271 | A:T240 – A:A243 – A:I289 – A:F278 – A:R275 – A:E271 |
| **Chain B** (0.91/1.22) Avg. Path (1.19/1.55) | B:T240 – B:A243 – B:I289 – B:L277 – B:V274 – B:E271 | B:T240 – B:A243 – B:I289 – B:F278 – B:R275 – B:E271 |
| **Chain C** (0.83/1.65) Avg. Paths (1.13/2.06) | C:T240 – C:A243 – C:I289 – C:F278 – C:V274 – C:E271 | C:T240 – C:R244 – C:A247 – C:I288 – C:A281 – C:L277 – C:V274 – C:E271 |
| **Chain D** (1.19/1.79) Avg. Paths (1.59/2.17) | D:T240 – D:A243 – D:I289 – D:L277 – D:I273 – D:E271 | D:T240 – D:R244 – D:A247 – D:F255 – D:L277 – D:I273 – D:E271 |
| **Chain E** (1.01/1.69) Avg. Paths (1.32/2.01) | E:T240 – E:A243 – E:I289 – E:L277 – E:V274 – E:E271 | E:T240 – E:R244 – E:A247 – E:S287 – E:A281 – E:L277 – E:V274 – E:E271 |
| **Chain F** (0.98/0.94) Avg. Paths (1.34/1.21) | F:T240 – F:F290 – F:V257 – F:V274 – F:E271 | F:T240 – F:F290 – F:V257 – F:V274 – F:E271 |

Table S7. Intra-protomer optimal paths for the katanin hexamer for the ATP binding pocket (Walker A - P235/T240) to the CTT binding channel (PL1 - W266/E271). The residues in the paths are indicated as (chain):(amino acid)(residue ID). The path lengths and average lengths from suboptimal paths are provided under the chain names in order of (APO/CPX).

| **Katanin – Hexamer Intra-Protomer Paths – Walker A to PL2** | | |
|---|---|---|
| **Setup** | **APO** | **CPX** |

| | | |
|---|---|---|
| **Chain A** (0.96/0.88) Avg. Path (1.21/1.07) | A:P235 – A:A232 – A:V230 – A:F353 – A:L349 – A:K314 – A:R312 | A:T240 – A:A243 – A:F290 – A:V335 – A:M321 – A:V319 – A:E316 – A:R312 |
| **Chain B** (1.06/1.55) Avg. Path (1.35/1.86) | B:P235 – B:N340 – B:E344 – B:D346 – B:K314 – B:R312 | B:T240 – B:A243 – B:I289 – B:F278 – B:R275 – B:E316 – B:R312 |
| **Chain C** (1.09/1.90) Avg. Paths (1.36/2.27) | C:T240 – C:A243 – C:I289 – C:F278 – C:R275 – C:E316 – C:R312 | C:P235 – C:G233 – C:L231 – C:M229 – C:R352 – C:L318 – C:S315 – C:R312 |
| **Chain D** (1.55/1.93) Avg. Paths (1.93/2.47) | D:T240 – D:A243 – D:I289 – D:F278 – D:R275 – D:E316 – D:R312 | D:P235 – D:A232 – D:V230 – D:R352 – D:L318 – D:S315 – D:R312 |
| **Chain E** (1.08/2.04) Avg. Paths (1.38/2.33) | E:P235 – E:N340 – E:E344 – E:D346 – E:K314 – E:R312 | E:T240 – E:R244 – E:A247 – E:S287 – E:A281 – E:L277 – E:V274 – E:E271 – E:R312 |
| **Chain F** (1.20/1.26) Avg. Paths (1.52/1.49) | F:T240 – F:F290 – F:V257 – F:V274 – F:E271 – F:R312 | F:T240 – F:F290 – F:V257 – F:V274 – F:E271 – F:R312 |

Table S8. Intra-protomer optimal paths for the katanin hexamer for ATP binding pocket (Walker A - P235/T240) to the CTT binding channel (PL2 - H307/R312). The residues in the paths are indicated as (chain):(amino acid)(residue ID). The path lengths and average lengths from suboptimal paths are provided under the chain names in order of (APO/CPX).

| **Katanin – Hexamer Intra-Protomer Paths – PL1 to HBD tip** | | |
|---|---|---|
| **Setup** | **APO** | **CPX** |
| **Chain A** (1.70/1.59) Avg. Path (1.78/1.67) | A:E271 – A:I273 – A:L277 – A:I289 – A:L336 – A:L231 – A:I359 – A:L361 – A:S397 – A:D400 – A:L404 – A:A408 – A:N411 – A:Y416 – A:T418 | A:E271 – A:R275 – A:F278 – A:I289 – A:L336 – A:L231 – A:I359 – A:P362 – A:R367 – A:I371 – A:M375 – A:T378 – A:K434 – A:M430 – A:L426 |
| **Chain B** (1.98/2.34) Avg. Path (2.07/2.46) | B:E271 – B:V274 – B:L277 – B:I289 – B:L336 – B:L231 – B:I359 – B:L361 – B:D363 – B:D365 – B:K368 – B:I371 – B:C405 – B:A408 – B:V412 – B:L433 – B:A429 – B:L426 | B:E271 – B:R275 – B:F278 – B:I289 – B:L336 – B:L231 – B:I359 – B:T237 – B:D400 – B:L404 – B:F443 – B:V438 – B:E436 – B:L433 – B:M430 – B:L426 |
| **Chain C** (2.05/3.31) Avg. Paths (2.18/3.44) | C:E271 – C:V274 – C:F278 – C:I289 – C:L336 – C:L231 – C:I359 – C:P362 – C:A366 – C:K369 – C:E372 – C:M375 – C:P379 – C:K434 – C:M430 – C:L426 | C:E271 – C:R275 – C:F278 – C:I289 – C:L336 – C:V230 – C:A232 – C:T237 – C:D400 – C:L404 – C:A408 – C:L413 – C:T418 |

| | | |
|---|---|---|
| **Chain D** (2.25/2.96) Avg. Paths (2.39/3.11) | D:E271 – D:I273 – D:L277 – D:I289 – D:L336 – D:L231 – D:I359 – D:L361 – D:S397 – D:V401 – D:C405 – D:A408 – D:V438 – D:E436 – D:L433 – D:M430 – D:L426 | D:E271 – D:I273 – D:L277 – D:I289 – D:L336 – D:L231 – D:I359 – D:L361 – D:S397 – D:V401 – D:C405 – D:A409 – D:V412 – D:L433 - D:A429 – D:L426 |
| **Chain E** (2.00/2.95) Avg. Paths (2.09/3.07) | E:E271 – E:V274 – E:L277 – E:I289 – E:L336 – E:L231 – E:I359 – E:L361 – E:S397 – E:D400 – E:L404 – E:A408 – E:V438 – E:E436 – E:L433 – E:M430 – E:L426 | E:E271 – E:V274 – E:L277 – E:A281 – E:S287 – E:A247 – E:R244 – E:L241 – E:G238 – E:G398 – E:V401 – E:L404 – E:A408 – E:V412 – E:L433 – E:M430 – E:L426 |
| **Chain F** (1.76/1.54) Avg. Paths (1.88/1.63) | F:E271 – F:I273 – F:L277 – F:I289 – F:L336 – F:L231 – F:I359 – F:L361 – F:R367 – F:I371 – F:C405 – F:A409 – F:L413 – F:T418 | F:E271 – F:V274 – F:V257 – F:F290 – F:T240 – F:G238 – F:G398 – F:V402 – F:R406 – F:I410 – F:R414 – F:Y416 – F:T418 |

Table S9. Intra-protomer optimal paths for the katanin hexamer from the CTT binding channel (PL1 - W266/E271) to the HBD tip (T418/L426). The residues in the paths are indicated as (chain):(amino acid)(residue ID). The path lengths and average lengths from suboptimal paths are provided under the chain names in order of (APO/CPX).

| **Katanin – Hexamer Intra-Protomer Paths – PL2 to HBD tip** | | |
|---|---|---|
| **Setup** | **APO** | **CPX** |
| **Chain A** (1.69/1.62) Avg. Path (1.78/1.70) | A:R312 – A:K314 – A:L349 – A:F353 – A:V230 – A:A232 – A:I359 – A:L361 – A:S397 – A:D400 – A:L404 – A:A408 – A:N411 – A:Y416 – A:T418 | A:R312 – A:S315 – A:L318 – A:R352 – A:V230 – A:I357 – A:I359 – A:P362 – A:R367 – A:I371 – A:M375 – A:T378 – A:K434 – A:M430 – A:L426 |
| **Chain B** (1.85/2.47) Avg. Path (1.94/2.61) | B:R312 – B:S315 – B:L318 – B:R352 – B:K355 – B:I357 – B:P360 – B:P362 – B:R367 – B:I371 – B:C405 – B:A408 – B:V412 – B:L433 – B:A429 – B:L426 | B:R312 – B:S315 – B:L318 – B:R352 – B:K355 – B:I357 – B:I359 – B:T237 – B:D400 – B:L404 – B:F443 – B:V438 – B:E436 – B:L433 – B:M430 – B:L426 |
| **Chain C** (2.32/3.33) Avg. Paths (2.43/3.49) | C:R312 – C:E316 – C:R275 – C:F278 – C:I289 – C:L336 – C:L231 – C:I359 – C:P362 – C:A366 – C:K369 – C:E372 – C:M375 – C:P379 – C:K434 – C:M430 – C:L426 | C:R312 – C:S315 – C:L318 – C:R352 – C:K355 – C:I357 – C:I359 – C:P362 – C:R367 – C:V401 – C:L404 – C:A408 – C:L413 – C:T418 |

| | | |
|---|---|---|
| **Chain D**<br>(2.61/2.92)<br>Avg. Paths (2.73/3.12) | D:R312 – D:E316 – D:R275 – D:F278 – D:I289 – D:L336 – D:L231 – D:I359 – D:L361 – D:S397 – D:V401 – D:C405 – D:A408 – D:V438 – D:E436 – D:L433 – D:M430 – D:L426 | D:R312 – D:S315 – D:L318 – D:R352 – D:K355 – D:I357 – D:I359 – D:L361 – D:S397 – D:V401 – D:C405 – D:A409 – D:V412 – D:L433 – D:A429 – D:L426 |
| **Chain E**<br>(1.87/3.30)<br>Avg. Paths (1.95/3.42) | E:R312 – E:K314 – E:D346 – E:P342 – E:A232 – E:I359 – E:L361 – E:S397 – E:D400 – E:L404 – E:A408 – E:V438 – E:E436 – E:L433 – E:M430 – E:L426 | E:R312 – E:E271 – E:V274 – E:L277 – E:A281 – E:S287 – E:A247 – E:R244 – E:L241 – E:G238 – E:G398 – E:V401 – E:L404 – E:A408 – E:V412 – E:L433 – E:M430 – E:L426 |
| **Chain F**<br>(1.92/1.86)<br>Avg. Paths (2.04/1.94) | F:R312 – F:K314 – F:L297 – F:I294 – F:A338 – F:L231 – F:I359 – F:L361 – F:R367 – F:I371 – F:C405 – F:A409 – F:L413 – F:T418 | F:R312 – F:E271 – F:V274 – F:V257 – F:F290 – F:T240 – F:G238 – F:G398 – F:V402 – F:R406 – F:I410 – F:R414 – F:Y416 – F:T418 |

Table S10. Intra-protomer optimal paths for the katanin hexamer from the CTT binding channel (PL2 - H307/R312) to the HBD tip (T418/L426). The residues in the paths are indicated as (chain):(amino acid)(residue ID). The path lengths and average lengths from suboptimal paths are provided under the chain names in order of (APO/CPX).

| **Katanin – Hexamer Inter-Protomer Paths – WA to PL1** | | |
|---|---|---|
| **Setup** | **APO** | **CPX** |
| **A to F**<br>(4.99/4.99)<br>Avg. Path (5.27/5.17) | A:T240 – A:F290 – A:S258 – A:T260 – B:V313 – B:S310 – B:H307 – C:E308 – C:R312 – C:S270 – C:S264 – D:D269 – D:K265 – E:R267 – E:K265 – F:W266 | A:T240 – A:A243 – A:F290 – A:V257 – A:L262 – A:S264 – B:R267 – B:K265 – C:W266 – C:S264 – C:T260 – D:E271 – D:R267 – D:K265 – E:R267 – E:K265 – F:W266 |
| **B to A**<br>(1.58/1.97)<br>Avg. Path (1.85/2.23) | B:T240 – B:A243 – B:I289 – B:L277 – B:K272 – A:S264 – A:W266 | B:T240 – B:R244 – B:A247 – B:I288 – B:A281 – B:L277 – B:I273 – B:D269 – A:S264 – A:W266 |
| **C to B**<br>(2.21/2.71)<br>Avg. Paths (2.38/2.89) | C:T240 – C:A243 – C:I289 – C:F278 – C:R275 – C:E316 – C:R312 – C:E308 – B:H307 – B:S310 – B:V313 – B:E271 | C:P235 – C:G233 – C:F358 – C:R356 – C:R351 – B:P235 – B:A232 – B:V230 – B:L336 – B:I289 – B:F278 – B:R275 – B:E271 |
| **D to C**<br>(1.76/2.73)<br>Avg. Paths (2.16/3.09) | D:T240 – D:A243 – D:I289 – D:L277 – D:K272 – C:S264 – C:W266 | D:T240 – D:R244 – D:A247 – D:T253 – D:S287 – D:L280 – D:L276 – D:K272 – C:T260 – C:S264 – C:W266 |

| | | |
|---|---|---|
| **E to D**<br>(2.01/2.36)<br>Avg. Paths (2.26/2.64) | E:P235 – E:N340 – E:P342 – E:L345 – E:L349 – E:S315 – D:T260 – D:S264 – D:W266 | E:T240 – E:R244 – E:A247 – E:S287 – E:A281 – E:L277 – E:I273 – E:D269 – E:R267 – D:W266 |
| **F to E**<br>(1.47/1.81)<br>Avg. Paths (1.77/2.03) | F:T240 – F:F290 – F:F255 – F:L277 – F:I273 – F:G268 – E:W266 | F:T240 – F:F290 – F:T256 – F:L277 – F:I273 – F:D269 – F:R267 – E:W266 |

Table S11. Inter-protomer optimal paths for the katanin hexamer for the ATP binding pocket (Walker A - P235/T240) to the CTT binding channel (PL1 - W266/E271). The residues in the paths are indicated as (chain):(amino acid)(residue ID). The path lengths and average lengths from suboptimal paths are provided under the chain names in order of (APO/CPX).

| **Katanin – Hexamer Inter-Protomer Paths – WA to PL2** | | |
|---|---|---|
| **Setup** | **APO** | **CPX** |
| **A to F**<br>(5.17/5.24)<br>Avg. Path (5.37/5.42) | A:T240 – A:F290 – A:S258 – A:T260 – B:V313 – B:S310 – B:H307 – C:E308 – C:R312 – C:S270 – C:S264 – D:D269 – D:K265 – E:R267 – E:K265 – E:S263 – F:R312 | A:T240 – A:A243 – A:F290 – A:V257 – A:L262 – A:S264 – B:R267 – B:K265 – C:W266 – C:S264 – C:T260 – D:E271 – D:R267 – D:K265 – E:W266 – E:S264 – E:T260 – F:R312 |
| **B to A**<br>(2.02/2.35)<br>Avg. Path (2.22/2.58) | B:T240 – B:A243 – B:I289 – B:L277 – B:K272 – A:S264 – A:E271 – A:R312 | B:T240 – B:A243 – B:I289 – B:F278 – B:R275 – B:K272 – A:D261 – A:V274 – A:E316 – A:R312 |
| **C to B**<br>(1.75/2.84)<br>Avg. Paths (1.97/3.07) | C:T240 – C:A243 – C:I289 – C:F278 – C:R275 – C:E316 – C:R312 – C:E308 – B:H307 | C:P235 – C:G233 – C:F358 – C:R356 – C:R351 – B:P235 – B:G233 – B:F358 – B:K355 – B:R352 – B:L318 – B:S315 – B:R312 |
| **D to C**<br>(2.36/4.45)<br>Avg. Paths (2.69/4.71) | D:T240 – D:A243 – D:I289 – D:L277 – D:K272 – C:S264 – C:S270 – C:R312 | D:T240 – D:R244 – D:A247 – D:T253 – D:S287 – D:L280 – D:L276 – D:K272 – C:T260 – C:S263 – C:K265 – C:R267 – C:E271 – C:V313 – C:S315 – C:R312 |
| **E to D**<br>(2.31/3.39)<br>Avg. Paths (2.57/3.69) | E:P235 – E:N340 – E:P342 – E:L345 – E:L349 – E:S315 – D:T260 – D:S263 – D:S310 – D:H307 | E:T240 – E:R244 – E:A247 – E:S287 – E:A281 – E:L277 – E:I273 – E:D269 – D:K265 – D:R267 – D:E271 – D:R312 |
| **F to E**<br>(2.04/2.52)<br>Avg. Paths (2.27/2.72) | F:T240 – F:F290 – F:V257 – F:V274 – F:E271 – E:S263 – E:V313 – E:S315 – E:R312 | F:T240 – F:F290 – F:T256 – F:L277 – F:I273 – F:D269 – F:R267 – E:W266 – E:G268 – E:R312 |

Table S12. Inter-protomer optimal paths for katanin hexamer for the ATP binding pocket (Walker A - P235/T240) to the CTT binding channel (PL2 - H307/R312). The residues in the paths are indicated as (chain):(amino acid)(residue ID). The path lengths and average lengths from suboptimal paths are provided under the chain names in order of (APO/CPX).

| Katanin – Hexamer Inter-Protomer Paths – PL1 to HBD tip | | |
|---|---|---|
| **Setup** | **APO** | **CPX** |
| **A to F** (6.53/5.69) Avg. Path (6.65/5.77) | A:W266 – A:S264 – A:T260 – B:V313 – B:S310 – B:H307 – C:E308 – C:R312 – C:S270 – C:S264 – D:D269 – D:K265 – E:R267 – E:K265 – F:D269 – F:I273 – F:L277 – F:I289 – F:L336 – F:L231 – F:I359 – F:L361 – F:R367 – F:I371 – F:C405 – F:A409 – F:L413 – F:T418 | A:W266 – B:R267 – B:K265 – C:W266 – C:S264 – C:T260 – D:E271 – D:R267 – D:K265 – E:W266 – F:R267 – F:D269 – F:I273 – F:L277 – F:I289 – F:A243 – F:L241 – F:G238 – F:G398 – F:V402 – F:R406 – F:I410 – F:R414 – F:Y416 – F:T418 |
| **B to A** (1.85/2.07) Avg. Path (1.95/2.17) | B:E271 – A:T260 – A:S258 – A:I291 – A:A337 – A:L231 – A:I359 – A:L361 – A:S397 – A:D400 – A:L404 – A:A408 – A:N411 – A:Y416 – A:T418 | B:E271 – A:T260 – A:V257 – A:I291 – A:A337 – A:L231 – A:I359 - A362 – A:R367 – A:I371 – A:M375 – A:T378 – A:K434 – A:M430 – A:L426 |
| **C to B** (3.01/2.62) Avg. Paths (3.09/2.73) | C:E271 – C:R312 – C:E308 – B:H307 – B:R311 – B:S315 – B:L318 – B:R352 – B:K355 – B:I357 – B:P360 – B:P362 – B:R367 – B:I371 – B:C405 – B:A408 – B:V412 – B:L433 – B:A429 – B:L426 | C:E271 – C:R275 – C:F278 – C:V333 – C:A228 – C:R352 – B:G236 – B:D400 – B:L404 – B:F443 – B:V438 – B:E436 – B:L433 – B:M430 – B:L426 |
| **D to C** (2.69/5.16) Avg. Paths (2.83/5.27) | D:E271 – C:T260 – C:S258 – C:I291 – C:A337 – C:L231 – C:I359 – C:P362 – C:A366 – C:K369 – C:E372 – C:M375 – C:P379 – C:K434 – C:M430 – C:L426 | D:E271 – C:T260 – C:S263 – C:K265 – C:R267 – C:D269 – C:I273 – C:L277 – C:I289 – C:L336 – C:V230 – C:A232 – C:T237 – C:D400 – C:L404 – C:A408 – C:L413 – C:T418 |
| **E to D** (2.52/2.99) Avg. Paths (2.64/3.17) | E:E271 – D:D261 – D:V257 - D290 – D:L336 – D:L231 – D:I359 – D:L361 – D:S397 – D:V401 – D:C405 – D:A408 – D:V438 – D:E436 – D:L433 – D:M430 – D:L426 | E:E271 – E:V274 – E:F278 – E:V333 – E:V335 – E:M229 – E:R356 – E:F469 – D:S451 – D:A446 – D:D442 – D:V438 – D:E436 – D:L433 – D:A429 – D:L426 |
| **F to E** (1.98/2.96) Avg. Paths (2.07/3.14) | F:E271 – F:R312 – E:T296 – E:T339 – E:A232 – E:I359 – E:L361 – E:S397 – E:D400 – E:L404 – E:A408 – E:V438 – E:E436 – E:L433 – E:M430 – E:L426 | F:E271 – F:R312 – E:T296 – E:I294 – E:T339 – E:P235 – E:A399 – E:V402 – E:C405 – E:A408 – E:V412 – E:L433 – E:M430 – E:L426 |

Table S13. Inter-protomer optimal paths for the katanin hexamer from the CTT binding channel (PL1 - W266/E271) to the HBD tip (T418/L426). The residues in the paths are indicated as (chain):(amino acid)(residue ID). The path lengths and average lengths from suboptimal paths are provided under the chain names in order of (APO/CPX).

| Katanin – Hexamer Inter-Protomer Paths – PL2 to HBD tip | | |
|---|---|---|
| **Setup** | **APO** | **CPX** |
| **A to F**<br>(6.94/6.32)<br>Avg. Path (7.05/6.39) | A:R312 – A:E316 – A:I273 – A:D261 – B:E271 – B:V313 – B:S310 – B:H307 – C:E308 – C:R312 – C:S270 – C:S264 – D:D269 – D:K265 – E:R267 – E:K265 – F:D269 – F:I273 – F:L277 – F:I289 – F:L336 – F:L231 – F:I359 – F:L361 – F:R367 – F:I371 – F:C405 – F:A409 – F:L413 – F:T418 | A:R312 – A:E271 – A:S264 – B:R267 – B:K265 – C:W266 – C:S264 – C:T260 – D:E271 – D:R267 – D:K265 – E:W266 – F:R267 – F:D269 – F:I273 – F:L277 – F:I289 – F:A243 – F:L241 – F:G238 – F:G398 – F:V402 – F:R406 – F:I410 – F:R414 – F:Y416 – F:T418 |
| **B to A**<br>(1.98/1.93)<br>Avg. Path (2.07/2.02) | B:R312 – A:T260 – A:S258 – A:I291 – A:A337 – A:L231 – A:I359 – A:L361 – A:S397 – A:D400 – A:L404 – A:A408 – A:N411 – A:Y416 - A:T418 | B:R312 – A:S259 – A:I291 – A:A337 – A:L231 – A:I359 – A:P362 – A:R367 – A:I371 – A:M375 – A:T378 – A:K434 – A:M430 – A:L426 |
| **C to B**<br>(2.69/2.17)<br>Avg. Paths (2.78/2.34) | C:H307 – B:E306 – B:R311 – B:S315 – B:L318 – B:R352 – B:K355 – B:I357 – B:P360 – B:P362 – B:R367 – B:I371 – B:C405 – B:A408 – B:V412 – B:L433 – B:A429 – B:L426 | C:R312 – C:S315 – C:L318 – C:R352 – B:E236 – B:D400 – B:L404 – B:F443 – B:V438 – B:E436 – B:L433 – B:M430 – B:L426 |
| **D to C**<br>(2.72/5.62)<br>Avg. Paths (2.85/5.73) | D:R312 – C:T260 – C:S258 – C:I291 – C:A337 – C:L231 – C:I359 – C:P362 – C:A366 – C:K369 – C:E372 – C:M375 – C:P379 – C:K434 – C:M430 – C:L426 | D:R312 – D:E271 – C:T260 – C:S263 – C:K265 – C:R267 – C:D269 – C:I273 – C:L277 – C:I289 – C:L336 – C:V230 – C:A232 – C:T237 – C:D400 – C:L404 – C:A408 – C:L413 – C:T418 |
| **E to D**<br>(2.46/3.34)<br>Avg. Paths (2.59/3.49) | E:R312 – D:T260 – D:V257 – D:F290 – D:L336 – D:L231 – D:I359 – D:L361 – D:S397 – D:V401 – D:C405 – D:A408 – D:V438 – D:E436 – D:L433 – D:M430 – D:L426 | E:R312 – E:E316 – E:R275 – E:F278 – E:V333 – E:V335 – E:M229 – E:R356 – E:F469 – D:S451 – D:A446 – D:D442 – D:V438 – D:E436 – D:L433 – D:A429 – D:L426 |
| **F to E**<br>(1.76/2.64)<br>Avg. Paths (1.85/2.82) | F:R312 – E:T296 – E:T339 – E:A232 – E:I359 – E:L361 – E:S397 – E:D400 – E:L404 – E:A408 – E:V438 – E:E436 – E:L433 – E:M430 – E:L426 | F:R312 – E:T296 – E:I294 – E:T339 – E:P235 – E:A399 – E:V402 – E:C405 – E:A408 – E:V412 – E:L433 – E:M430 – E:L426 |

Table S14. Intra-protomer optimal paths for the katanin hexamer from the CTT binding channel (PL2 - H307/R312) to the HBD tip (T418/L426). The residues in the paths are indicated as (chain):(amino acid)(residue ID). The path lengths and average lengths from suboptimal paths are provided under the chain names in order of (APO/CPX).

| Setup | Top 5% $C_B$ ClpB - Monomer |
|---|---|
| APO | Q385, G206, P670, A382, I381, I774, N711, V570, V673, D708, I771, V203, Q778, I181, R379, L800, E207, A617, A628, C615, E340, V343, Y380, G209, I571, L766, Y671, L386, P547 |
| CPX | P670, L766, A382, Q385, G610, I381, G211, C615, V673, G206, L386, D708, A772, N711, K616, P765, A628, E207, Y671, R384, L800, I771, R379, Y380, V210, S672, T315, V216, G813 |

Table S15. The highly central (top 5%) residues for the ClpB monomer network. These positions are shown in Fig. S17.

| Chain | Top 5% $C_B$ ClpB – Hexamer<br>APO |
|---|---|
| A | V548, E555, Y656, D388, M562, L540, Q334, V210, E396, P651, S400, E538, G211, P208, L204, Q692, I205, P650, L618, E658, D392, R331, R183, L559, R188, A534, L817, S672 |
| B | Y671, L540, R331, I546, V548, P547, D392, K335, A665, A649, G206, Q385, Q692, S587, G655, G652, G660, P650, I536, F709, V654, G702, A537, R588, S377, V666, I674, E396, G659, M242, S400, I381, D677, R645, L646, T700, A397, Y380, D388, T544, V210, Q334 |
| C | L540, R331, I546, V548, P547, D392, K335, A665, A649, G206, Q385, Q692, S587, G655, G652, G660, P650, I536, F709, V654, G702, A537, R588, S377, V666, I674, E396, G659, M242, S400, I381, D677, R645, L646, T700, A397, Y380, D388, T544, V210, Q334 |
| D | M551, V548, R588, V539, I598, A589, S549, R252, M638, L376, R332, E636, S644, V343, K640, R189, E340, Q334, A583, T700 |
| E | E553, K558, P208, E318, E324, D388, P547, G545, I546, S549, N582, M622, F255, V539, D392, L329, I571, L540, V577, A295, I632, I774, M629, E636, A583, M638, Q385, S635, D677 |
| F | G652, T198, R561, A649, A292, Q822, V609, T282, N200, L566, V193, N201, V311, L280, E279, M551, L817, L614, V203, V548, M403, E557, T375, E553, R550, P651, G648, E759 |

Table S16. The highly central (top 5%) residues for the ClpB hexamer network of the APO setup. These positions are shown in Fig. S18.

| Chain | Top 5% $C_B$ ClpB – Hexamer CPX |
|---|---|
| **A** | D388, A830, R331, E553, S549, K558, Q334, R550, V210, E826, S400, K335, S554, P650, R815, A820, L559, L204, E256, E538, G206, P208, D685, Q564, I205, Q385, G211, T607, A682, G652, S644, E678, R556 |
| **B** | Y671, I546, L540, D392, G652, G660, V654, A665, P208, K199, A537, A249, R252, Q334, V666, I205, P650, G655, P651, M638, A649, A397, M551, E396, L646, G648, V548, G209 |
| **C** | V548, R550, V654, R556, D388, P651, Y653, P650, G652, V210, A249, L204, G211, E330, E256, L376, T607, K389, V539, K250, E396, R561 |
| **D** | P208, A328, R588, Y653, V548, Q334, P650, G545, L540, I205, P651, L245, I323, L646, T607, C310, I754, L240, A583, T700, R756, V246, S644, T314, V193, N200 |
| **E** | N711, Y380, A327, G545, L540, P208, L646, D392, V338, E340, D593, E396, A665, A649, V539, T607, Y671, A592, S377, F333, D701, S672, A541, A583, P670, Q192, D708, F601, T314, L386, L675, L604, I181, D633, S400, T375, N582 |
| **F** | G243, I598, T198, L240, I777, R550, L782, V570, V180, V311, V548, G610, M403, I774, T346, Q822, F621, A373, V193, R781, I536, L566, N201, L779, M622, N200, K616, R188, L817 |

Table S17. The highly central (top 5%) residues for the ClpB hexamer network of the CPX setup. These positions are shown in Fig. S18.

| ClpB - Monomer | | |
|---|---|---|
| **Setup** | **APO** | **CPX** |
| **WA1 to PL1**<br>(0.66/0.91)<br>Avg. Path (0.82/1.067) | T213 - E217 - L168 - I166 - Y164 - R258 | T213 - E217 - L168 - A239 - T165 - V262 - R258 |
| **WA2 to PL3**<br>(0.61/0.51)<br>Avg. Path (0.79/0.65) | T612 - K616 - M629 - I674 - L662 - G660 | T612 - C615 - I674 - L662 - G660 |
| **WA2 to PL1**<br>(2.78/2.51)<br>Avg. Paths (2.85/2.60) | T612 - K616 - A619 - V673 - Y671 - Y380 - S377 - L386 - E340 - V338 - V336 - Q334 - N201 - L308 - A267 - N264 - G261 - R258 | T612 - C615 - V673 - Y671 - Y380 - S377 - Q385 - E207 - T315 - E318 - Y322 - G285 - A295 - E256 - R258 |

| | | |
|---|---|---|
| **WA1 to PL3** (1.52/1.31) Avg. Paths (1.77/1.50) | G206 - Q385 - D383 - R667 - T663 - G660 | G206 - P208 - R384 - A382 - D708 - T663 - G660 |

Table S18. Optimal paths for the ClpB monomer for the ATP binding pockets in NBD1 and NBD2 (Walker A1 - G206/T213, Walker A2 - G605/T612) to pore loops with conserved tyrosine residues in NBD1 and NBD2 (PL1 - Y251/R258, PL3 - Y653/G660). The path lengths and average lengths from suboptimal paths are provided under the setup name in order of (APO/CPX).

| **ClpB – Hexamer Intra-Protomer Paths – WA1 to PL1** | | |
|---|---|---|
| **Setup** | **APO** | **CPX** |
| **Chain A** (0.54/1.31) Avg. Path (0.70/1.70) | A:G605 - A:F603 - A:I715 - A:I674 - A:L662 - A:G660 | A:T612 - A:C615 - A:I674 - A:L662 - A:G660 |
| **Chain B** (0.87/1.15) Avg. Path (0.96/1.33) | B:T612 - B:C615 - B:I674 - B:L662 - B:G660 | B:T612 - B:I715 - B:I674 - B:L662 - B:G660 |
| **Chain C** (1.06/2.01) Avg. Paths (1.34/2.33) | C:G605 - C:F603 - C:I715 - C:I674 - C:L662 - C:G660 | C:T612 - C:L675 - C:V673 - C:V666 - C:T663 - C:G660 |
| **Chain D** (1.05/1.35) Avg. Paths (1.31/1.72) | D:T612 - D:C615 - D:V673 - D:V666 - D:T663 - D:G660 | D:T612 - D:D677 - D:I632 - D:L662 - D:G660 |
| **Chain E** (0.68/0.94) Avg. Paths (0.89/1.19) | E:T612 - E:C615 - E:I674 - E:L662 - E:G660 | E:T612 - E:C615 - E:I674 - E:L662 - E:G660 |
| **Chain F** (0.66/0.91) Avg. Paths (0.89/1.28) | F:G605 - F:I715 - F:I674 - F:L662 - F:G660 | F:G605 - F:M716 - F:L676 - F:I632 - F:G660 |

Table S19. Intra-protomer optimal paths for the ClpB hexamer from the pore loop with conserved tyrosine residue in NBD1 (PL1 - Y251/R258) to the ATP binding pockets in NBD1 (Walker A1 - G206/T213). The residues in the paths are indicated as (chain):(amino acid)(residue ID). The path lengths and average lengths from suboptimal paths are provided under the chain names in order of (APO/CPX).

| ClpB – Hexamer Intra-Protomer Paths – WA2 to PL3 | | |
|---|---|---|
| **Setup** | **APO** | **CPX** |
| **Chain A** (0.54/1.31) Avg. Path (0.70/1.70) | A:G605 - A:F603 - A:I715 - A:I674 - A:L662 - A:G660 | A:T612 - A:C615 - A:I674 - A:L662 - A:G660 |
| **Chain B** (0.87/1.15) Avg. Path (0.96/1.33) | B:T612 - B:C615 - B:I674 - B:L662 - B:G660 | B:T612 - B:I715 - B:I674 - B:L662 - B:G660 |
| **Chain C** (1.06/2.01) Avg. Paths (1.34/2.33) | C:G605 - C:F603 - C:I715 - C:I674 - C:L662 - C:G660 | C:T612 - C:L675 - C:V673 - C:V666 - C:T663 - C:G660 |
| **Chain D** (1.05/1.35) Avg. Paths (1.31/1.72) | D:T612 - D:C615 - D:V673 - D:V666 - D:T663 - D:G660 | D:T612 - D:D677 - D:I632 - D:L662 - D:G660 |
| **Chain E** (0.68/0.94) Avg. Paths (0.89/1.19) | E:T612 - E:C615 - E:I674 - E:L662 - E:G660 | E:T612 - E:C615 - E:I674 - E:L662 - E:G660 |
| **Chain F** (0.66/0.91) Avg. Paths (0.89/1.28) | F:G605 - F:I715 - F:I674 - F:L662 - F:G660 | F:G605 - F:M716 - F:L676 - F:I632 - F:G660 |

Table S20. Intra-protomer optimal paths for the ClpB hexamer from the pore loop with conserved tyrosine residue in NBD2 (PL3 - Y653/G660) to the ATP binding pockets in NBD2 (Walker A2 - G605/T612). The residues in the paths are indicated as (chain):(amino acid)(residue ID). The path lengths and average lengths from suboptimal paths are provided under the chain names in order of (APO/CPX).

| ClpB – Hexamer Intra-Protomer Paths – WA2 to PL1 | | |
|---|---|---|
| **Setup** | **APO** | **CPX** |

| | | |
|---|---|---|
| **Chain A** (2.97/4.49) Avg. Path (3.01/4.57) | A:T612 - A:K616 - A:F621 - A:K558 - A:M551 - A:V548 - A:A537 - A:L393 - A:K389 - A:Q385 - A:V210 - A:L204 - A:V311 - A:F276 - A:L240 - A:L245 - A:R258 | A:T612 - A:C615 - A:L618 - A:M562 - A:K558 - A:E553 - A:R550 - A:V548 - A:L540 - A:L393 - A:A390 - A:L386 - A:G211 - A:A214 - A:E217 - A:A239 - A:Y164 - A:R258 |
| **Chain B** (3.31/4.68) Avg. Path (3.37/4.79) | B:T612 - B:C615 - B:V673 - B:Y671 - B:I546 - B:A541 - B:V539 - B:E396 - A:V193 - A:P202 - A:V311 - A:F276 - A:L240 - A:L245 - A:F255 - A:G253 - B:Y251 | B:T612 - B:L675 - B:V673 - B:Y671 - B:I546 - B:A541 - B:V539 - B:E396 - A:V193 - A:V311 - A:F276 - A:L240 - A:L245 - A:F255 - A:R252 - B:Y251 |
| **Chain C** (5.53/5.59) Avg. Paths (5.59/5.68) | C:G605 - C:F603 - C:F601 - C:R756 - C:Q692 - D:D677 - D:I715 - D:V580 - D:I584 - D:R588 - D:M551 - D:S549 - D:L540 - D:E396 - C:R196 - C:L194 - C:I274 - C:L266 - C:V262 - C:R258 | C:T612 - C:C615 - C:L618 - C:M562 - C:L559 - C:R556 - C:R550 - C:V548 - C:A537 - C:L393 - C:D388 - C:G211 - C:A214 - C:E217 - C:L168 - C:T165 - C:V262 - C:R258 |
| **Chain D** (3.79/4.77) Avg. Paths (3.91/4.89) | D:T612 - D:C615 - D:L618 - D:M562 - D:K558 - D:E555 - D:M552 - D:R550 - D:V548 - D:Y380 - D:H378 - D:L386 - D:E340 - D:V338 - D:L204 - D:V311 - D:L275 - D:L266 - D:V262 - D:R258 | D:G605 - D:F603 - D:F601 - D:R756 - E:T607 - E:K611 - E:C615 - E:V673 - E:Y671 - E:A541 - E:V539 - E:E396 - D:R196 - D:L194 - D:I274 - D:L266 - D:G261 - D:R258 |
| **Chain E** (4.80/4.88) Avg. Paths (4.85/4.99) | E:T612 - E:K616 - E:A619 - E:F621 - E:K558 - E:E553 - E:S549 - E:I546 - E:A541 - E:V539 - E:E396 - D:R196 - D:N201 - D:L308 - D:L263 - D:L259 - D:E256 - D:G253 - E:Y251 | E:T612 - E:C615 - E:V673 - E:Y671 - E:A541 - E:E538 - E:E535 - E:V531 - E:I365 - E:K354 - E:G352 - E:I181 - E:D184 - E:R188 - E:I191 - E:L194 - E:L275 - E:L266 - E:V262 - E:R258 |
| **Chain F** (3.45/4.45) Avg. Paths (3.52/4.51) | F:T612 - F:C615 - F:L618 - F:F621 - F:M562 - F:K558 - F:S554 - F:M552 - F:T375 - F:V343 - F:P341 - F:A339 - F:V336 - F:P202 - F:C310 - F:L308 - F:L263 - F:R258 | F:T612 - F:C615 - F:L618 - F:F621 - F:R550 - F:V548 - F:A537 - F:A372 - F:A374 - F:S342 - F:E340 - F:V338 - F:L204 - F:P202 - F:C310 - F:L308 - F:L263 - F:K260 - F:R258 |

Table S21. Intra-protomer optimal paths for the ClpB hexamer from the pore loop with conserved tyrosine residue in NBD1 (PL1 - Y251/R258) to the ATP binding pockets in NBD2 (Walker A2 - G605/T612). The residues in the paths are indicated as (chain):(amino acid)(residue ID). The path lengths and average lengths from suboptimal paths are provided under the chain names in order of (APO/CPX).

| ClpB – Hexamer Intra-Protomer Paths – WA1 to PL3 | | |
|---|---|---|
| **Setup** | **APO** | **CPX** |

| | | |
|---|---|---|
| **Chain A**<br>(2.69/3.80)<br>Avg. Path (2.75/3.90) | A:G206 - A:Q385 - A:K389 - A:L393 - A:A537 - A:V548 - A:M551 - A:K558 - A:F621 - A:A619 - A:S672 - A:A665 - A:G660 | A:T213 - A:I215 - A:T190 - A:V193 - B:E396 - B:V539 - B:A541 - B:I546 - B:Y671 - B:V666 - B:L662 - B:R645 - A:P651 - A:Y653 |
| **Chain B**<br>(2.48/3.30)<br>Avg. Path (2.60/4.45) | B:G206 - B:Q385 - B:K389 - B:L393 - B:L540 - B:I546 - B:Y671 - B:R669 - B:A665 - B:G660 | B:G206 - B:Q385 - B:K389 - B:L393 - B:L540 - B:I546 - B:Y671 - B:A665 - B:G660 |
| **Chain C**<br>(4.42/5.30)<br>Avg. Paths (4.48/5.40) | C:G206 - C:L204 - C:P202 - C:V193 - D:E396 - D:L540 - D:S549 - D:M551 - D:R556 - D:L559 - D:F621 - D:A619 - D:S672 - D:A665 - D:Y661 - D:R645 - C:P651 - C:Y653 | C:T213 - C:I215 - C:T190 - C:Q192 - D:S399 - D:A397 - D:E535 - D:E538 - D:A541 - D:I546 - D:V548 - D:R588 - D:I584 - D:V713 - D:I674 - D:L662 - D:R645 - C:P651 - C:Y653 |
| **Chain D**<br>(3.78/3.47)<br>Avg. Paths (3.88/3.61) | D:G206 - D:E340 - D:L386 - D:H378 - D:Y380 - D:V548 - D:M551 - D:A589 - D:S587 - D:N711 - D:V666 - D:T663 - D:G660 | D:T213 - D:V216 - D:T190 - D:V193 - E:E396 - E:V539 - E:A541 - E:Y671 - E:A665 - E:G660 - D:P651 - D:Y653 |
| **Chain E**<br>(4.20/3.32)<br>Avg. Paths (4.27/3.54) | E:G206 - E:V338 - E:E340 - E:L386 - E:A390 - E:L393 - E:V539 - E:A541 - E:I546 - E:S549 - E:E553 - E:K558 - E:F621 - E:A619 - E:V673 - E:V666 - E:T663 - E:G660 | E:G206 - E:V338 - E:E340 - E:L386 - E:K389 - E:L393 - E:V539 - E:A541 - E:Y671 - E:A665 - E:G660 |
| **Chain F**<br>(2.96/4.05)<br>Avg. Paths (3.08/4.12) | F:G206 - F:A339 - F:P341 - F:V343 - F:T375 - F:M552 - F:S554 - F:K558 - F:M562 - F:F621 - F:A619 - F:S672 - F:L662 - F:G660 | F:G206 - F:V338 - F:E340 - F:S342 - F:A374 - F:A372 - F:A537 - F:V548 - F:R550 - F:F621 - F:L618 - F:C615 - F:I674 - F:L662 - F:G660 |

Table S22. Intra-protomer optimal paths for the ClpB hexamer from the pore loop with conserved tyrosine residue in NBD2 (PL3 - Y653/G660) to the ATP binding pockets in NBD2 (Walker A1 - G206/T213). The residues in the paths are indicated as (chain):(amino acid)(residue ID). The path lengths and average lengths from suboptimal paths are provided under the chain names in order of (APO/CPX).

| **ClpB – Hexamer Inter-Protomer Paths – WA1 to PL1** | | |
|---|---|---|
| **Setup** | **APO** | **CPX** |

| | | |
|---|---|---|
| **Chain A** (1.22/2.23) Avg. Path (1.43/2.34) | B:T213 - B:E217 - B:A239 - B:D241 - B:G243 - A:E256 - A:R258 | B:G206 - B:Q385 - B:K389 - B:D392 - B:D395 - A:R196 - A:L194 - A:V311 - A:F276 - A:A239 - A:Y164 - A:R258 |
| **Chain B** (2.01/2.54) Avg. Path (2.12/2.78) | C:G206 - C:P208 - C:D388 - C:D392 - C:D395 - B:R196 - B:H309 - B:L266 - B:V262 - B:R258 | C:T213 - C:E217 - C:L168 - C:T165 - C:V262 - C:R258 - C:F255 - C:Y251 - B:Y251 |
| **Chain C** (1.79/2.16) Avg. Paths (2.04/2.48) | D:T213 - D:E217 - D:A239 - D:D241 - D:G243 - C:E256 - C:R258 | D:T213 - D:F276 - D:L240 - D:L245 - C:E256 - C:R258 |
| **Chain D** (1.83/2.30) Avg. Paths (1.94/2.79) | E:G206 - E:V338 - E:E340 - E:L386 - E:I391 - E:D395 - D:R196 - D:L194 - D:H309 - D:L266 - D:V262 - D:R258 | E:G206 - E:P208 - D:A328 - D:N297 - D:A295 - D:F255 - D:R258 |
| **Chain E** (2.06/3.26) Avg. Paths (2.50/3.80) | F:T213 - F:A239 - F:M242 - E:A328 - E:K300 - E:M298 - E:R258 | F:T213 - F:A239 - F:D241 - F:G243 - E:A327 - E:L329 - E:K300 - E:M298 - E:R258 |
| **Chain F** (1.57/2.23) Avg. Paths (1.65/2.34) | A:G206 - A:Q385 - A:K389 - A:D392 - A:D395 - F:R196 - F:N201 - F:L308 - F:L263 - F:R258 | A:G206 - A:Q385 - A:K389 - A:D392 - A:D395 - F:R196 - F:L194 - F:C310 - F:L308 - F:L263 - F:K260 - F:R258 |

Table S23. Inter-protomer optimal paths for the ClpB hexamer from the pore loop with conserved tyrosine residue in NBD1 (PL1 - Y251/R258) to the ATP binding pockets in NBD1 (Walker A1 - G206/T213). The residues in the paths are indicated as (chain):(amino acid)(residue ID). The path lengths and average lengths from suboptimal paths are provided under the chain names in order of (APO/CPX).

| **ClpB – Hexamer Inter-Protomer Paths – WA2 to PL3** | | |
|---|---|---|
| **Setup** | **APO** | **CPX** |
| **Chain A** (1.43/1.57) Avg. Path (1.63/1.86) | B:T612 - B:L675 - B:D633 - A:L699 - A:T663 - A:G660 | B:T612 - B:L675 - B:I632 - B:L646 - A:P651 - A:Y653 |
| **Chain B** (1.50/1.91) Avg. Path (1.78/2.17) | C:G605 - C:S718 - C:V679 - C:A682 - C:S642 - B:Y653 | C:G605 - C:T607 - B:R756 - B:S600 - B:V714 - B:I674 - B:L662 - B:G660 |
| **Chain C** (1.64/1.73) Avg. Paths (1.83/2.20) | D:G605 - D:F603 - D:T717 - D:E678 - C:N688 - C:V643 - C:G660 | D:T612 - D:D677 - D:I632 - D:R645 - C:P651 - C:Y653 |

| | | |
|---|---|---|
| **Chain D** (1.31/1.67) Avg. Paths (1.53/2.00) | E:T612 - E:L676 - E:M634 - E:F637 - E:S642 - D:Y653 | E:T612 - E:L675 - E:I632 - E:L646 - D:P651 - D:Y653 |
| **Chain E** (1.98/2.57) Avg. Paths (2.07/2.76) | F:G605 - F:T607 - F:V811 - F:D809 - F:P816 - F:R819 - F:Q822 - F:E826 - E:R586 - E:I598 - E:F709 - E:T663 - E:G660 | F:T612 - F:G610 - F:A814 - F:L817 - F:I821 - F:E826 - E:R586 - E:I584 - E:V713 - E:S672 - E:A665 - E:G660 |
| **Chain F** (1.77/3.08) Avg. Paths (1.92/3.21) | A:G605 - A:F603 - A:I715 - A:I674 - A:L662 - A:G660 - A:E658 - F:P651 - F:Y653 | A:G605 - A:G608 - A:L817 - A:I821 - A:E826 - A:A830 - F:I598 - F:S600 - F:V714 - F:I674 - F:L662 - F:G660 |

Table S24. Inter-protomer optimal paths for the ClpB hexamer from the pore loop with conserved tyrosine residue in NBD2 (PL3 - Y653/G660) to the ATP binding pockets in NBD2 (Walker A2 - G605/T612). The residues in the paths are indicated as (chain):(amino acid)(residue ID). The path lengths and average lengths from suboptimal paths are provided under the chain names in order of (APO/CPX).

| **ClpB – Hexamer Inter-Protomer Paths – WA2 to PL1** | | |
|---|---|---|
| **Setup** | **APO** | **CPX** |
| **Chain A** (2.87/3.62) Avg. Path (2.94/3.74) | B:T612 - B:C615 - B:V673 - B:Y671 - B:I546 - B:A541 - B:V539 - B:E396 - A:R196 - A:L194 - A:H309 - A:L275 - A:L266 - A:V262 - A:R258 | B:T612 - B:L675 - B:V673 - B:Y671 - B:I546 - B:A541 - B:V539 - B:E396 - A:V193 - A:V311 - A:F276 - A:A239 - A:Y164 - A:R258 |
| **Chain B** (4.27/5.41) Avg. Path (4.33/5.50) | C:G605 - C:P765 - C:G767 - C:I771 - C:L817 - C:I821 - C:E826 - C:A830 - B:S587 - B:V548 - B:A541 - B:V539 - B:E396 - A:V193 - A:P202 - A:V311 - A:F276 - A:L240 - A:L245 - A:F255 - A:G253 - B:Y251 | C:T612 - C:C615 - C:L618 - C:M562 - C:L559 - C:R556 - C:R550 - C:V548 - C:A541 - C:V539 - C:E396 - B:R196 - B:L194 - B:L308 - B:L266 - B:V262 - B:R258 |
| **Chain C** (3.79/4.75) Avg. Paths (3.87/4.88) | D:T612 - D:C615 - D:L618 - D:M562 - D:K558 - D:E555 - D:M551 - D:S549 - D:L540 - D:E396 - C:R196 - C:L194 - C:I274 - C:L266 - C:V262 - C:R258 | D:T612 - D:L614 - D:V577 - D:S581 - D:R585 - D:R588 - D:V548 - D:I546 - D:A541 - D:E538 - D:E535 - D:A397 - D:S399 - C:L194 - C:I274 - C:L266 - C:V262 - C:R258 |
| **Chain D** (3.94/3.30) Avg. Paths (4.03/3.46) | E:T612 - E:K616 - E:A619 - E:F621 - E:K558 - E:E553 - E:S549 - E:I546 - E:A541 - E:V539 - E:E396 - D:R196 - D:L194 - D:H309 - D:L266 - D:V262 - D:R258 | E:T612 - E:C615 - E:V673 - E:Y671 - E:A541 - E:V539 - E:E396 - D:R196 - D:L194 - D:I274 - D:L266 - D:G261 - D:R258 |

| | | |
|---|---|---|
| **Chain E**<br>(3.56/4.18)<br>Avg. Paths (3.65/4.28) | F:T612 - F:C615 - F:L618 - F:F621 - F:M562 - F:K558 - F:S554 - F:M552 - F:V548 - F:E538 - F:A397 - F:S400 - F:M403 - E:Q195 - E:I274 - E:L238 - E:V262 - E:R258 | F:T612 - F:C615 - F:L618 - F:F621 - F:R550 - F:V548 - F:E538 - F:E535 - F:A397 - F:S400 - F:M403 - E:Q192 - E:L194 - E:L275 - E:L266 - E:V262 - E:R258 |
| **Chain F**<br>(3.02/4.41)<br>Avg. Paths (3.06/4.49) | A:T612 - A:K616 - A:F621 - A:K558 - A:M551 - A:S549 - A:P547 - A:A541 - A:V539 - A:E396 - F:R196 - F:N201 - F:L308 - F:L263 - F:R258 | A:T612 - A:C615 - A:L618 - A:M562 - A:K558 - A:E553 - A:R550 - A:V548 - A:A541 - A:V539 - A:E396 - F:V193 - F:N201 - F:L308 - F:L263 - F:K260 - F:R258 |

Table S25. Inter-protomer optimal paths for the ClpB hexamer from the pore loop with conserved tyrosine residue in NBD1 (PL1 - Y251/R258) to the ATP binding pockets in NBD2 (Walker A2 - G605/T612). The residues in the paths are indicated as (chain):(amino acid)(residue ID). The path lengths and average lengths from suboptimal paths are provided under the chain names in order of (APO/CPX).

| **ClpB – Hexamer Inter-Protomer Paths – WA1 to PL3** | | |
|---|---|---|
| **Setup** | **APO** | **CPX** |
| **Chain A**<br>(3.31/4.02)<br>Avg. Path (3.42/4.15) | B:G206 - B:Q385 - B:K389 - B:L393 - B:L540 - B:I546 - B:Y671 - B:V673 - B:L675 - B:D633 - A:L699 - A:T663 - A:G660 | B:G206 - B:Q385 - B:K389 - B:L393 - B:L540 - B:I546 - B:Y671 - B:V666 - B:L662 - B:R645 - A:P651 - A:Y653 |
| **Chain B**<br>(3.85/5.14)<br>Avg. Path (3.91/5.23) | C:G206 - C:P208 - C:D388 - C:D392 - B:Q334 - B:V336 - B:V338 - B:E340 - B:T346 - B:A373 - B:A537 - B:P547 - B:Y671 - B:R669 - B:A665 - B:G660 | C:T213 - C:G211 - C:D388 - C:D392 - C:D395 - B:V193 - B:T190 - B:I215 - B:G211 - B:P387 - B:A390 - B:I536 - B:E538 - B:A541 - B:I546 - B:Y671 - B:A665 - B:G660 |
| **Chain C**<br>(4.48/3.76)<br>Avg. Paths (4.59/3.89) | D:G206 - D:E340 - D:L386 - D:H378 - D:Y380 - D:V548 - D:R550 - D:M552 - D:R556 - D:L559 - D:F621 - D:A619 - D:S672 - D:A665 - D:Y661 - D:R645 - C:P651 - C:Y653 | D:T213 - D:V216 - D:T190 - D:V193 - E:E396 - E:V539 - E:A541 - E:Y671 - E:A665 - E:G660 - D:P651 - D:Y653 - C:Y653 |

| | | |
|---|---|---|
| **Chain D** (4.86/4.19) Avg. Paths (4.90/4.40) | E:G206 - E:V338 - E:E340 - E:L386 - E:I391 - E:D395 - D:V193 - D:V336 - D:V338 - D:E340 - D:L386 - D:H378 - D:Y380 - D:V548 - D:M551 - D:A589 - D:S587 - D:N711 - D:V666 - D:T663 - D:G660 | E:G206 - E:V338 - E:E340 - E:L386 - E:K389 - E:L393 - E:V539 - E:A541 - E:Y671 - E:A665 - E:G660 - D:P651 - D:Y653 |
| **Chain E** (4.51/5.63) Avg. Paths (4.57/5.70) | F:G206 - F:A339 - F:P341 - F:V343 - F:T375 - F:M552 - F:S554 - F:K558 - F:M562 - F:L566 - F:Q573 - F:H764 - F:L766 - F:I771 - F:L817 - F:I821 - F:E826 - E:R586 - E:I598 - E:F709 - E:T663 - E:G660 | F:G206 - F:V338 - F:P341 - F:D345 - F:I347 - F:L350 - F:I394 - F:A397 - F:S400 - F:M403 - E:Q192 - E:R188 - E:D184 - E:I181 - E:G352 - E:K354 - E:I365 - E:V531 - E:E535 - E:E538 - E:A541 - E:Y671 - E:A665 - E:G660 |
| **Chain F** (3.77/5.25) Avg. Paths (3.82/5.30) | A:G206 - A:Q385 - A:K389 - A:L393 - A:E396 - F:Q192 - F:R189 - F:E186 - F:A339 - F:P341 - F:V343 - F:T375 - F:M552 - F:S554 - F:K558 - F:M562 - F:F621 - F:A619 - F:S672 - F:L662 - F:G660 | A:G206 - A:Q385 - A:K389 - A:L393 - A:E396 - F:V193 - F:V336 - F:V338 - F:E340 - F:S342 - F:A374 - F:A372 - F:A537 - F:V548 - F:R550 - F:F621 - F:L618 - F:C615 - F:I674 - F:L662 - F:G660 |

Table S26. Inter-protomer optimal paths for the ClpB hexamer from the pore loop with conserved tyrosine residue in NBD2 (PL3 - Y653/G660) to the ATP binding pockets in NBD2 (Walker A1 - G206/T213). The residues in the paths are indicated as (chain):(amino acid)(residue ID). The path lengths and average lengths from suboptimal paths are provided under the chain names in order of (APO/CPX).

| PL1-WA1 | | | | PL3-WA2 | | | | PL1-WA2 | | | | PL3-WA1 | | | |
|---|---|---|---|---|---|---|---|---|---|---|---|---|---|---|---|
| Notes | SS | APO | CPX | Notes | SS | APO | CPX | Notes | SS | APO | CPX | Notes | SS | APO | CPX |
| | L1 | 0.421 | - | Sink | H16 | 0.2102 | 0.3915 | | S1 | - | 0.103 | Sink | L4 | - | 0.3888 |
| | S1 | 0.3117 | 0.3515 | | S9 | 0.2668 | - | | H1 | - | 0.1245 | | H11 | 0.2455 | 0.2672 |
| | H1 | 0.4135 | 0.3952 | PL4 | H17 | 0.1557 | - | | L3 | 0.507 | - | | L12 | 0.596 | 0.5357 |
| Sink | H3 | 0.4975 | 0.5808 | | H18 | 0.2555 | 0.3033 | WA1 | L4 | - | 0.4945 | | H12 | 0.1648 | - |
| | S3 | 0.5695 | 0.246 | | L20 | - | 0.1805 | WA1 | H3 | - | 0.1465 | | H13 | 0.1695 | - |
| | H4 | 0.2815 | 0.1495 | WB2 | S10 | 0.2696 | 0.2777 | Source | L6 | - | 0.143 | PL4 | H17 | 0.114 | - |
| Source | L6 | 0.1507 | 0.144 | | | | | Source | H5 | 0.2828 | 0.2654 | | H18 | 0.3231 | 0.2772 |
| Source | H5 | 0.2519 | 0.2529 | | | | | WB1 | S4 | 0.171 | - | | L20 | 0.4408 | 0.1975 |
| | | | | | | | | PL2 | L8 | 0.1852 | 0.423 | | S12 | - | 0.3515 |
| | | | | | | | | | H7 | 0.1506 | - | | L24 | - | 0.1414 |
| | | | | | | | | | L9 | 0.242 | - | | | | |
| | | | | | | | | | H8 | - | 0.5116 | | | | |
| | | | | | | | | | H9 | 0.302 | - | | | | |
| | | | | | | | | | S6 | 0.8218 | - | | | | |
| | | | | | | | | | L11 | 0.5052 | - | | | | |
| | | | | | | | | | H11 | 0.6598 | 0.3278 | | | | |
| | | | | | | | | | L12 | 0.74 | 0.3726 | | | | |
| | | | | | | | | Sink | H16 | 0.5405 | 0.9335 | | | | |
| | | | | | | | | | L20 | 0.9965 | 0.2905 | | | | |
| | | | | | | | | WB2 | S10 | 0.803 | 0.3905 | | | | |
| | | | | | | | | | S12 | - | 0.444 | | | | |
| | | | | | | | | | L24 | - | 0.2775 | | | | |
| | | | | | | | | | S13 | - | 0.3155 | | | | |

Table S27. Average node degeneracy of secondary structures comprising allosteric communication networks in monomeric ClpB. The four panels show the average node degeneracy for PL1-WA1, PL3-WA2, PL1-WA2, and PL3-WA1 communications of monomeric ClpB in APO and CPX setups. The protomer origin of secondary structures is listed. A dash ("–") indicates that the secondary structure did not participate in the network in that setup. Values are shaded on a green gradient from light (degeneracy ≈ 0) to dark (degeneracy ≈ 1). The functional regions and the sink and source regions of the network, that the listed secondary structure consisted of, are noted.

| A | | | | B | | | | C | | | | D | | | | E | | | | F | | | |
|---|---|---|---|---|---|---|---|---|---|---|---|---|---|---|---|---|---|---|---|---|---|---|---|
| Notes | SS | APO | CPX | Notes | SS | APO | CPX | Notes | SS | APO | CPX | Notes | SS | APO | CPX | Notes | SS | APO | CPX | Notes | SS | APO | CPX |
| SS in protomer A | | | | SS in protomer B | | | | SS in protomer C | | | | SS in protomer D | | | | SS in protomer E | | | | SS in protomer F | | | |
| | L1 | 0.214 | 0.863 | | L1 | 0.375 | - | | L1 | 0.156 | 0.239 | | S1 | - | 0.294 | | L1 | 0.244 | - | | L3 | 0.447 | 0.341 |
| | S1 | - | 0.303 | | S1 | 0.202 | 0.303 | | S1 | 0.197 | 0.272 | | H1 | - | 0.296 | | S1 | 0.132 | - | | S2 | 0.414 | 0.356 |
| | H1 | - | 0.643 | | H1 | 0.349 | 0.365 | | H1 | 0.263 | 0.296 | | H2 | 0.110 | - | | H1 | 0.128 | - | Source | H5 | 0.326 | 0.531 |
| | S2 | 0.348 | - | Sink | H3 | 0.283 | 0.262 | Sink | H3 | 0.273 | 0.303 | Sink | H3 | 0.248 | 0.280 | | H2 | - | 0.233 | WB1 | S4 | 0.106 | 0.215 |
| Sink | H3 | - | 0.330 | | S3 | 0.284 | 0.267 | | S3 | 0.242 | 0.200 | | L5 | 0.107 | - | Sink | H3 | 0.337 | 0.256 | | L9 | 0.417 | 0.409 |
| | S3 | 0.297 | 0.212 | | H4 | 0.206 | - | | H4 | 0.180 | - | | S3 | 0.197 | 0.201 | | S3 | 0.278 | 0.277 | | S5 | 0.121 | 0.244 |
| | H4 | 0.318 | - | Source | H5 | 0.466 | 0.525 | Source | H5 | 0.303 | 0.465 | | H4 | - | 0.143 | Source | H5 | 0.674 | 0.619 | | S6 | 0.192 | 0.153 |
| Source | H5 | 0.247 | 0.143 | WB1 | S4 | - | 0.126 | WB1 | S4 | 0.125 | - | Source | H5 | 0.405 | 0.411 | WB1 | S4 | - | 0.331 | | | | |
| WB1 | S4 | 0.299 | - | | | | | | | | | WB1 | S4 | 0.278 | 0.104 | | L9 | - | 0.270 | | | | |
| | S5 | 0.310 | - | | | | | | | | | | L9 | 0.102 | - | | | | | | | | |
| | | | | | | | | | | | | | S5 | 0.144 | - | | | | | | | | |

Table S28. Average node degeneracy of secondary structures comprising intra-protomer PL1-WA1 allosteric communication networks in hexameric ClpB. The six panels show the average node degeneracy for intra-protomer communications of protomer A-F in APO and CPX setups. The protomer origin of secondary structures is listed. A dash ("–") indicates that the secondary structure did not participate in the network in that setup. Values are shaded on a green gradient from light (degeneracy ≈ 0) to dark (degeneracy ≈ 1). The functional regions and the sink and source regions of the network, that the listed secondary structure consisted of, are noted.

| A | | | | B | | | | C | | | | D | | | | E | | | | F | | | |
|---|---|---|---|---|---|---|---|---|---|---|---|---|---|---|---|---|---|---|---|---|---|---|---|
| Notes | SS | APO | CPX | Notes | SS | APO | CPX | Notes | SS | APO | CPX | Notes | SS | APO | CPX | Notes | SS | APO | CPX | Notes | SS | APO | CPX |
| SS in protomer A | | | | SS in protomer B | | | | SS in protomer C | | | | SS in protomer D | | | | SS in protomer E | | | | SS in protomer F | | | |
| | S8 | 0.360 | - | | S8 | - | 0.176 | | S8 | 0.315 | - | Sink | H16 | 0.316 | 0.124 | Sink | H16 | 0.290 | 0.267 | | S8 | 0.397 | 0.374 |
| Sink | H16 | - | 0.275 | Sink | H16 | 0.295 | 0.161 | Sink | H16 | - | 0.316 | | S9 | - | 0.152 | | S9 | 0.102 | - | | S9 | 0.211 | 0.340 |
| | S9 | 0.145 | - | | S9 | 0.264 | 0.201 | | S9 | 0.116 | - | | H18 | 0.301 | 0.240 | | H18 | 0.308 | 0.269 | | H18 | 0.313 | 0.447 |
| | H18 | 0.282 | 0.303 | | H18 | 0.490 | 0.501 | | H18 | 0.273 | 0.311 | | L20 | 0.153 | - | | L20 | 0.188 | 0.242 | | L20 | 0.174 | - |
| | L20 | 0.138 | 0.136 | WB2 | S10 | 0.262 | 0.328 | | L20 | - | 0.146 | WB2 | S10 | 0.318 | 0.256 | WB2 | S10 | 0.324 | 0.303 | WB2 | S10 | 0.300 | 0.439 |
| WB2 | S10 | 0.265 | 0.304 | | S13 | - | 0.229 | WB2 | S10 | 0.241 | 0.268 | | L24 | 0.117 | 0.173 | | L24 | 0.128 | 0.123 | | S13 | 0.305 | 0.405 |
| | S13 | 0.268 | - | | | | | | L24 | 0.153 | 0.273 | | S13 | 0.142 | 0.168 | | S13 | 0.181 | 0.147 | | L25 | - | 0.145 |
| | | | | | | | | | S13 | 0.295 | 0.267 | | | | | | | | | | | | |
| | | | | | | | | | L25 | 0.128 | - | | | | | | | | | | | | |

Table S29. Average node degeneracy of secondary structures comprising intra-protomer PL3-WA2 allosteric communication networks in hexameric ClpB. The six panels show the average node degeneracy for intra-protomer communications of protomer A-F in APO and CPX setups. The protomer origin of secondary structures is listed. A dash ("–") indicates that the secondary structure did not participate in the network in that setup. Values are shaded on a green gradient from light (degeneracy ≈ 0) to dark (degeneracy ≈ 1). The functional regions and the sink and source regions of the network, that the listed secondary structure consisted of, are noted.

| A | | | |
|---|---|---|---|
| Notes | SS | APO | CPX |
| **SS in protomer A** | | | |
| | H2 | - | 0.479 |
| | L3 | - | 0.103 |
| Sink | H3 | - | 0.489 |
| Sink | L4 | 0.499 | - |
| | L12 | 0.608 | - |
| | H12 | 0.496 | - |
| | H13 | 0.272 | - |
| | L14 | 0.597 | - |
| | H14 | 0.561 | - |
| | L19 | - | 0.991 |
| WA2 | H16 | 0.596 | - |
| | H18 | 0.283 | - |
| | L20 | 0.342 | - |
| | S10 | 0.246 | - |
| **SS in protomer B** | | | |
| | H12 | - | 0.412 |
| | H13 | - | 0.433 |
| | L14 | - | 1.000 |
| PL3 | H17 | - | 0.436 |
| | H18 | - | 0.347 |
| | L20 | - | 1.000 |

| B | | | |
|---|---|---|---|
| Notes | SS | APO | CPX |
| **SS in protomer B** | | | |
| Sink | L4 | 0.342 | 0.159 |
| | H11 | 0.111 | 0.104 |
| | L12 | 0.360 | 0.531 |
| | H12 | 0.364 | 0.300 |
| | H13 | 0.236 | 0.343 |
| | L14 | 0.500 | 1.000 |
| | H18 | 0.295 | 0.310 |
| | L20 | 1.000 | 1.000 |

| C | | | |
|---|---|---|---|
| Notes | SS | APO | CPX |
| **SS in protomer C** | | | |
| | H2 | 0.396 | 0.609 |
| | L3 | 0.141 | - |
| Sink | S2 | 0.832 | - |
| | H13 | - | 0.500 |
| Source | L19 | 0.996 | 0.933 |
| **SS in protomer D** | | | |
| | H12 | 0.449 | 0.484 |
| | H13 | 0.298 | 0.415 |
| | L14 | 0.431 | 0.466 |
| | H14 | 0.327 | - |
| | H15 | 0.262 | 0.621 |
| | L16 | - | 0.203 |
| WA2 | H16 | 0.604 | - |
| | S9 | - | 0.388 |
| PL3 | H17 | 0.959 | 0.884 |
| | H18 | 0.380 | 0.359 |
| | L20 | 0.580 | - |
| WB2 | S10 | - | 0.344 |
| | L24 | 0.230 | 0.124 |
| | S13 | - | 0.730 |

| D | | | |
|---|---|---|---|
| Notes | SS | APO | CPX |
| **SS in protomer D** | | | |
| | S6 | 0.410 | 0.458 |
| | L11 | 0.534 | 0.125 |
| | H11 | 0.381 | 0.384 |
| | L12 | 0.851 | - |
| | L14 | 0.639 | - |
| | H14 | 0.323 | - |
| | H15 | 0.191 | - |
| WA2 | H16 | 0.402 | - |
| Source | L19 | - | 0.867 |
| | H18 | 0.331 | - |
| | L20 | 0.232 | - |
| WB2 | S10 | 0.480 | - |
| | L24 | 0.153 | - |
| **SS in protomer E** | | | |
| | H12 | - | 0.352 |
| | H13 | - | 0.434 |
| PL4 | H17 | - | 0.237 |
| | H18 | - | 0.251 |
| | L20 | - | 1.000 |

| E | | | |
|---|---|---|---|
| Notes | SS | APO | CPX |
| **SS in protomer E** | | | |
| | S6 | 0.813 | 0.982 |
| | L11 | 0.396 | 0.982 |
| | H11 | 0.295 | 0.202 |
| | L12 | 0.782 | 0.422 |
| | H12 | 0.392 | 0.204 |
| | H13 | 0.325 | 0.384 |
| | L14 | 1.000 | - |
| | H14 | 1.000 | - |
| WA2 | H16 | 0.442 | - |
| | H18 | 0.395 | 0.329 |
| | L20 | 0.353 | 0.808 |
| WB2 | S10 | 0.302 | - |
| | L24 | - | 0.198 |

| F | | | |
|---|---|---|---|
| Notes | SS | APO | CPX |
| **SS in protomer F** | | | |
| | S6 | - | 0.991 |
| | L11 | 0.533 | 0.387 |
| | H10 | - | 0.828 |
| | H11 | 0.996 | 0.277 |
| | H13 | - | 0.353 |
| | L14 | 0.857 | 1.000 |
| | H14 | 0.406 | - |
| WA2 | H16 | 0.494 | 0.519 |
| | S9 | - | 0.248 |
| | H18 | 0.261 | 0.442 |
| | L20 | 0.398 | 0.167 |
| WB2 | S10 | 0.115 | 0.295 |

Table S30. Average node degeneracy of secondary structures comprising intra-protomer PL1-WA2 allosteric communication networks in hexameric ClpB. The six panels show the average node degeneracy for intra-protomer communications of protomer A-F in APO and CPX setups. The protomer origin of secondary structures is listed. A dash (“–”) indicates that the secondary structure did not participate in the network in that setup. Values are shaded on a green gradient from light (degeneracy ≈ 0) to dark (degeneracy ≈ 1). The functional regions and the sink and source regions of the network, that the listed secondary structure consisted of, are noted.

| A | | | |
|---|---|---|---|
| Notes | SS | APO | CPX |
| **SS in protomer A** | | | |
| | H2 | - | 0.479 |
| | L3 | - | 0.103 |
| Sink | H3 | - | 0.489 |
| Sink | L4 | 0.499 | - |
| | L12 | 0.608 | - |
| | H12 | 0.496 | - |
| | H13 | 0.272 | - |
| | L14 | 0.597 | - |
| | H14 | 0.561 | - |
| | L19 | - | 0.991 |
| WA2 | H16 | 0.596 | - |
| | H18 | 0.283 | - |
| | L20 | 0.342 | - |
| | S10 | 0.246 | - |
| **SS in protomer B** | | | |
| | H12 | - | 0.412 |
| | H13 | - | 0.433 |
| | L14 | - | 1.000 |
| PL3 | H17 | - | 0.436 |
| | H18 | - | 0.347 |
| | L20 | - | 1.000 |

| B | | | |
|---|---|---|---|
| Notes | SS | APO | CPX |
| **SS in protomer B** | | | |
| Sink | L4 | 0.342 | 0.159 |
| | H11 | 0.111 | 0.104 |
| | L12 | 0.360 | 0.531 |
| | H12 | 0.364 | 0.300 |
| | H13 | 0.236 | 0.343 |
| | L14 | 0.500 | 1.000 |
| | H18 | 0.295 | 0.310 |
| | L20 | 1.000 | 1.000 |

| C | | | |
|---|---|---|---|
| Notes | SS | APO | CPX |
| **SS in protomer C** | | | |
| | H2 | 0.396 | 0.609 |
| | L3 | 0.141 | - |
| Sink | S2 | 0.832 | - |
| | H13 | - | 0.500 |
| Source | L19 | 0.996 | 0.933 |
| **SS in protomer D** | | | |
| | H12 | 0.449 | 0.484 |
| | H13 | 0.298 | 0.415 |
| | L14 | 0.431 | 0.466 |
| | H14 | 0.327 | - |
| | H15 | 0.262 | 0.621 |
| | L16 | - | 0.203 |
| WA2 | H16 | 0.604 | - |
| | S9 | - | 0.388 |
| PL3 | H17 | 0.959 | 0.884 |
| | H18 | 0.380 | 0.359 |
| | L20 | 0.580 | - |
| WB2 | S10 | - | 0.344 |
| | L24 | 0.230 | 0.124 |
| | S13 | - | 0.730 |

| D | | | |
|---|---|---|---|
| Notes | SS | APO | CPX |
| **SS in protomer D** | | | |
| | S6 | 0.410 | 0.458 |
| | L11 | 0.534 | 0.125 |
| | H11 | 0.381 | 0.384 |
| | L12 | 0.851 | - |
| | L14 | 0.639 | - |
| | H14 | 0.323 | - |
| | H15 | 0.191 | - |
| WA2 | H16 | 0.402 | - |
| Source | L19 | - | 0.867 |
| | H18 | 0.331 | - |
| | L20 | 0.232 | - |
| WB2 | S10 | 0.480 | - |
| | L24 | 0.153 | - |
| **SS in protomer E** | | | |
| | H12 | - | 0.352 |
| | H13 | - | 0.434 |
| PL4 | H17 | - | 0.237 |
| | H18 | - | 0.251 |
| | L20 | - | 1.000 |

| E | | | |
|---|---|---|---|
| Notes | SS | APO | CPX |
| **SS in protomer E** | | | |
| | S6 | 0.813 | 0.982 |
| | L11 | 0.396 | 0.982 |
| | H11 | 0.295 | 0.202 |
| | L12 | 0.782 | 0.422 |
| | H12 | 0.392 | 0.204 |
| | H13 | 0.325 | 0.384 |
| | L14 | 1.000 | - |
| | H14 | 1.000 | - |
| WA2 | H16 | 0.442 | - |
| | H18 | 0.395 | 0.329 |
| | L20 | 0.353 | 0.808 |
| WB2 | S10 | 0.302 | - |
| | L24 | - | 0.198 |

| F | | | |
|---|---|---|---|
| Notes | SS | APO | CPX |
| **SS in protomer F** | | | |
| | S6 | - | 0.991 |
| | L11 | 0.533 | 0.387 |
| | H10 | - | 0.828 |
| | H11 | 0.996 | 0.277 |
| | H13 | - | 0.353 |
| | L14 | 0.857 | 1.000 |
| | H14 | 0.406 | - |
| WA2 | H16 | 0.494 | 0.519 |
| | S9 | - | 0.248 |
| | H18 | 0.261 | 0.442 |
| | L20 | 0.398 | 0.167 |
| WB2 | S10 | 0.115 | 0.295 |

Table S31. Average node degeneracy of secondary structures comprising intra-protomer PL3-WA1 allosteric communication networks in hexameric ClpB. The six panels show the average node degeneracy for intra-protomer communications of protomer A-F in APO and CPX setups. The protomer origin of secondary structures is listed. A dash (“–”) indicates that the secondary structure did not participate in the network in that setup. Values are shaded on a green gradient from light (degeneracy ≈ 0) to dark (degeneracy ≈ 1). The functional regions and the sink and source regions of the network, that the listed secondary structure consisted of, are noted.

| | APO | | | CPX | | |
|---|---|---|---|---|---|---|
| | Inter-protomer transition pair | | Percentage (%) | Inter-protomer transition pair | | Percentage (%) |
| A | | | | | | |
| B | L6-B | L6-A | 98.70 | L6-B | L6-A | 90.00 |
| | **L3-A** | **H12-B** | **68.15** | **H2-A** | **H12-B** | **79.25** |
| C | **L3-C** | **H12-D** | **80.95** | | | |
| | S9-D | H20-C | 67.20 | | | |
| D | | | | L17-E | H22-D | 94.85 |
| | | | | **L3-D** | **H12-E** | **81.15** |
| E | L6-E | L6-D | 99.35 | | | |
| | **H2-D** | **H12-E** | **58.85** | | | |
| F | | | | | | |

Table S32. Percentage occurrence of inter-protomer secondary-structure (SS) transition pairs in the intra-protomer PL1–WA2 communication network of hexameric ClpB. For each protomer (A–F) and setup (APO, CPX), the observed SS–protomer pairs (formatted as “SS–Protomer”) greater than 50% are listed along with their occurrence percentages out of the total 20000 dynamic-correlation paths.

| | APO | | | CPX | | |
|---|---|---|---|---|---|---|
| | Inter-protomer transition pair | | Percentage (%) | Inter-protomer transition pair | | Percentage (%) |
| A | | | | L19-A | H17-B | 99.55 |
| | | | | **H12-B** | **H2-A** | **88.15** |
| B | | | | | | |
| C | L19-C | H17-D | 97.30 | L19-C | H17-D | 93.25 |
| | **H12-D** | **H2-C** | **85.75** | **H12-D** | **H2-C** | **95.60** |
| D | | | | **H12-E** | **H2-D** | **85.15** |
| | | | | L19-D | L19-E | 73.35 |
| E | | | | | | |
| F | | | | | | |

Table S33. Percentage occurrence of inter-protomer secondary-structure (SS) transition pairs in the intra-protomer PL3–WA1 communication network of hexameric ClpB. For each protomer (A–F) and setup (APO, CPX), the observed SS–protomer pairs (formatted as “SS–Protomer”) greater than 50% are listed along with their occurrence percentages out of the total 20000 dynamic-correlation paths.

**A**

| Notes | SS | APO | CPX |
|---|---|---|---|
| **SS in protomer A** | | | |
| | L1 | - | 0.979 |
| | S1 | - | 0.366 |
| | H1 | - | 0.265 |
| | H2 | - | 0.262 |
| | L3 | - | 0.502 |
| | S2 | - | 0.101 |
| | H3 | - | 0.230 |
| | S3 | - | 0.584 |
| Source | L6 | 0.194 | - |
| Source | H5 | 0.312 | - |
| | S4 | - | 0.511 |
| | S5 | - | 0.504 |
| **SS in protomer B** | | | |
| | H1 | 0.168 | - |
| Sink | L4 | - | 0.157 |
| Sink | H3 | 0.315 | - |
| | S3 | 0.347 | - |
| | H4 | 0.425 | - |
| WB1 | S4 | 0.167 | - |
| | L12 | - | 0.645 |
| | H12 | - | 0.374 |

**B**

| Notes | SS | APO | CPX |
|---|---|---|---|
| **SS in protomer B** | | | |
| | H2 | 0.326 | - |
| | L3 | 0.274 | - |
| Source | H5 | 0.573 | - |
| | S4 | 0.247 | - |
| | L9 | 0.656 | - |
| **SS in protomer C** | | | |
| | L1 | - | 0.183 |
| | S1 | - | 0.247 |
| | H1 | - | 0.518 |
| Sink | L4 | 0.378 | - |
| Sink | H3 | - | 0.278 |
| | S3 | - | 0.187 |
| | H4 | - | 0.128 |
| WB1 | L6 | - | 0.445 |
| PL1 | H5 | - | 0.309 |
| | L11 | 0.224 | - |
| | H12 | 0.346 | - |

**C**

| Notes | SS | APO | CPX |
|---|---|---|---|
| **SS in protomer C** | | | |
| Source | L6 | 0.486 | 0.105 |
| Source | H5 | 0.417 | 0.892 |
| **SS in protomer D** | | | |
| | H1 | 0.144 | 0.172 |
| Sink | H3 | 0.275 | 0.512 |
| | S3 | 0.365 | 0.256 |
| | H4 | 0.350 | 0.306 |
| WB1 | S4 | 0.157 | 0.618 |

**D**

| Notes | SS | APO | CPX |
|---|---|---|---|
| **SS in protomer D** | | | |
| | H2 | 0.343 | - |
| | L3 | 0.364 | - |
| Source | L6 | - | 0.230 |
| Source | H5 | 0.614 | 0.329 |
| WB1 | S4 | 0.161 | - |
| PL2 | L8 | - | 0.367 |
| | H7 | - | 0.342 |
| | L9 | 0.372 | - |
| | H9 | - | 1.000 |
| **SS in protomer E** | | | |
| Sink | L4 | - | 1.000 |
| | S6 | 0.759 | - |
| | L11 | 0.963 | - |
| | L12 | 0.746 | - |
| | H12 | 0.266 | - |

**E**

| Notes | SS | APO | CPX |
|---|---|---|---|
| **SS in protomer E** | | | |
| Source | L6 | - | 0.112 |
| Source | H5 | 0.218 | 0.208 |
| PL2 | L8 | 0.824 | 0.308 |
| | H7 | 0.331 | 0.375 |
| | H9 | 0.954 | 0.980 |
| **SS in protomer F** | | | |
| Sink | H3 | 0.254 | 0.342 |
| | S3 | 0.312 | 0.490 |
| | H4 | 0.367 | 0.970 |
| WB1 | S4 | 0.141 | - |

**F**

| Notes | SS | APO | CPX |
|---|---|---|---|
| **SS in protomer A** | | | |
| | H2 | - | 0.385 |
| | L3 | 0.527 | 0.739 |
| Source | H5 | 0.477 | 0.590 |
| | L9 | 0.425 | 0.753 |
| **SS in protomer F** | | | |
| | L4 | 0.544 | 0.187 |
| | L11 | - | 0.158 |
| | L12 | 0.286 | 0.486 |
| Sink | H12 | 0.339 | 0.337 |

Table S34. Average node degeneracy of secondary structures comprising inter-protomer PL1-WA1 allosteric communication networks in hexameric ClpB. The six panels show the average node degeneracy for inter-protomer communications of protomer A-F in APO and CPX setups. The protomer origin of secondary structures is listed. A dash ("–") indicates that the secondary structure did not participate in the network in that setup. Values are shaded on a green gradient from light (degeneracy ≈ 0) to dark (degeneracy ≈ 1). The functional regions and the sink and source regions of the network, that the listed secondary structure consisted of, are noted.

**A**

| Notes | SS | APO | CPX |
|---|---|---|---|
| **SS in protomer A** | | | |
| Source | L19 | - | 0.405 |
| | H18 | 0.293 | - |
| | H20 | 0.157 | - |
| | S11 | 0.295 | - |
| | S12 | 0.165 | - |
| | L24 | 0.285 | - |
| **SS in protomer B** | | | |
| | S8 | - | 0.116 |
| Sink | H16 | 0.230 | 0.143 |
| | S9 | 0.653 | 0.297 |
| PL4 | H17 | - | 0.264 |
| | H18 | - | 0.261 |
| WB2 | S10 | 0.373 | 0.249 |
| | S13 | - | 0.219 |

**B**

| Notes | SS | APO | CPX |
|---|---|---|---|
| **SS in protomer B** | | | |
| | S8 | - | 0.324 |
| | S9 | - | 0.140 |
| Source | L19 | 0.253 | - |
| | H18 | - | 0.936 |
| WB2 | S10 | - | 0.359 |
| | S13 | - | 0.249 |
| | H22 | - | 0.324 |
| | L27 | - | 0.278 |
| **SS in protomer C** | | | |
| Sink | L17 | - | 0.500 |
| | S8 | 0.604 | - |
| | S9 | 0.265 | - |
| PL4 | H17 | 0.273 | - |
| | H18 | 0.177 | - |
| WB2 | S10 | 0.181 | - |
| | H19 | 0.206 | - |
| | L21 | 0.131 | - |
| | S13 | 0.309 | - |
| | L25 | 0.268 | - |

**C**

| Notes | SS | APO | CPX |
|---|---|---|---|
| **SS in protomer C** | | | |
| Source | H17 | 0.175 | - |
| | L19 | - | 0.383 |
| | H18 | 0.251 | - |
| | H20 | 0.201 | - |
| | S11 | 0.216 | - |
| **SS in protomer D** | | | |
| | S8 | 0.626 | - |
| | S9 | 0.248 | 0.320 |
| PL4 | H17 | - | 0.250 |
| | H18 | - | 0.163 |
| WB2 | S10 | 0.229 | 0.264 |
| | H19 | 0.234 | - |
| | S13 | 0.317 | 0.162 |
| | L25 | 0.199 | - |
| | S14 | 0.121 | - |

**D**

| Notes | SS | APO | CPX |
|---|---|---|---|
| **SS in protomer D** | | | |
| | H15 | - | 0.576 |
| | H18 | - | 0.282 |
| | L20 | - | 0.409 |
| | S13 | - | 0.982 |
| **SS in protomer E** | | | |
| Sink | H16 | 0.362 | 0.932 |
| | S9 | 0.221 | - |
| PL4 | H17 | 0.324 | - |
| WB2 | S10 | 0.305 | - |
| | S13 | 0.138 | - |
| | H25 | - | 0.380 |

**E**

| Notes | SS | APO | CPX |
|---|---|---|---|
| **SS in protomer E** | | | |
| | H15 | 0.506 | 0.576 |
| | L16 | 0.348 | 0.282 |
| | H18 | 0.370 | 0.409 |
| | L20 | 0.284 | 0.982 |
| WB2 | S10 | 0.121 | - |
| | L24 | 0.175 | - |
| | S13 | 0.638 | - |
| **SS in protomer F** | | | |
| Sink | L17 | 0.477 | 0.932 |
| | L30 | 0.461 | 0.380 |
| | H25 | 0.352 | - |

**F**

| Notes | SS | APO | CPX |
|---|---|---|---|
| **SS in protomer A** | | | |
| | S8 | 0.354 | - |
| | L17 | - | 0.360 |
| | S9 | 0.168 | - |
| Source | L19 | 0.914 | - |
| | H18 | 0.247 | - |
| | L20 | 0.129 | - |
| WB2 | S10 | 0.261 | - |
| | S13 | 0.299 | - |
| | L28 | - | 0.119 |
| | L30 | - | 0.131 |
| | H25 | - | 0.372 |
| **SS in protomer F** | | | |
| | L16 | - | 1.000 |
| | S8 | - | 0.500 |
| | S9 | - | 0.407 |
| | **H18** | - | 0.593 |
| WB2 | S10 | - | 0.334 |
| | S13 | - | 0.454 |

Table S35. Average node degeneracy of secondary structures comprising inter-protomer PL3-WA2 allosteric communication networks in hexameric ClpB. The six panels show the average node degeneracy for inter-protomer communications of protomer A-F in APO and CPX setups. The protomer origin of secondary structures is listed. A dash (“–”) indicates that the secondary structure did not participate in the network in that setup. Values are shaded on a green gradient from light (degeneracy ≈ 0) to dark (degeneracy ≈ 1). The functional regions and the sink and source regions of the network, that the listed secondary structure consisted of, are noted.

| A | | | |
|---|---|---|---|
| Notes | SS | APO | CPX |
| **SS in protomer A** | | | |
| | L1 | - | 0.991 |
| | S1 | - | 0.359 |
| | H1 | - | 0.295 |
| | H2 | 0.319 | 0.277 |
| | L3 | 0.412 | 0.188 |
| | S2 | 0.101 | - |
| | H3 | - | 0.152 |
| | S3 | 0.483 | 0.612 |
| | H4 | 0.347 | |
| Source | H5 | 0.386 | |
| | S4 | 0.397 | 0.506 |
| | L9 | 0.202 | |
| | S5 | 0.397 | 0.506 |
| **SS in protomer B** | | | |
| | H12 | 0.380 | 0.393 |
| | H13 | 0.372 | 0.518 |
| | L14 | 0.500 | 1.000 |
| Sink | H16 | 0.500 | 0.189 |
| | L20 | 1.000 | 1.000 |
| | S10 | 1.000 | 0.685 |
| | S13 | - | 0.188 |

| B | | | |
|---|---|---|---|
| Notes | SS | APO | CPX |
| **SS in protomer A** | | | |
| | H2 | 0.325 | - |
| | L3 | 0.398 | - |
| | S2 | 0.111 | - |
| | S3 | 0.575 | - |
| | H4 | 0.381 | - |
| PL1 | L6 | 0.933 | - |
| PL1 | H5 | 0.213 | - |
| | S4 | 0.377 | - |
| | L9 | 0.159 | - |
| | S5 | 0.426 | - |
| **SS in protomer B** | | | |
| | H2 | - | 0.729 |
| | L3 | - | 0.663 |
| Source | H5 | - | 1.000 |
| | S4 | - | 0.306 |
| | L9 | - | 0.331 |
| | H12 | 0.375 | - |
| | H13 | 0.347 | - |
| | L14 | 1.000 | - |
| | H15 | 0.334 | - |
| **SS in protomer C** | | | |
| | H12 | - | 0.330 |
| | H13 | - | 0.296 |
| | L14 | - | 1.000 |
| | H14 | - | 0.500 |
| Sink | H16 | - | 0.630 |
| | L28 | 1.000 | - |
| | H23 | 0.986 | - |
| | H25 | 0.893 | - |

| C | | | |
|---|---|---|---|
| Notes | SS | APO | CPX |
| **SS in protomer C** | | | |
| | H2 | 0.991 | 0.655 |
| | L3 | 0.751 | 0.466 |
| Source | H5 | 0.973 | 0.997 |
| | S4 | 0.954 | 0.674 |
| | L9 | - | 0.332 |
| **SS in protomer D** | | | |
| | H12 | 0.341 | 0.387 |
| | H13 | 0.277 | 0.439 |
| | L14 | 0.453 | 0.500 |
| | H14 | 0.510 | 0.601 |
| Sink | H16 | 0.466 | 0.367 |

| D | | | |
|---|---|---|---|
| Notes | SS | APO | CPX |
| **SS in protomer D** | | | |
| | H2 | 0.418 | 0.567 |
| | L3 | 0.448 | 0.707 |
| Source | H5 | 0.939 | 0.630 |
| | S4 | 0.223 | 0.557 |
| | L9 | 0.708 | 0.510 |
| **SS in protomer E** | | | |
| | H12 | 0.303 | 0.355 |
| | H13 | 0.314 | 0.517 |
| | L14 | 1.000 | - |
| | H14 | 1.000 | - |
| Sink | H16 | 0.450 | 0.398 |
| | L20 | - | 0.999 |
| | S10 | - | 0.526 |
| | S13 | - | 0.140 |

| E | | | |
|---|---|---|---|
| Notes | SS | APO | CPX |
| **SS in protomer E** | | | |
| | H2 | - | 0.999 |
| | L3 | 1.000 | - |
| | S3 | 1.000 | 0.231 |
| Source | H5 | 1.000 | 0.884 |
| | S4 | 1.000 | 0.500 |
| **SS in protomer F** | | | |
| | H12 | 0.580 | 0.570 |
| | H13 | 0.285 | 0.273 |
| | L14 | 0.613 | 1.000 |
| | H14 | 0.335 | - |
| Sink | H16 | 0.406 | 0.363 |

| F | | | |
|---|---|---|---|
| Notes | SS | APO | CPX |
| **SS in protomer A** | | | |
| | H12 | 0.338 | 0.352 |
| | H13 | 0.339 | 0.302 |
| | L14 | 0.576 | 0.674 |
| | H14 | 0.535 | 0.959 |
| Sink | H16 | 0.329 | 0.651 |
| **SS in protomer F** | | | |
| | H2 | 0.199 | 0.406 |
| | L3 | 0.537 | 0.697 |
| Source | H5 | 0.626 | 0.604 |
| | L9 | 0.397 | 0.771 |

Table S36. Average node degeneracy of secondary structures comprising inter-protomer PL1-WA2 allosteric communication networks in hexameric ClpB. The six panels show the average node degeneracy for inter-protomer communications of protomer A-F in APO and CPX setups. The protomer origin of secondary structures is listed. A dash ("–") indicates that the secondary structure did not participate in the network in that setup. Values are shaded on a green gradient from light (degeneracy ≈ 0) to dark (degeneracy ≈ 1). The functional regions and the sink and source regions of the network, that the listed secondary structure consisted of, are noted.

**A**

| Notes | SS | APO | CPX |
|---|---|---|---|
| **SS in protomer A** | | | |
| | H18 | 0.471 | 0.946 |
| | S11 | 0.315 | - |
| | L24 | 0.119 | - |
| **SS in protomer B** | | | |
| Sink | L4 | 0.277 | 0.126 |
| | H11 | 0.150 | 0.675 |
| | L12 | 0.391 | 0.322 |
| | H12 | 0.376 | 0.336 |
| | H13 | 0.248 | 1.000 |
| | L14 | 0.500 | 0.312 |
| | S9 | 0.630 | 0.351 |
| | L20 | 1.000 | 1.000 |
| | S10 | 0.806 | - |

**B**

| Notes | SS | APO | CPX |
|---|---|---|---|
| **SS in protomer B** | | | |
| | H2 | 0.465 | 0.635 |
| WA1 | S2 | 0.396 | 0.340 |
| | S6 | 0.385 | 0.535 |
| | L11 | 0.514 | 0.661 |
| | H11 | 0.526 | 0.461 |
| | L12 | 0.338 | 0.134 |
| | H12 | 0.382 | 0.568 |
| | H13 | 0.492 | 0.383 |
| | L14 | 0.500 | 1.000 |
| | H18 | 0.356 | 0.353 |
| | L20 | 1.000 | 1.000 |
| **SS in protomer C** | | | |
| Sink | L4 | 0.467 | - |

**C**

| Notes | SS | APO | CPX |
|---|---|---|---|
| **SS in protomer C** | | | |
| Source | L19 | 0.949 | - |
| **SS in protomer D** | | | |
| | H2 | - | 0.459 |
| | L3 | - | 0.124 |
| Sink | H3 | - | 0.383 |
| | S6 | 0.387 | - |
| | L11 | 0.983 | - |
| | H11 | 0.599 | - |
| | L12 | 0.917 | - |
| | L14 | 0.668 | - |
| | H14 | 0.358 | - |
| WA2 | H16 | 0.388 | - |
| | S9 | 0.106 | - |
| PL4 | H17 | 0.794 | - |
| PL3 | L19 | - | 0.904 |
| | H18 | 0.343 | - |
| | L20 | 0.346 | - |
| | S10 | 0.246 | - |
| **SS in protomer E** | | | |
| | H12 | | 0.352 |
| | H13 | | 0.434 |
| PL4 | H17 | | 0.238 |
| | H18 | | 0.251 |
| | L20 | | 1.000 |

**D**

| Notes | SS | APO | CPX |
|---|---|---|---|
| **SS in protomer D** | | | |
| | H2 | 0.991 | - |
| | L3 | 0.185 | - |
| WA1 | S2 | 0.114 | - |
| | S6 | 0.937 | - |
| | L11 | 1.000 | - |
| | H11 | 0.667 | - |
| | L12 | 0.940 | - |
| | L14 | 0.671 | - |
| | H14 | 0.319 | - |
| | H15 | 0.284 | - |
| WA2 | H16 | 0.415 | |
| Source | L19 | - | 0.809 |
| | H18 | 0.536 | - |
| | L20 | 0.169 | - |
| | S10 | 0.526 | - |
| | L24 | 0.169 | - |
| **SS in protomer E** | | | |
| | S6 | - | 0.992 |
| | L11 | - | 0.992 |
| | H11 | - | 0.259 |
| | L12 | - | 0.340 |
| | H12 | 0.278 | 0.267 |
| | H13 | - | 0.364 |
| PL3 | H17 | - | 0.297 |
| | H18 | - | 0.265 |
| | L20 | - | 0.771 |
| | L24 | - | 0.258 |

**E**

| Notes | SS | APO | CPX |
|---|---|---|---|
| **SS in protomer E** | | | |
| | L2 | - | 1.000 |
| | H2 | - | 0.933 |
| | H10 | - | 0.405 |
| | S7 | - | 0.579 |
| | H11 | - | 0.205 |
| | L13 | - | 0.426 |
| | H13 | - | 0.475 |
| | H15 | 0.738 | - |
| | L16 | 0.436 | - |
| | H18 | 0.359 | 0.380 |
| | L20 | 0.266 | 1.000 |
| | S10 | 0.118 | - |
| | L24 | 0.233 | - |
| | S13 | 0.539 | - |
| **SS in protomer F** | | | |
| | S6 | - | 0.997 |
| | L11 | 0.715 | 0.300 |
| | H11 | 1.000 | - |
| | L12 | - | 0.455 |
| | H12 | - | 0.378 |
| | L14 | 0.865 | - |
| | H14 | 0.366 | - |
| | L15 | 0.354 | - |
| | L17 | 0.323 | - |
| WA2 | H16 | 0.389 | - |
| | L28 | 0.323 | - |
| | H23 | 0.342 | - |
| | L30 | 0.634 | - |
| | H25 | 0.483 | - |

**F**

| Notes | SS | APO | CPX |
|---|---|---|---|
| **SS in protomer A** | | | |
| | L4 | 0.445 | - |
| | L12 | 0.618 | 1.000 |
| | H12 | 0.435 | 0.596 |
| **SS in protomer F** | | | |
| | H2 | 0.481 | 0.962 |
| | L3 | - | 0.160 |
| Sink | S2 | - | 0.489 |
| | S6 | - | 0.750 |
| | L11 | 0.717 | 0.392 |
| | H10 | - | 0.687 |
| | H11 | 1.000 | 0.290 |
| | H13 | - | 0.378 |
| | L14 | 0.908 | 1.000 |
| | H14 | 0.500 | - |
| WA2 | H16 | 0.986 | 0.658 |
| | S9 | - | 0.177 |
| | H18 | 0.239 | 0.821 |
| | L20 | 0.482 | 0.156 |
| | S10 | - | 0.289 |

Table S37. Average node degeneracy of secondary structures comprising inter-protomer PL3-WA1 allosteric communication networks in hexameric ClpB. The six panels show the average node degeneracy for inter-protomer communications of protomer A-F in APO and CPX setups. The protomer origin of secondary structures is listed. A dash (“–”) indicates that the secondary structure did not participate in the network in that setup. Values are shaded on a green gradient from light (degeneracy ≈ 0) to dark (degeneracy ≈ 1). The functional regions and the sink and source regions of the network, that the listed secondary structure consisted of, are noted.

| WA to PL1 | | | | WA to PL2 | | | | PL1 to HBD tip | | | | PL2 to HBD tip | | | |
|---|---|---|---|---|---|---|---|---|---|---|---|---|---|---|---|
| Notes | SS | APO | CPX | Notes | SS | APO | CPX | Notes | SS | APO | CPX | Notes | SS | APO | CPX |
| | H1 | 0.32 | - | | H1 | 0.173 | - | | B1 | 0.469 | - | | B1 | 0.458 | - |
| | L2 | 0.28 | - | | L2 | 0.227 | - | WA | L4 | - | 0.314 | WA | L4 | - | 0.3 |
| | B2 | 0.324 | 0.294 | Source | H3 | - | 0.271 | WA | H3 | - | 0.269 | WA | H3 | - | 0.339 |
| Source | H3 | - | 0.375 | | B2 | 0.247 | 0.174 | | B2 | 0.246 | 0.178 | | B2 | - | 0.145 |
| Sink | H4 | 0.27 | - | PL1 | L6 | 0.168 | 0.404 | Source | L6 | 0.439 | 0.449 | PL1 | L6 | - | 0.398 |
| Sink | L6 | - | 0.405 | PL1 | H4 | 0.17 | - | WB | B3 | 0.289 | 0.656 | WB | B3 | - | 0.655 |
| WB | B3 | 0.879 | 0.662 | WB | B3 | 0.715 | 0.645 | WB | L8 | 0.296 | 0.197 | WB | L8 | 0.357 | 0.268 |
| WB | L8 | - | 0.108 | WB | L8 | 0.145 | 0.181 | | B4 | 0.318 | 0.116 | | H5 | 0.57 | - |
| | | | | | H5 | 0.182 | - | | L13 | 0.999 | - | Source | H6 | 0.996 | 0.373 |
| | | | | Sink | L9 | 0.126 | 0.169 | | L15 | 0.485 | 0.276 | | B4 | 0.468 | 0.212 |
| | | | | Sink | H6 | 0.414 | 0.433 | | H10 | 0.371 | 0.494 | | L13 | 0.977 | - |
| | | | | | B4 | 0.166 | 0.233 | | | | | | L15 | 0.467 | 0.25 |
| | | | | | | | | | | | | | H10 | 0.324 | 0.495 |

Table S38. Average node degeneracy of secondary structures of the katanin monomer. The four panels show the average node degeneracy for the different paths including ATP binding pocket (sources: WA - P235 and T240) to the CTT binding channel (sinks: PL1 - W266 and E271, PL2 - H307 and R312) and from the CTT binding pore (sources: PL1 - W266 and E271, PL2 - H307 and R312) to the HBD tip (sinks: T418 and L426). A dash ("–") indicates that the secondary structure did not participate in the network in that setup. Values are shaded on a green gradient from light (degeneracy ≈ 0) to dark (degeneracy ≈ 1). The functional regions and the sink and source regions of the network, that the listed secondary structure consisted of, are noted.

| A | | | | B | | | | C | | | | D | | | | E | | | | F | | | |
|---|---|---|---|---|---|---|---|---|---|---|---|---|---|---|---|---|---|---|---|---|---|---|---|
| Notes | SS | APO | CPX | Notes | SS | APO | CPX | Notes | SS | APO | CPX | Notes | SS | APO | CPX | Notes | SS | APO | CPX | Notes | SS | APO | CPX |
| SS in protomer A | | | | SS in protomer B | | | | SS in protomer C | | | | SS in protomer D | | | | SS in protomer E | | | | SS in protomer F | | | |
| | B1 | 0.268 | 0.179 | | B1 | 0.150 | 0.171 | | B1 | 0.16 | 0.16 | | B1 | 0.19 | 0.15 | | B1 | 0.10 | - | | B1 | - | 0.14 |
| Source | L4 | 0.250 | 0.244 | Source | L4 | 0.193 | 0.173 | Source | L4 | 0.12 | 0.17 | Source | L4 | 0.13 | 0.22 | Source | L4 | 0.19 | 0.22 | Source | L4 | 0.22 | 0.18 |
| Source | H3 | - | 0.176 | Source | H3 | 0.194 | 0.168 | Source | H3 | 0.16 | 0.25 | Source | H3 | 0.20 | 0.19 | Source | H3 | 0.20 | 0.27 | Source | H3 | 0.31 | 0.20 |
| | B2 | - | 0.126 | | B2 | 0.218 | - | | B2 | - | 0.10 | | L5 | - | 0.14 | | L5 | - | 0.16 | | L5 | 0.11 | - |
| | L6 | 0.113 | - | Sink | H4 | 0.280 | 0.315 | Sink | L6 | 0.50 | 0.50 | | B2 | 0.22 | - | | B2 | 0.14 | 0.20 | | B2 | 0.20 | 0.19 |
| Sink | H4 | 0.251 | 0.327 | | L7 | 0.172 | 0.145 | Sink | H4 | 0.30 | 0.26 | Sink | L6 | 0.16 | - | Sink | H4 | 0.25 | 0.30 | Sink | H4 | 0.29 | 0.27 |
| | L7 | - | 0.161 | | B3 | 0.257 | 0.285 | | L7 | 0.26 | 0.38 | Sink | H4 | 0.22 | 0.27 | | L7 | - | 0.81 | | L7 | 0.29 | - |
| | B3 | 0.175 | 0.264 | | L10 | 0.148 | - | | B3 | 0.34 | 0.24 | | L7 | 0.16 | 0.25 | | B3 | 0.21 | - | | B3 | 0.18 | 0.24 |
| | L10 | 0.471 | 0.202 | | B4 | 0.143 | 0.221 | | L10 | 0.11 | 0.132 | | B3 | 0.25 | 0.25 | | L10 | - | 0.17 | | L8 | - | 0.12 |
| | B4 | 0.257 | 0.175 | | B5 | - | 0.122 | | B4 | 0.20 | 0.193 | | L10 | - | 0.11 | | B4 | 0.13 | - | | B4 | - | 0.17 |
| | B5 | 0.120 | 0.123 | | L13 | - | 0.174 | SS in protomer B | | | | | B4 | 0.16 | 0.16 | | | | | | | | |
| | | | | | | | | | L6 | - | 0.500 | | B5 | - | 0.11 | | | | | | | | |

Table S39. Average node degeneracy of secondary structures comprising intra-protomer WA to PL1 allosteric communication networks in hexameric katanin. The six panels show the average node degeneracy for intra-protomer communications of protomer A-F in APO and CPX setups. The protomer origin of secondary structures is listed. A dash ("–") indicates that the secondary structure did not participate in the network in that setup. Values are shaded on a green gradient from light (degeneracy ≈ 0) to dark (degeneracy ≈ 1). The functional regions and the sink and source regions of the network, that the listed secondary structure consisted of, are noted.

| Notes | SS | APO | CPX |
|---|---|---|---|
| **SS in protomer A** | | | |
| | B1 | 0.19 | 0.18 |
| Source | L4 | 0.24 | 0.25 |
| Source | H3 | - | 0.23 |
| | B2 | 0.21 | 0.12 |
| | L6 | 0.88 | 0.48 |
| | H4 | - | 0.37 |
| | L7 | - | 0.14 |
| | B3 | 0.43 | 0.26 |
| | L8 | 0.12 | - |
| | L10 | - | 0.18 |
| | B4 | 0.20 | 0.17 |
| | B5 | - | 0.12 |
| **SS in protomer B** | | | |
| | L6 | - | 1.00 |
| | L9 | 0.98 | - |
| | H6 | 0.35 | - |
| **SS in protomer C** | | | |
| | L6 | 0.38 | 0.99 |
| | H4 | 0.42 | - |
| | L9 | 1.00 | - |
| | H6 | 0.32 | - |
| **SS in protomer D** | | | |
| | L6 | 0.57 | 1.00 |
| | H4 | 0.86 | 0.95 |
| **SS in protomer E** | | | |
| | L6 | 0.98 | 0.50 |
| **SS in protomer F** | | | |
| | L6 | 0.30 | 0.50 |
| Sink | H4 | 0.13 | - |

| Notes | SS | APO | CPX |
|---|---|---|---|
| **SS in protomer A** | | | |
| | L6 | 0.46 | 0.35 |
| | H4 | 0.16 | 0.17 |
| **SS in protomer B** | | | |
| | B1 | 0.14 | 0.17 |
| Source | L4 | 0.20 | 0.18 |
| Source | H3 | 0.19 | 0.18 |
| | B2 | 0.22 | - |
| | L6 | - | 0.16 |
| | H4 | 0.30 | 0.34 |
| | L7 | 0.16 | - |
| | B3 | 0.27 | 0.40 |
| | L10 | 0.15 | - |
| | B4 | 0.16 | 0.36 |
| | B5 | - | 0.11 |
| | L13 | - | 0.17 |

| Notes | SS | APO | CPX |
|---|---|---|---|
| **SS in protomer B** | | | |
| | B1 | - | 0.25 |
| | L4 | - | 0.28 |
| Sink | L6 | - | 0.20 |
| Sink | H4 | 0.37 | 0.24 |
| | B3 | - | 0.68 |
| | L9 | 0.99 | - |
| | H6 | 0.30 | - |
| | B4 | - | 0.59 |
| | L13 | - | 0.19 |
| **SS in protomer C** | | | |
| | B1 | 0.24 | 0.16 |
| Source | L4 | - | 0.28 |
| Source | H3 | 0.21 | 0.19 |
| | L6 | - | 0.21 |
| | H4 | 0.27 | 0.11 |
| | L7 | 0.14 | - |
| | B3 | 0.33 | 0.33 |
| | L9 | 1.00 | - |
| | H6 | 0.29 | - |
| | B4 | 0.19 | 0.17 |
| | H7 | - | 0.32 |
| | B5 | - | 0.32 |

| Notes | SS | APO | CPX |
|---|---|---|---|
| **SS in protomer C** | | | |
| Sink | L6 | 0.90 | 0.59 |
| Sink | H4 | 0.24 | - |
| **SS in protomer D** | | | |
| | B1 | 0.19 | 0.15 |
| Source | L4 | 0.12 | 0.22 |
| Source | H3 | 0.19 | 0.19 |
| | L5 | - | 0.14 |
| | B2 | 0.22 | - |
| | H4 | 0.33 | 0.30 |
| | L7 | 0.20 | 0.25 |
| | B3 | 0.30 | 0.25 |
| | L10 | - | 0.11 |
| | B4 | 0.14 | 0.16 |
| | B5 | - | 0.10 |

| Notes | SS | APO | CPX |
|---|---|---|---|
| **SS in protomer D** | | | |
| Sink | L6 | 0.36 | 0.39 |
| Sink | H4 | 0.18 | - |
| **SS in protomer E** | | | |
| Source | L4 | 0.14 | 0.22 |
| Source | H3 | 0.19 | 0.26 |
| | L5 | - | 0.15 |
| | B2 | 0.13 | 0.20 |
| | L6 | - | 0.37 |
| | H4 | 0.16 | 0.36 |
| | L7 | - | 0.84 |
| | B3 | 0.15 | - |
| | H6 | 0.16 | - |
| | L10 | - | 0.16 |
| | L11 | 0.17 | - |
| | H7 | 0.19 | - |
| | H13 | 0.10 | - |

| Notes | SS | APO | CPX |
|---|---|---|---|
| **SS in protomer E** | | | |
| Sink | L6 | 0.37 | 0.17 |
| Sink | H4 | 0.21 | 0.45 |
| **SS in protomer F** | | | |
| | B1 | 0.11 | - |
| Source | L4 | 0.16 | 0.21 |
| Source | H3 | 0.17 | 0.30 |
| | B2 | 0.20 | 0.20 |
| | L6 | 0.62 | 0.49 |
| | H4 | 0.26 | 0.30 |
| | L7 | - | 0.25 |
| | B3 | 0.30 | 0.18 |
| | B4 | 0.24 | - |

Table S40. Average node degeneracy of secondary structures comprising inter-protomer WA to PL1 allosteric communication networks in hexameric katanin. The six panels show the average node degeneracy for inter-protomer communications of protomer A-F in APO and CPX setups. The protomer origin of secondary structures is listed. A dash ("–") indicates that the secondary structure did not participate in the network in that setup. Values are shaded on a green gradient from light (degeneracy ≈ 0) to dark (degeneracy ≈ 1). The functional regions and the sink and source regions of the network, that the listed secondary structure consisted of, are noted.

| A | | | |
|---|---|---|---|
| Notes | SS | APO | CPX |
| **SS in protomer A** | | | |
| | B1 | 0.38 | 0.36 |
| | B2 | 0.48 | - |
| Source | L6 | 0.32 | - |
| Source | H4 | 0.22 | 0.33 |
| | B3 | 0.20 | 0.72 |
| | L10 | 0.19 | 0.19 |
| | B4 | 0.23 | 0.35 |
| | B5 | - | 0.25 |
| | L13 | 0.66 | 0.65 |
| | H8 | 0.25 | 0.26 |
| | L14 | 0.11 | 0.30 |
| | H9 | - | 0.18 |
| | L15 | 0.20 | 0.13 |
| | H10 | 0.30 | 0.28 |
| Sink | H11 | 0.30 | 0.25 |
| | L17 | 0.21 | 0.32 |

| B | | | |
|---|---|---|---|
| Notes | SS | APO | CPX |
| **SS in protomer B** | | | |
| | B1 | 0.36 | 0.32 |
| | L4 | - | 0.39 |
| Source | H4 | 0.35 | 0.29 |
| | B3 | 0.71 | 0.94 |
| | L10 | 0.24 | |
| | B4 | 0.33 | 0.89 |
| | B5 | 0.28 | 0.11 |
| | L13 | 0.46 | 0.40 |
| | H8 | 0.38 | - |
| | L15 | - | 0.27 |
| | H10 | 0.37 | 0.31 |
| Sink | H11 | 0.26 | 0.25 |
| | L17 | 0.14 | 0.32 |
| | H12 | - | 0.18 |

| C | | | |
|---|---|---|---|
| Notes | SS | APO | CPX |
| **SS in protomer C** | | | |
| | B1 | 0.55 | 0.39 |
| | L4 | - | 0.27 |
| Source | L6 | 0.32 | 0.50 |
| Source | H4 | 0.28 | 0.27 |
| | L7 | - | 0.13 |
| | B3 | 0.46 | 0.53 |
| | L10 | - | 0.14 |
| | B4 | 0.45 | 0.33 |
| | B5 | - | 0.11 |
| | L13 | 0.66 | 0.50 |
| | H8 | 0.36 | 0.21 |
| | L14 | 0.27 | 0.12 |
| | L15 | - | 0.22 |
| | H10 | 0.26 | 0.30 |
| Sink | H11 | 0.20 | 0.23 |
| | L17 | 0.10 | - |
| **SS in protomer B** | | | |
| | L6 | - | 0.500 |

| D | | | |
|---|---|---|---|
| Notes | SS | APO | CPX |
| **SS in protomer D** | | | |
| | B1 | 0.49 | 0.37 |
| | L4 | - | 0.51 |
| | H3 | - | 0.26 |
| | L5 | - | 0.13 |
| | B2 | 0.49 | 0.15 |
| Source | L6 | 0.32 | - |
| Source | H4 | 0.28 | 0.42 |
| | B3 | 0.33 | 0.35 |
| | B4 | 0.34 | 0.37 |
| | L13 | 0.62 | 0.34 |
| | L15 | 0.29 | 0.28 |
| | H10 | 0.28 | 0.29 |
| Sink | H11 | 0.23 | 0.23 |
| | L17 | 0.38 | 0.24 |

| E | | | |
|---|---|---|---|
| Notes | SS | APO | CPX |
| **SS in protomer E** | | | |
| | B1 | 0.50 | - |
| | L4 | - | 0.32 |
| | H3 | - | 0.30 |
| Source | L6 | 0.19 | 0.21 |
| Source | H4 | 0.25 | 0.26 |
| | L7 | - | 0.61 |
| | B3 | 0.39 | 0.24 |
| | L8 | 0.20 | - |
| | B4 | 0.44 | - |
| | L13 | 0.97 | - |
| | H8 | 0.14 | - |
| | L15 | 0.40 | 0.55 |
| | H10 | 0.30 | 0.33 |
| Sink | H11 | 0.26 | 0.27 |
| | L17 | 0.19 | 0.17 |

| F | | | |
|---|---|---|---|
| Notes | SS | APO | CPX |
| **SS in protomer F** | | | |
| | B1 | 0.46 | - |
| Source | L4 | - | 0.37 |
| Source | H3 | - | 0.30 |
| | B2 | 0.29 | 0.18 |
| | L6 | 0.49 | 0.23 |
| Sink | H4 | 0.25 | 0.46 |
| | B3 | 0.24 | 0.25 |
| | L8 | 0.24 | 0.34 |
| | B4 | 0.30 | - |
| | L13 | 0.86 | - |
| | H8 | 0.42 | - |
| | L15 | 0.14 | 0.26 |
| | H10 | 0.29 | 0.32 |
| | H11 | 0.31 | 0.29 |

Table S41. Average node degeneracy of secondary structures comprising intra-protomer PL1 to HBD tip allosteric communication networks in hexameric katanin. The six panels show the average node degeneracy for intra-protomer communications of protomer A-F in APO and CPX setups. The protomer origin of secondary structures is listed. A dash ("–") indicates that the secondary structure did not participate in the network in that setup. Values are shaded on a green gradient from light (degeneracy ≈ 0) to dark (degeneracy ≈ 1). The functional regions and the sink and source regions of the network, that the listed secondary structure consisted of, are noted.

| Notes | SS | APO | CPX |
|---|---|---|---|
| **SS in protomer A** | | | |
| Source | L6 | 0.50 | 0.36 |
| Source | H4 | 0.35 | - |
| **SS in protomer B** | | | |
| | L6 | - | 1.00 |
| | H4 | 0.39 | - |
| | L9 | 1.00 | - |
| | H6 | 0.49 | - |
| **SS in protomer C** | | | |
| | L6 | 0.99 | 1.00 |
| | H4 | 0.94 | - |
| | L9 | 1.00 | - |
| | H6 | 0.88 | - |
| **SS in protomer D** | | | |
| | L6 | 1.00 | 1.00 |
| | H4 | 0.99 | 1.00 |
| **SS in protomer E** | | | |
| | L6 | 1.00 | 1.00 |
| **SS in protomer F** | | | |
| | B1 | 0.99 | - |
| | L4 | - | 0.38 |
| | H3 | - | 0.37 |
| | B2 | - | 0.38 |
| | L6 | - | 1.00 |
| | H4 | 0.98 | 0.58 |
| | B3 | 0.92 | 0.34 |
| | B4 | 0.92 | - |
| | L13 | 0.96 | - |
| | H8 | 0.45 | - |
| | L15 | 0.19 | 0.27 |
| | H10 | 0.32 | 0.31 |
| Sink | H11 | 0.32 | 0.30 |

| Notes | SS | APO | CPX |
|---|---|---|---|
| **SS in protomer A** | | | |
| | B1 | 0.35 | 0.46 |
| | B2 | - | 0.56 |
| | L6 | 0.93 | 0.50 |
| | H4 | - | 0.21 |
| | B3 | 0.46 | 0.29 |
| | B4 | 0.33 | 0.43 |
| | B5 | - | 0.23 |
| | L13 | 0.66 | 0.63 |
| | H8 | 0.26 | 0.26 |
| | L14 | 0.11 | 0.29 |
| | H9 | - | 0.16 |
| | L15 | 0.18 | 0.13 |
| | H10 | 0.30 | 0.27 |
| Sink | H11 | 0.29 | 0.25 |
| | L17 | 0.20 | 0.30 |
| **SS in protomer B** | | | |
| Source | H4 | 0.41 | - |

| Notes | SS | APO | CPX |
|---|---|---|---|
| **SS in protomer B** | | | |
| | B1 | - | 0.18 |
| | L4 | - | 0.42 |
| | L6 | - | 0.50 |
| | H4 | - | 0.23 |
| | B3 | - | 0.46 |
| | L9 | 1.00 | - |
| | H6 | 0.49 | - |
| | B4 | - | 0.45 |
| | H7 | 0.31 | - |
| | L12 | 0.33 | - |
| | B5 | 0.45 | - |
| | L13 | 0.42 | 0.33 |
| | H8 | 0.39 | - |
| | L15 | - | 0.21 |
| | H10 | 0.37 | 0.31 |
| Sink | H11 | 0.26 | 0.25 |
| | L17 | 0.14 | 0.30 |
| | H12 | - | 0.16 |
| **SS in protomer C** | | | |
| | B1 | - | 0.38 |
| Source | L6 | 0.50 | - |
| Source | H4 | 0.47 | 0.21 |
| | L9 | 1.00 | - |
| | H6 | 0.48 | - |
| | L10 | - | 0.15 |
| | B4 | - | 0.17 |
| | H7 | - | 0.21 |

| Notes | SS | APO | CPX |
|---|---|---|---|
| **SS in protomer C** | | | |
| | B1 | 0.86 | 0.40 |
| | L4 | - | 0.27 |
| | L6 | 1.00 | 0.99 |
| | H4 | - | 0.31 |
| | B3 | 0.49 | 0.61 |
| | L10 | - | 0.11 |
| | B4 | 0.33 | 0.34 |
| | L13 | 0.65 | 0.50 |
| | H8 | 0.34 | 0.20 |
| | L14 | 0.17 | 0.12 |
| | L15 | - | 0.23 |
| | H10 | 0.25 | 0.32 |
| Sink | H11 | 0.18 | 0.23 |
| | L17 | 0.12 | - |
| **SS in protomer D** | | | |
| Source | L6 | - | 0.18 |
| Source | H4 | 0.33 | 0.21 |

| Notes | SS | APO | CPX |
|---|---|---|---|
| **SS in protomer D** | | | |
| | B1 | 0.49 | - |
| | B2 | 0.67 | - |
| | L6 | 0.44 | - |
| | B3 | 0.49 | - |
| | B4 | 0.34 | - |
| | L13 | 0.62 | - |
| | L15 | 0.30 | - |
| | H10 | 0.28 | 0.27 |
| Sink | H11 | 0.24 | 0.22 |
| | L17 | 0.40 | 0.21 |
| | H12 | - | 0.25 |
| | L18 | - | 1.00 |
| **SS in protomer E** | | | |
| | B1 | - | 0.47 |
| Source | L6 | 0.44 | - |
| Source | H4 | - | 0.33 |
| | L10 | - | 0.95 |
| | B4 | - | 0.95 |
| | B5 | - | 0.99 |
| | H13 | - | 0.99 |

| Notes | SS | APO | CPX |
|---|---|---|---|
| **SS in protomer E** | | | |
| | B1 | 0.47 | - |
| | L4 | - | 0.34 |
| | L8 | 0.36 | 0.33 |
| | H5 | 0.32 | 0.83 |
| | B4 | 0.50 | 0.78 |
| | L11 | - | 0.12 |
| | L13 | 0.88 | - |
| | H8 | 0.18 | - |
| | L15 | 0.36 | 0.33 |
| | H10 | 0.29 | 0.30 |
| Sink | H11 | 0.25 | 0.23 |
| | L17 | 0.18 | 0.17 |
| | H12 | - | 0.11 |
| **SS in protomer F** | | | |
| Source | H4 | 0.50 | 0.42 |
| | H6 | 0.94 | 1.00 |

Table S42. Average node degeneracy of secondary structures comprising inter-protomer PL1 to HBD tip allosteric communication networks in hexameric katanin. The six panels show the average node degeneracy for inter-protomer communications of protomer A-F in APO and CPX setups. The protomer origin of secondary structures is listed. A dash (“–”) indicates that the secondary structure did not participate in the network in that setup. Values are shaded on a green gradient from light (degeneracy ≈ 0) to dark (degeneracy ≈ 1). The functional regions and the sink and source regions of the network, that the listed secondary structure consisted of, are noted.

| A | | | | B | | | | C | | | | D | | | | E | | | | F | | | |
|---|---|---|---|---|---|---|---|---|---|---|---|---|---|---|---|---|---|---|---|---|---|---|---|
| Notes | SS | APO | CPX | Notes | SS | APO | CPX | Notes | SS | APO | CPX | Notes | SS | APO | CPX | Notes | SS | APO | CPX | Notes | SS | APO | CPX |
| SS in protomer A | | | | SS in protomer B | | | | SS in protomer C | | | | SS in protomer D | | | | SS in protomer E | | | | SS in protomer F | | | |
| | L3 | - | - | | B1 | 0.11 | 0.18 | | B1 | 0.18 | 0.24 | | B1 | 0.18 | 0.15 | | B1 | 0.10 | - | | B1 | 0.12 | - |
| | B1 | 0.30 | 0.21 | Source | L4 | 0.12 | 0.15 | Source | L4 | 0.12 | 0.19 | Source | L4 | 0.12 | 0.21 | Source | L4 | 0.14 | 0.22 | Source | L4 | 0.17 | 0.22 |
| Source | L4 | 0.25 | 0.25 | Source | H3 | 0.22 | 0.18 | Source | H3 | 0.20 | 0.21 | Source | H3 | 0.19 | 0.20 | Source | H3 | 0.15 | 0.28 | Source | H3 | 0.17 | 0.30 |
| Source | H3 | - | 0.20 | | B2 | 0.22 | - | | H4 | 0.31 | 0.16 | | L5 | - | 0.12 | | L5 | - | 0.16 | | B2 | 0.20 | 0.31 |
| | H4 | - | 0.20 | | H4 | 0.17 | 0.23 | | L7 | 0.21 | 0.11 | | B2 | 0.20 | - | | B2 | - | 0.20 | | H4 | 0.33 | 0.41 |
| | B3 | - | 0.12 | | B3 | 0.13 | 0.22 | | B3 | 0.33 | 0.21 | | H4 | 0.25 | 0.19 | | H4 | - | 0.31 | | L7 | - | 0.26 |
| Sink | H6 | 0.28 | 0.25 | Sink | H6 | 0.24 | 0.24 | Sink | H6 | 0.21 | 0.28 | | L7 | 0.17 | 0.19 | | L7 | - | 0.81 | | B3 | 0.24 | 0.18 |
| | B4 | 0.17 | 0.21 | | B4 | 0.12 | 0.15 | | L10 | 0.10 | - | | B3 | 0.27 | 0.18 | | B3 | 0.15 | - | | L8 | 0.11 | - |
| | L11 | 0.11 | - | | L11 | 0.13 | - | | B4 | 0.20 | 0.22 | Sink | H6 | 0.28 | 0.25 | Sink | H6 | 0.19 | 0.20 | | H5 | 0.16 | - |
| | H7 | 0.39 | 0.27 | | H7 | 0.15 | 0.40 | | L11 | - | 0.11 | | B4 | 0.15 | - | | L10 | - | 0.19 | Sink | H6 | 0.20 | 0.27 |
| | L12 | 0.16 | - | | L12 | - | 0.26 | | H7 | - | 0.37 | | H7 | - | 0.27 | | L11 | 0.18 | - | | B4 | 0.18 | - |
| | B5 | 0.17 | 0.17 | | B5 | - | 0.16 | | L12 | - | 0.15 | | L12 | - | 0.16 | | H7 | 0.20 | - | | | | |
| | L13 | - | 0.13 | | L13 | - | 0.22 | | B5 | - | 0.14 | | B5 | - | 0.13 | | H13 | 0.12 | - | | | | |
| | | | | | | | | | | | | | L13 | - | - | | | | | | | | |

Table S43. Average node degeneracy of secondary structures comprising intra-protomer WA to PL2 allosteric communication networks in hexameric katanin. The six panels show the average node degeneracy for intra-protomer communications of protomer A-F in APO and CPX setups. The protomer origin of secondary structures is listed. A dash (“–”) indicates that the secondary structure did not participate in the network in that setup. Values are shaded on a green gradient from light (degeneracy ≈ 0) to dark (degeneracy ≈ 1). The functional regions and the sink and source regions of the network, that the listed secondary structure consisted of, are noted.

| Notes | SS | APO | CPX |
|---|---|---|---|
| **SS in protomer A** | | | |
| | B1 | 0.18 | 0.18 |
| Source | L4 | 0.23 | 0.25 |
| Source | H3 | - | 0.23 |
| | B2 | 0.17 | 0.12 |
| | L6 | 0.90 | 0.48 |
| | H4 | - | 0.37 |
| | L7 | - | 0.13 |
| | B3 | 0.44 | 0.26 |
| | L8 | 0.11 | - |
| | L10 | - | 0.18 |
| | B4 | 0.18 | 0.17 |
| | B5 | - | 0.14 |
| **SS in protomer B** | | | |
| | L6 | - | 1.00 |
| | L9 | 0.99 | - |
| | H6 | 0.35 | - |
| **SS in protomer C** | | | |
| | L6 | 0.93 | 0.99 |
| | H4 | 0.41 | - |
| | L9 | 1.00 | - |
| | H6 | 0.31 | - |
| **SS in protomer D** | | | |
| | L6 | 0.30 | 1.00 |
| | H4 | 0.35 | 0.95 |
| **SS in protomer E** | | | |
| | L6 | 0.57 | 1.00 |
| | H4 | 0.20 | - |
| **SS in protomer F** | | | |
| | L6 | 0.20 | - |
| Sink | H6 | 0.25 | 0.48 |

| Notes | SS | APO | CPX |
|---|---|---|---|
| **SS in protomer A** | | | |
| | L6 | 0.55 | 0.90 |
| | H4 | 0.34 | 0.37 |
| Sink | H6 | 0.29 | 0.22 |
| **SS in protomer B** | | | |
| | B1 | 0.16 | 0.17 |
| Source | L4 | 0.23 | 0.19 |
| Source | H3 | 0.24 | 0.21 |
| | B2 | 0.24 | - |
| | H4 | 0.35 | 0.36 |
| | L7 | 0.48 | - |
| | B3 | 0.32 | 0.44 |
| | L10 | 0.13 | - |
| | B4 | 0.16 | 0.41 |
| | L13 | - | 0.16 |

| Notes | SS | APO | CPX |
|---|---|---|---|
| **SS in protomer B** | | | |
| | B1 | - | 0.23 |
| | L4 | - | 0.38 |
| Sink | L9 | 0.50 | - |
| Sink | H6 | 0.14 | 0.48 |
| | H7 | - | 1.00 |
| | L12 | - | 0.86 |
| | B5 | - | 0.38 |
| | L13 | - | 0.33 |
| **SS in protomer C** | | | |
| | B1 | 0.26 | 0.19 |
| Source | L4 | 0.11 | 0.22 |
| Source | H3 | 0.20 | 0.19 |
| | B2 | | - |
| | L6 | | - |
| | H4 | 0.29 | - |
| | L7 | 0.18 | - |
| | B3 | 0.32 | 0.37 |
| | L9 | 1.00 | - |
| | H6 | 0.26 | - |
| | B4 | 0.19 | 0.22 |
| | H7 | - | 0.34 |
| | L12 | - | 0.12 |
| | B5 | - | 0.27 |

| Notes | SS | APO | CPX |
|---|---|---|---|
| **SS in protomer C** | | | |
| | L6 | 0.91 | 0.99 |
| | H4 | 0.46 | 0.32 |
| Sink | H6 | 0.18 | 0.39 |
| **SS in protomer D** | | | |
| | B1 | 0.24 | 0.15 |
| Source | L4 | 0.11 | 0.23 |
| Source | H3 | 0.22 | 0.21 |
| | L5 | - | 0.13 |
| | B2 | 0.20 | - |
| | H4 | 0.37 | 0.32 |
| | L7 | 0.14 | 0.21 |
| | B3 | 0.41 | 0.26 |
| | B4 | 0.17 | 0.16 |

| Notes | SS | APO | CPX |
|---|---|---|---|
| **SS in protomer D** | | | |
| | L6 | 0.59 | 0.81 |
| | H4 | - | 0.83 |
| Sink | H6 | 0.53 | 0.19 |
| | H7 | - | 0.16 |
| | L13 | - | 0.14 |
| | L18 | - | 0.11 |
| **SS in protomer E** | | | |
| Source | L4 | 0.14 | 0.21 |
| Source | H3 | 0.18 | 0.25 |
| | L5 | - | 0.14 |
| | B2 | 0.12 | 0.17 |
| | L6 | 0.16 | - |
| | H4 | 0.17 | 0.32 |
| | L7 | - | 0.70 |
| | B3 | 0.13 | - |
| | H6 | 0.16 | - |
| | L10 | - | 0.13 |
| | L11 | 0.17 | - |
| | H7 | 0.17 | - |
| | H13 | 0.11 | 0.13 |

| Notes | SS | APO | CPX |
|---|---|---|---|
| **SS in protomer E** | | | |
| | L6 | 0.79 | 0.47 |
| | H4 | 0.20 | 0.12 |
| Sink | H6 | 0.32 | 0.48 |
| **SS in protomer F** | | | |
| Source | L4 | 0.13 | 0.21 |
| Source | H3 | 0.17 | 0.29 |
| | B2 | 0.19 | 0.16 |
| | L6 | 0.35 | 0.49 |
| | H4 | 0.39 | 0.29 |
| | L7 | - | 0.22 |
| | B3 | 0.29 | 0.19 |
| | B4 | 0.25 | - |

Table S44. Average node degeneracy of secondary structures comprising inter-protomer WA to PL2 allosteric communication networks in hexameric katanin. The six panels show the average node degeneracy for intra-protomer communications of protomer A-F in APO and CPX setups. The protomer origin of secondary structures is listed. A dash ("–") indicates that the secondary structure did not participate in the network in that setup. Values are shaded on a green gradient from light (degeneracy ≈ 0) to dark (degeneracy ≈ 1). The functional regions and the sink and source regions of the network, that the listed secondary structure consisted of, are noted.

| A | | | |
|---|---|---|---|
| Notes | SS | APO | CPX |
| SS in protomer A | | | |
| | B1 | 0.40 | 0.39 |
| Source | H6 | 0.32 | 0.59 |
| | H7 | 0.91 | 0.97 |
| | L12 | 0.12 | 0.18 |
| | B5 | 0.25 | 0.38 |
| | L13 | 0.64 | 0.48 |
| | H8 | 0.25 | 0.26 |
| | L14 | 0.11 | 0.28 |
| | H9 | - | 0.16 |
| | L15 | 0.19 | 0.14 |
| | H10 | 0.30 | 0.27 |
| Sink | H11 | 0.30 | 0.25 |
| | L17 | 0.20 | 0.30 |

| B | | | |
|---|---|---|---|
| Notes | SS | APO | CPX |
| SS in protomer B | | | |
| | B1 | - | 0.19 |
| | L4 | - | 0.32 |
| Source | H6 | 0.38 | 0.49 |
| | H7 | 0.30 | 1.00 |
| | L12 | 0.35 | 0.99 |
| | B5 | 0.44 | 0.45 |
| | L13 | 0.41 | 0.42 |
| | H8 | 0.38 | - |
| | H10 | 0.36 | 0.27 |
| Sink | H11 | 0.26 | 0.24 |
| | L17 | 0.15 | 0.26 |
| | H12 | | 0.14 |

| C | | | |
|---|---|---|---|
| Notes | SS | APO | CPX |
| SS in protomer C | | | |
| | B1 | 0.59 | 0.27 |
| | L4 | - | 0.28 |
| | H4 | 0.40 | - |
| | B3 | 0.83 | - |
| Source | H6 | 0.28 | 0.37 |
| | B4 | 0.48 | - |
| | H7 | - | 0.83 |
| | L12 | - | 0.31 |
| | B5 | - | 0.29 |
| | L13 | 0.66 | 0.49 |
| | H8 | 0.35 | 0.21 |
| | L14 | 0.29 | 0.12 |
| | L15 | - | 0.23 |
| | H10 | 0.27 | 0.27 |
| Sink | H11 | 0.20 | 0.23 |

| D | | | |
|---|---|---|---|
| Notes | SS | APO | CPX |
| SS in protomer D | | | |
| | B1 | 0.49 | 0.33 |
| | L4 | - | 0.22 |
| | H4 | 0.15 | - |
| | B3 | 0.20 | - |
| | H5 | 0.53 | - |
| Source | H6 | 0.36 | 0.73 |
| | B4 | 0.31 | - |
| | H7 | - | 0.99 |
| | L12 | - | 0.52 |
| | B5 | - | 0.49 |
| | L13 | 0.47 | 0.65 |
| | L15 | 0.29 | 0.40 |
| | H10 | 0.28 | 0.29 |
| Sink | H11 | 0.23 | 0.22 |
| | L17 | 0.38 | 0.25 |

| E | | | |
|---|---|---|---|
| Notes | SS | APO | CPX |
| SS in protomer E | | | |
| | B1 | 0.58 | - |
| | L4 | - | 0.32 |
| | H3 | - | 0.34 |
| | H4 | - | 0.44 |
| | L7 | - | 0.90 |
| Source | H6 | 0.49 | 0.22 |
| | L11 | 0.36 | - |
| | H7 | 0.20 | - |
| | B5 | 0.15 | - |
| | L13 | 0.54 | - |
| | H8 | 0.14 | - |
| | L15 | 0.39 | 0.59 |
| | H10 | 0.30 | 0.36 |
| Sink | H11 | 0.26 | 0.28 |
| | L17 | 0.20 | 0.15 |
| | H12 | 0.15 | - |

| F | | | |
|---|---|---|---|
| Notes | SS | APO | CPX |
| SS in protomer F | | | |
| | B1 | 0.34 | - |
| | L4 | 0.17 | 0.41 |
| | H3 | - | 0.31 |
| | B2 | - | 0.25 |
| | H4 | - | 0.49 |
| | B3 | - | 0.35 |
| | H5 | 0.99 | - |
| Source | H6 | 0.53 | 0.40 |
| | B4 | 0.39 | - |
| | L13 | 0.64 | - |
| | H8 | 0.35 | - |
| | L15 | 0.21 | 0.27 |
| | H10 | 0.30 | 0.31 |
| Sink | H11 | 0.31 | 0.30 |

Table S45. Average node degeneracy of secondary structures comprising intra-protomer PL2 to HBD tip allosteric communication networks in hexameric katanin. The six panels show the average node degeneracy for intra-protomer communications of protomer A-F in APO and CPX setups. The protomer origin of secondary structures is listed. A dash ("–") indicates that the secondary structure did not participate in the network in that setup. Values are shaded on a green gradient from light (degeneracy ≈ 0) to dark (degeneracy ≈ 1). The functional regions and the sink and source regions of the network, that the listed secondary structure consisted of, are noted.

| Notes | SS | APO | CPX |
|---|---|---|---|
| **SS in protomer A** | | | |
| | L6 | 0.96 | 0.32 |
| | H4 | 0.36 | 0.37 |
| Source | H6 | 0.46 | 0.22 |
| **SS in protomer B** | | | |
| | L6 | - | 1.00 |
| | H4 | 0.96 | - |
| | L9 | 1.00 | |
| | H6 | 0.65 | - |
| **SS in protomer C** | | | |
| | L6 | 0.99 | 1.00 |
| | H4 | 0.95 | - |
| | L9 | 1.00 | - |
| | H6 | 0.90 | - |
| **SS in protomer D** | | | |
| | L6 | 1.00 | 1.00 |
| | H4 | 0.99 | 1.00 |
| **SS in protomer E** | | | |
| | L6 | 1.00 | 1.00 |
| **SS in protomer F** | | | |
| | B1 | 0.99 | - |
| | L4 | - | 0.43 |
| | H3 | - | 0.48 |
| | B2 | - | 0.40 |
| | L6 | - | 1.00 |
| | H4 | 0.99 | 0.59 |
| | B3 | 0.94 | 0.34 |
| | B4 | 0.94 | - |
| | L13 | 0.96 | - |
| | H8 | 0.46 | |
| | L15 | 0.19 | 0.27 |
| | H10 | 0.32 | 0.31 |
| Sink | H11 | 0.32 | 0.25 |

| Notes | SS | APO | CPX |
|---|---|---|---|
| **SS in protomer A** | | | |
| | B1 | 0.48 | 0.37 |
| | L6 | 0.99 | 0.99 |
| | B3 | 0.46 | 0.96 |
| | B4 | 0.33 | 0.45 |
| | B5 | - | 0.21 |
| | L13 | 0.66 | 0.61 |
| | H8 | 0.26 | 0.22 |
| | L14 | 0.11 | 0.27 |
| | H9 | - | 0.15 |
| | L15 | 0.18 | 0.13 |
| | H10 | 0.30 | 0.25 |
| Sink | H11 | 0.30 | 0.24 |
| | L17 | 0.20 | 0.28 |
| **SS in protomer B** | | | |
| Source | H6 | 0.26 | 0.50 |

| Notes | SS | APO | CPX |
|---|---|---|---|
| **SS in protomer B** | | | |
| | L4 | - | 0.95 |
| | L9 | 0.50 | - |
| | H6 (sou | 0.48 | - |
| | H7 | 0.31 | - |
| | L12 | 0.32 | - |
| | B5 | 0.44 | - |
| | L13 | 0.41 | - |
| | H8 | 0.38 | - |
| | L15 | - | 0.24 |
| | H10 | 0.36 | 0.29 |
| Sink | H11 | 0.26 | 0.22 |
| | L17 | 0.15 | 0.17 |
| | H12 | - | 0.13 |
| **SS in protomer C** | | | |
| | L9 | 0.50 | - |
| Source | H6 | - | 0.48 |
| | H7 | - | 1.00 |

| Notes | SS | APO | CPX |
|---|---|---|---|
| **SS in protomer C** | | | |
| | B1 | 0.88 | 0.39 |
| | L4 | - | 0.27 |
| | L6 | 1.00 | 0.99 |
| | H4 | - | 0.31 |
| | L7 | - | 0.10 |
| | B3 | 0.49 | 0.58 |
| | L10 | - | 0.11 |
| | B4 | 0.33 | 0.34 |
| | L13 | 0.66 | 0.50 |
| | H8 | 0.34 | 0.20 |
| | L14 | 0.17 | 0.12 |
| | L15 | - | 0.23 |
| | H10 | 0.25 | 0.31 |
| Sink | H11 | 0.20 | 0.23 |
| | L17 | 0.12 | - |
| **SS in protomer D** | | | |
| | H4 | - | 1.00 |
| Source | H6 | 0.27 | 0.50 |

| Notes | SS | APO | CPX |
|---|---|---|---|
| **SS in protomer D** | | | |
| | B1 | 0.49 | - |
| | B2 | 0.59 | - |
| | L6 | 0.70 | - |
| | B3 | 0.49 | - |
| | B4 | 0.34 | - |
| | L13 | 0.63 | - |
| | L15 | 0.30 | - |
| | H10 | 0.28 | 0.27 |
| Sink | H11 | 0.24 | 0.21 |
| | L17 | 0.41 | 0.22 |
| | H12 | - | 0.25 |
| | L18 | - | 1.00 |
| **SS in protomer E** | | | |
| | B1 | - | 0.54 |
| | H4 | - | 0.33 |
| | H5 | - | 0.33 |
| | L9 | - | 0.33 |
| Source | H6 | 0.20 | 0.22 |
| | L10 | - | 0.67 |
| | B4 | - | 0.66 |
| | L11 | - | 0.28 |
| | B5 | - | 0.66 |
| | H1: | - | 0.98 |

| Notes | SS | APO | CPX |
|---|---|---|---|
| **SS in protomer E** | | | |
| | B1 | 0.46 | - |
| | L4 | - | 0.34 |
| | L8 | 0.20 | 0.33 |
| | H5 | 0.40 | 0.83 |
| | B4 | 0.49 | 0.78 |
| | L11 | - | 0.12 |
| | L13 | 0.85 | - |
| | H8 | 0.19 | |
| | L15 | 0.34 | 0.33 |
| | H10 | 0.29 | 0.30 |
| Sink | H11 | 0.22 | 0.23 |
| | L17 | 0.18 | 0.17 |
| | H12 | - | 0.11 |
| **SS in protomer F** | | | |
| Source | H6 | 0.49 | 0.50 |

Table S46. Average node degeneracy of secondary structures comprising inter-protomer PL2 to HBD tip allosteric communication networks in hexameric katanin. The six panels show the average node degeneracy for inter-protomer communications of protomer A-F in APO and CPX setups. The protomer origin of secondary structures is listed. A dash (“–”) indicates that the secondary structure did not participate in the network in that setup. Values are shaded on a green gradient from light (degeneracy ≈ 0) to dark (degeneracy ≈ 1). The functional regions and the sink and source regions of the network, that the listed secondary structure consisted of, are noted.

### References
